# Inference of single cell profiles from histology stains with the Single Cell omics from Histology Analysis Framework (SCHAF)

**DOI:** 10.1101/2023.03.21.533680

**Authors:** Charles Comiter, Xingjian Chen, Eeshit Dhaval Vaishnav, Koseki J. Kobayashi-Kirschvink, Metamia Ciampricotti, Ke Zhang, Jason Murray, Francesco Monticolo, Jianhuan Qi, Ryota Tanaka, Sonia E. Brodowska, Bo Li, Yiming Yang, Scott J. Rodig, Angeliki Karatza, Alvaro Quintanal Villalonga, Madison Turner, Kathleen L. Pfaff, Judit Jané-Valbuena, Michal Slyper, Julia Waldman, Sebastian Vigneau, Jingyi Wu, Timothy R. Blosser, Åsa Segerstolpe, Daniel L. Abravanel, Nikhil Wagle, Shadmehr Demehri, Xiaowei Zhuang, Charles M. Rudin, Johanna Klughammer, Orit Rozenblatt-Rosen, Collin M. Stultz, Jian Shu, Aviv Regev

**Affiliations:** Klarman Cell Observatory, Broad Institute of MIT and Harvard, Cambridge, MA 02142, USA; Department of Electrical Engineering and Computer Science, MIT, Cambridge, MA 02142, USA; Cutaneous Biology Research Center, Massachusetts General Hospital, Harvard Medical School, Boston, MA 02129, USA; Sequome Inc., 329 Oyster Point Blvd, Suite 300, South San Francisco, CA 94080, USA; Department of Biology, MIT, Cambridge, MA 02140, USA; Druckenmiller Center for Lung Cancer Research, Memorial Sloan Kettering Cancer Center, New York, NY 10065, USA; Department of Pathology, Brigham and Women’s Hospital, Boston, MA 02115, USA; Center for Immuno-Oncology, Dana-Farber Cancer Institute, Boston, MA 02115, USA; Department of Chemistry and Chemical Biology, Harvard University, Cambridge, MA 02138, USA; Howard Hughes Medical Institute; Gene Center and Department of Biochemistry, Ludwig Maximilians Universität München, Munich, Germany; Laser Biomedical Research Center, G. R. Harrison Spectroscopy Laboratory, Massachusetts Institute of Technology, Cambridge, MA 02139, USA; Center for Immunology and Inflammatory Diseases, Department of Medicine, Massachusetts General Hospital, Boston, MA, USA; Department of Medicine, Harvard Medical School, Boston, MA, USA; Department of Medical Oncology, Dana-Farber Cancer Institute, Boston, MA, USA; Department of Microbiology, Immunology, and Cancer Biology, University of Virginia, Charlottesville, VA 22908, USA; Divison of Cardiology, Department of Medicine, Massachusetts General Hospital, Harvard Medical School, Boston, MA 02129, USA; Department of Pathology, Johns Hopkins University School of Medicine, Baltimore, MD 21287, USA; Center for Cancer Immunology, Krantz Family Center for Cancer Research, Massachusetts General Hospital and Harvard Medical School, Boston, MA, USA; Harvard-MIT Program in Health Sciences and Technology, Cambridge, MA 02142, USA; Institute of Medical Engineering and Science, MIT, Cambridge, MA 02142, USA; Perlmutter Cancer Center, New York University Langone Health, New York, NY 10016, USA; Department of Medicine, University of Chicago, Chicago, IL 60637; Genentech, 1 DNA Way, South San Francisco, CA 94080, USA

## Abstract

Tissue biology involves an intricate balance between cell-intrinsic processes and interactions between cells organized in specific spatial patterns, which can be respectively captured by single cell profiling methods, such as single cell RNA-seq (scRNA-seq) and spatial transcriptomics, and histology imaging data, such as Hematoxylin-and-Eosin (H&E) stains. While single cell profiles provide rich molecular information, they can be challenging to collect routinely in the clinic and either lack spatial resolution or high gene throughput. Conversely, histological H&E assays have been a cornerstone of tissue pathology for decades, but do not directly report on molecular details, although the observed structure they capture arises from molecules and cells. Here, we leverage vision transformers and adversarial deep learning to develop the Single Cell omics from Histology Analysis Framework (SCHAF), which generates a tissue sample’s spatially-resolved whole transcriptome single cell omics dataset from its H&E histology image. We demonstrate SCHAF on a variety of tissues— including lung cancer, metastatic breast cancer, placentae, and whole mouse pups—training with matched samples analyzed by sc/snRNA-seq, H&E staining, and, when available, spatial transcriptomics. SCHAF generated appropriate single cell profiles from histology images in test data, related them spatially, and compared well to ground-truth scRNA-Seq, expert pathologist annotations, or direct spatial transcriptomic measurements, with some limitations. SCHAF opens the way to next-generation H&E analyses and an integrated understanding of cell and tissue biology in health and disease.

## INTRODUCTION

Advances in massively parallel, high-resolution molecular profiling now provide cellular and tissue level measurements at a genomic scale^1^. These include methods for massively parallel single cell or single-nucleus (sc/sn) profiling of RNA, chromatin, proteins, and their multi-modal combinations^2^. Simultaneous advancements in data-driven analytics, largely in artificial intelligence / machine learning (AI/ML)^3^, have allowed us to derive biological insights from such rich data^4–8^, in addition to key modalities such as H&E stains, which are used in both research and clinical practice^9^.

However, substantial challenges remain in realizing the promise of these methods in the context of tissue biology, especially for histopathology, a cornerstone of medicine. While the costs and complexity of single cell omics have reduced dramatically^10^, they remain relatively expensive and time consuming and are not yet routinely applied in clinical settings. Experiments also remain prone to technical variations, leading to inter-sample discrepancies and batch effects^11,12^. Moreover, single cell genomics does not directly capture spatial information nor is it directly related to the rich legacy of histology. While spatial transcriptomics methods, such as MERFISH^13^, seqFISH^+14^, Visium^15^, Xenium^16^, ISS^17^, Barista-seq^18^, smFISH^19^, osmFISH^20^, STARmap^21^, Slide-tags^22^, or Targeted ExSeq^23^ measure spatially-resolved expression data, their throughput is still limited, and they involve high per-sample costs and complexities. Computational methods^24–26^ have used limited spatial information measured experimentally to project cell profiles to spatial positions, but they have thus far required some shared variables for such mapping (*i.e.*, transcripts measured both in the dissociated cells and spatially in tissue). In sum, they do not spatially resolve single cell omics data based solely on the microscopic morphology in a histology image.

Recent advances in applied deep learning may open the way to address the challenge of mapping single cell profiles to histology. In other domains, deep learning methods have successfully related data modalities of the same entities even when they do not have nominally shared variables (such as audio and video^27–29)^. Moreover, in recent studies, models were trained to generate a tissue’s bulk RNA-seq or its non-single cell spatial transcriptomics measurements (55 micron diameter), from its histology image^30,31^. Given these successes, we hypothesized that deep learning could also be applied to the more difficult translation problem of inferring single cell expression profiles from histology.

Here, we present the Single Cell omics from Histology Analysis Framework (SCHAF), a deep learning-based framework for translation from histology images to single cell omics data. SCHAF is based on the assumption that a histology image and a single cell omics dataset from the same tissue sample can be explained by a single underlying latent distribution. Given a corpus of tissue data, where each sample has a histology image, sc/snRNA-seq data, and, when available, spatial transcriptomics data, SCHAF discovers this common latent space from these modalities across different samples. SCHAF then leverages this latent space to construct an inference engine mapping a histology image to its corresponding (model-generated) single cell molecular profiles. This inferred dataset has high genomic coverage, yielding expression information on all the genes in the training corpus, and is, in some cases, spatially resolved, with each predicted gene expression profile mapped at single cell resolution. Given the spatial information inherent in its predictions, SCHAF can further predict a spatial portrait of a tissue’s cell types. We demonstrate SCHAF’s performance on these tasks with data from a variety of tissue types, through multiple criteria, including a comparison of SCHAF predictions to experimentally measured spatial transcriptomics data and to expert pathologist annotations of cell types. We further demonstrate that SCHAF outperforms related models for the task of predicting transcriptomic information from H&E images alone.

## RESULTS

### SCHAF: Single Cell omics from Histology Analysis Framework

SCHAF infers single cell omics data from a corresponding histological image. The model is trained on a corpus of samples, each with a matched histology image and single cell transcriptomics data. SCHAF comes in two flavors depending on the type of single cell transcriptomics data available for training (**Fig. 1**).

**Figure 1.**
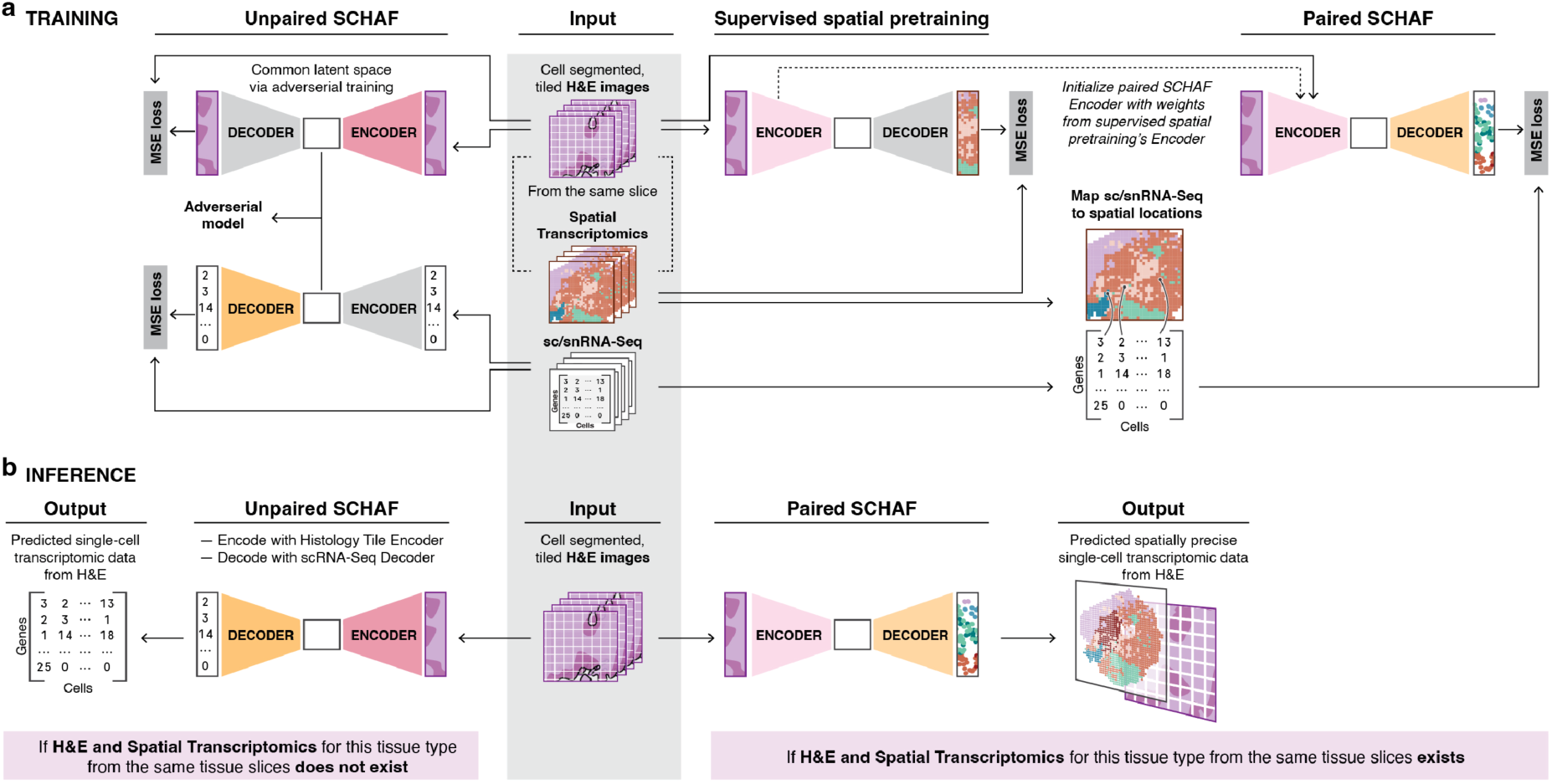
SCHAF learns to predict a tissue’s single cell omics data from its H&E image. **a.** Overview of training. Training data (input) consist of matched cell segmented and tiled histology (H&E) stains, scRNA-seq, and, for Paired SCHAF, spatial transcriptomics from the same section as the H&E from multiple samples. Paired SCHAF (right) is trained in three steps: supervised spatial pretraining of a vision transformer (initialized with weights from a pretrained H&E foundational model) to infer a cell’s spatial transcriptomic profile from its H&E tile; mapping of scRNA-seq onto spatial transcriptomics; and fine-tuning of the encoder of the pre-trained model and a decoder of several new linear layers to infer a cell’s scRNA-seq from its H&E tile. Unpaired SCHAF (left) is trained in two steps. First, a pretrained scRNA-seq transformer encoder, with frozen weights, and a decoder of several linear layers are trained to reconstruct scRNA-seq profiles. Then, a two-part encoder (a pretrained H&E foundational model, with frozen weights, followed by several linear layers) and a decoder, are trained to encode, via training against an adversarial model, to a latent space indistinguishable from the scRNA-seq encoder’s outputs, and to reconstruct the embeddings produced by the pretrained H&E foundational model. **b.** Overview of inference. An input H&E slide is first cell segmented and tiled. In Paired SCHAF, the model from the last part of training is used to generate spatially resolved, genome wide, single cell resolution profiles (right). In Unpaired SCHAF, the H&E tile encoder and the scRNA-seq decoder are used to generate high gene throughput, single cell resolution transcriptomic data with some spatial accuracy guarantees (left).

If spatial transcriptomics data from the same tissue section as the histology image are available in addition to corresponding sc/snRNA-seq, we train Paired SCHAF, through the following steps (**Fig. 1, right, Methods)**. First, Paired SCHAF tiles the histology image into partially overlapping smaller square tiles, each centered around an individual cell, capturing it and its surroundings. Paired SCHAF is then trained in two stages. In the first stage, the model is trained to predict the spatial transcriptomics profile corresponding to each tile (**Fig. 1a, supervised spatial pretraining**), where the initial weights for the image encoder are obtained from a previously developed whole-slide foundational digital pathology model^32^. In the second stage, additional transcriptome-wide expression data from genes not measured in the spatial transcriptomic experiments are mapped onto the spatial locations using Tangram^7^, a computational method that infers spatial locations for scRNA-seq, and Paired SCHAF is updated to predict these additional spatially resolved transcripts (**Fig. 1a, Paired SCHAF**). This second stage improves the model’s ability to infer accurate spatial locations for transcripts that were not captured in the spatial transcriptomics training data. At inference, Paired SCHAF only requires a histology image as input to generate single cell profiles at high (whole transcriptome) gene throughput that are spatially resolved at single cell resolution (as the generated profile is associated with the cell at the center of the tile).

As experimentally determined spatial transcriptomic information is not always available for training, we also develop Unpaired SCHAF, as a method for inferring single cell transcript profiles from histology images that does not rely on spatial transcriptomic training data (**Fig. 1, left, Methods**). Like Paired SCHAF, it first segments the histology image into partially-overlapping tiles, each centered on a cell (**Methods**; cells are segmented by a pretrained AI model for cell segmentation). Unpaired SCHAF then uses encoders for both histopathology and scRNA-seq to build a model that translates from one H&E tile to one scRNA-seq profile. We take the initial weights for each of these encoders from existing foundation models for both histopathology^32^ and scRNA-seq^33^ and use a discriminative network to impose that the latent representations from both encoders be similar, thereby enabling a model that translates from one H&E tile to one scRNA-seq profile. Like Paired SCHAF, the resulting predictions generated at inference have single cell resolution and high gene throughput, albeit with more limited spatial information. When histopathologist annotations are available for training data, they, with scRNA-seq cell type annotations, can also be incorporated into Unpaired SCHAF’s training to encourage the method to infer cell types with some spatial accuracy guarantees. While Paired SCHAF provides spatial transcriptomic information at single cell resolution, Unpaired SCHAF provides information about the distribution of RNA transcripts in a given tissue in addition to spatially resolved cell type information.

The final models (Paired SCHAF and Unpaired SCHAF) are fusions of different encoders and decoders, yielding an end-to-end process to generate a single cell expression profile from a histology image tile (**Fig. 1, bottom, Methods**). At inference, we split the segmented histology image into tiles centered on each cell and generate each cell’s expression profile using the corresponding SCHAF model to infer single cell resolution profiles. In principle, when spatially resolved transcript information was used during training (Paired SCHAF), we expect better performance than when it was not available (Unpaired SCHAF).

### Paired SCHAF infers spatial gene expression at single cell resolution

We developed and tested Paired SCHAF using three tissue corpora, each with H&E, spatial transcriptomics, and sc/snRNA-seq training data available (**Fig. 2a, top, and 2b, left**). **First**, in the one day old **mouse pup**, we found one single tissue slice assayed via H&E and experimentally measured spatial transcriptomics at single cell resolution (ExpSCR) via Xenium, profiling 365 genes over more than 1,000,000 cells^16^, and a second tissue from a different whole one day old mouse pup by scRNA-seq^34^. We partitioned the mouse pup data into four spatial folds and evaluated Paired SCHAF via cross validation, training, for each fold, a model using H&E, scRNA-seq, and ExpSCR data from the remaining three folds, evaluating on the fourth fold, and averaging results over all four folds (**Fig. 2a, top left**). **Second**, in estrogen receptor positive metastatic breast cancer (ER+ MBC), we relied on data we recently collected from one large tumor, analyzed with three adjacent sections^16^: the first was assayed by H&E and ExpSCR/Xenium, the second by scRNA-seq, and the third by Experimentally Measured spatial transcriptomics at low-resolution (ExpLR) via Visium (profiling 17,742 genes, at 55 micron spot resolution), used for validation. We again split the data into four folds, trained models using H&E, scRNA-seq, and ExpSCR data, and evaluated via cross validation and with the ExpLR/Visium data (**Fig. 2a, top right**). **Third**, in estrogen receptor negative metastatic breast cancer (ER-MBC), we used for training a whole single section assayed by H&E and ExpSCR/Xenium^16^. To further evaluate Paired SCHAF’s ability to generalize to new data samples that were not available during training, we trained it using the ER+ MBC data and evaluated its performance on the ER-MBC sample (**Fig. 2b-left**). For each of these three cases, we randomly partitioned the ExpSCR genes into two equally-sized sets: “training genes” (used in supervised spatial pretraining, **Fig. 1a, right**), and “hold-out genes” that are not used during training. We expect performance for training genes to be higher than for hold-out genes, and that the prediction for ER-MBC samples from a model trained only on ER+ MBC to be the most difficult task.

**Figure 2.**
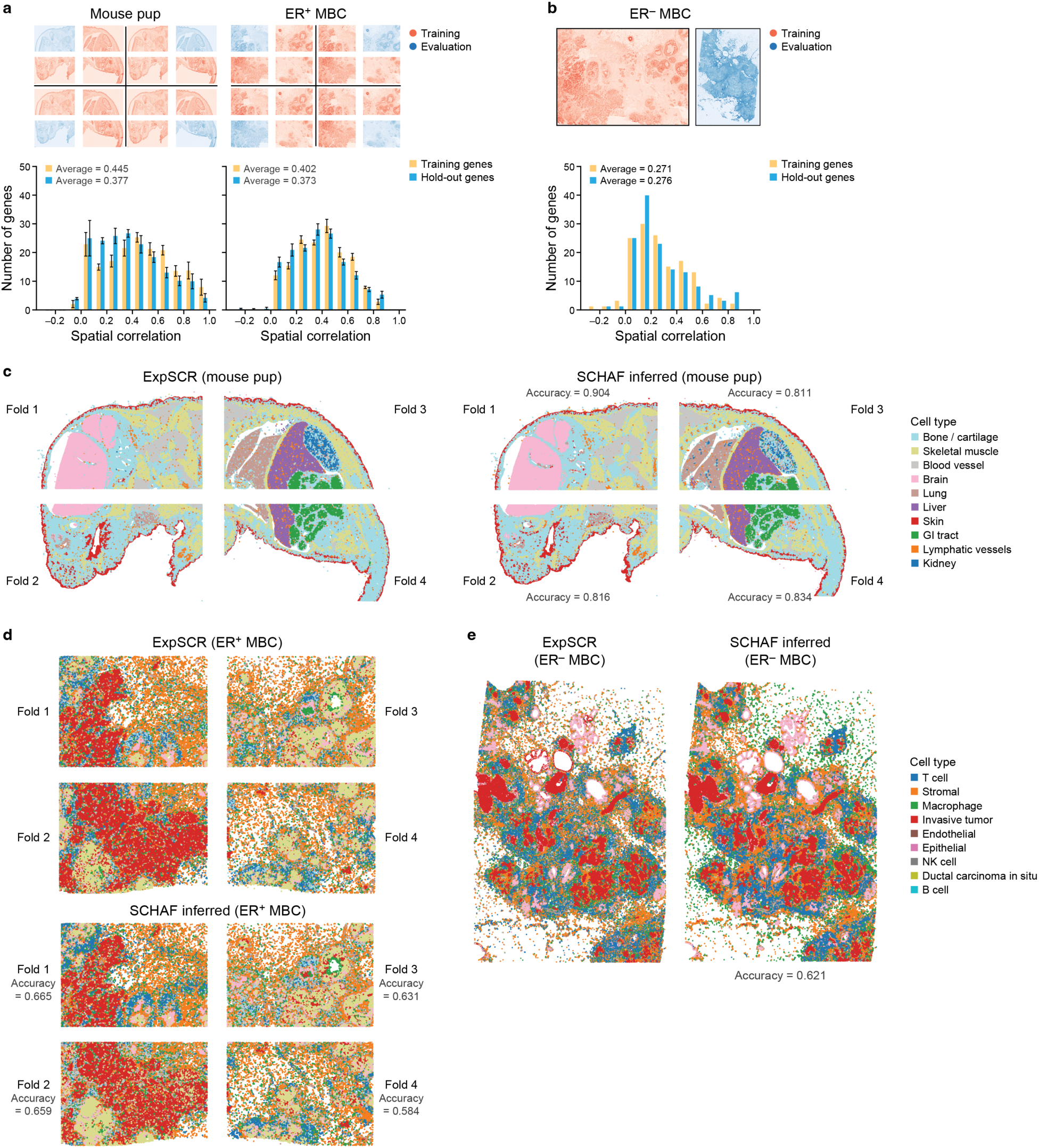
Paired SCHAF inferred profiles accurately recover spatial transcriptomics. **a.** Evaluation by cross validation. Top: A whole mouse pup (left) or ER+ MBC tumor (right) slide is partitioned into four equal-sized folds. Paired SCHAF is trained by 4-fold cross validation with three parts used for training (red) and one held out for evaluation (blue). The results are averaged. Bottom: Distribution of spatial correlations (Pearson’s correlation coefficient of the expression of a gene measured by spatial transcriptomics and predicted by Paired SCHAF) for all spatially-measured genes in training (yellow) and hold-out (blue). Error bars: standard error of folds assessed by cross validation. **b.** Evaluation on an independent tumor. Top: Slidea from an independent ER+ MBC tumor used in training (red) and an ER-MBC tumor used in evaluation (blue). Bottom: Distribution of spatial correlations (Pearson’s r) for all spatially-measured genes in training (yellow) and held out (blue) samples. **c-e.** Paired SCHAF inferred profiles preserve cell types. Spatial cell type annotations (called by a classifier trained on ground truth ExpSCR data, color legend) for measured (ExpSCR) and Paired SCHAF-inferred profiles in the Mouse Pup (**c**), ER+ MBC (**d**), and ER-MBC (**e**) datasets. Accuracy: proportion of cells in each fold assigned to the correct cell type label.

Paired-SCHAF performed well in inferring the spatial expression levels of both training and hold-out genes in four-fold spatial cross validation, based on Pearson’s correlation *vs*. ground truth ExpSCR data (**Fig. 2a, bottom**, mean 0.445 (training), 0.377 (hold-out) for mouse pup; 0.402 (training), 0.373 (hold-out) for ER+ MBC) with 48.7% and 52.1% of genes with strong (>0.4) spatial correlations^7^ for mouse pup and ER+ MBC, respectively. These were reduced, but still positive for predicting ER-MBC from an ER+ MBC model (**Fig. 2b**; mean Pearson’s correlation 0.276 (hold-out) and over 26% genes with high correlations (>0.4)).

Moreover, spatial cell type distributions of Paired SCHAF predictions (using a neural network trained to classify cell type from a single cell profile) preserve spatial patterns accurately (**Fig. 2c-e**), based on the proportion of SCHAF inferred profiles with classifier assigned annotations that were the same as those inferred from the ground truth ExpSCR profiles. Mouse Pup predictions had the highest accuracy across folds (**Fig. 2c**, 0.811-0.904), followed by the ER+ MBC (**Fig. 2d**, 0.584-0.665) and ER-MBC (**Fig. 2e**, 0.621), which still showed strong qualitative agreement. Notably, Paired SCHAF preserves heterogeneity *within* each cell type, such that the expression variation across predicted Paired SCHAF profiles is similar to that across measured ExpSCR (**Supplementary Fig. 1**).

While there are no other extant methods addressing the problem solved by Paired SCHAF, we compared Paired SCHAF’s inferences of training genes to inferences of training genes by four supervised methods (SPIRIT^35^, ST-Net^31^, DeepPT^36^, HE2RNA^30^) that learn transcriptomics from H&E. Paired SCHAF outperformed the other methods in all corpora both in gene level spatial correlation and in cell type accuracy (**Supplementary Fig. 2**).

### A gene-specific quality score identifies genes that Paired SCHAF will accurately predict

While Paired SCHAF infers spatially resolved single cell expression profiles at a higher accuracy than other approaches, not all predicted gene transcripts are spatially accurate. We hypothesized that genes whose predicted expression varies substantially across predicted cells in different spatial positions may be more accurate predictions than those with little spatial variation, which may indicate the models’ inability to capture their signal. Indeed, a gene’s standard deviation over single cell SCHAF predictions agreed well with its actual spatial correlation value for both training and hold-out genes (**Fig. 3a**; r=0.69 (hold-out genes, mouse pup) to 0.99 (training genes, ER+ MBC) including in ER-MBC (0.730 (hold-out) to 0.803 (training)). Based on this observation we defined a gene’s standard deviation over single cell SCHAF predictions as its predictive quality score (PQS). Because this predictive quality score is solely a function of the model *output*, it indicates *a priori* if the model’s prediction is likely to be accurate.

**Figure 3.**
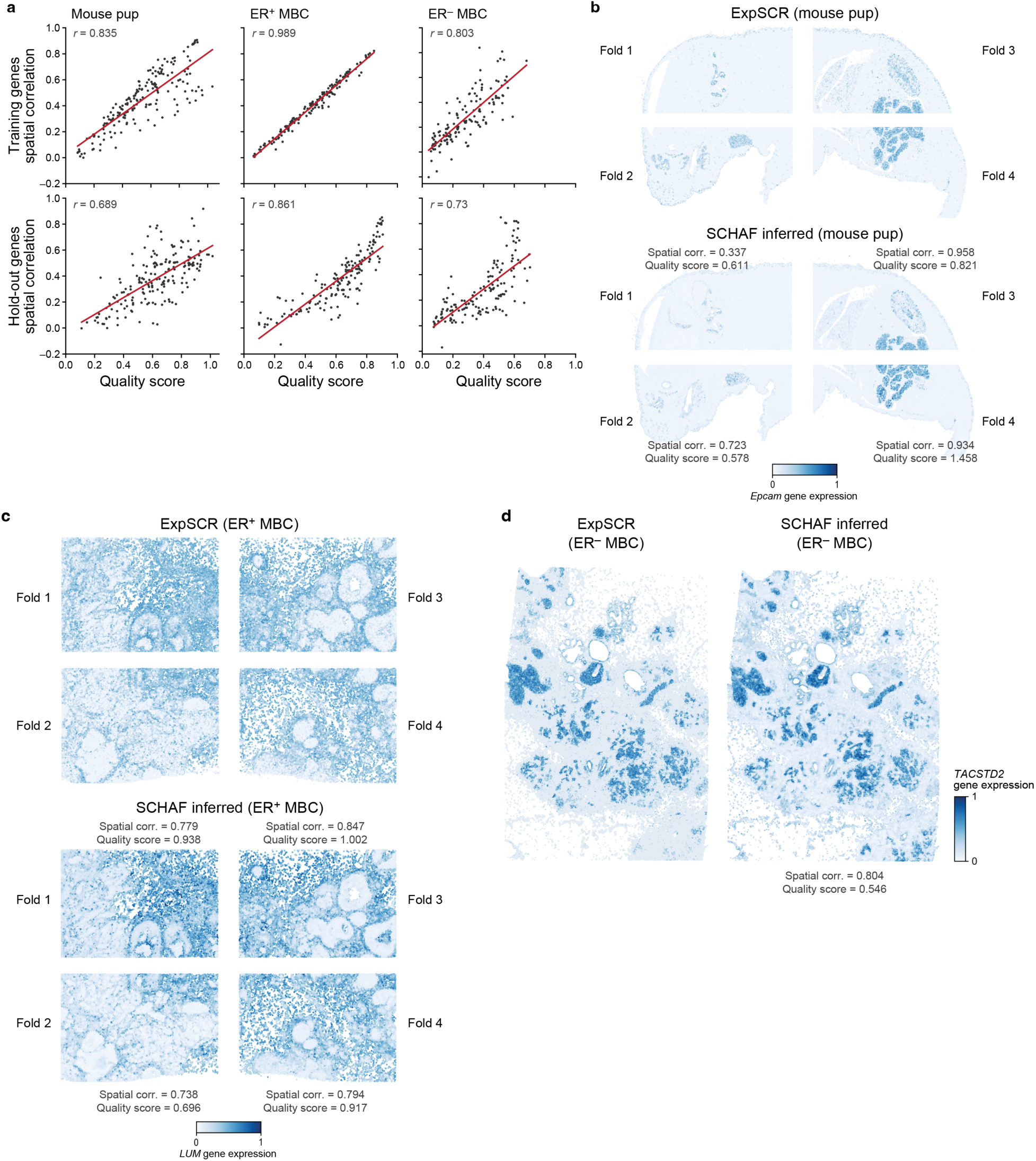
Quality scores anticipate which genes Paired SCHAF infers accurately. **a.** Quality scores agree with spatial correlations. Quality scores (x axis) and spatial correlation between measured and SCHAF-inferred expression levels (y axis) for genes (dots) in training (top) and hold-out (bottom) data in the three datasets. Red line: best fit. **b-d.** Enhanced spatial inference for genes with higher qualitative scores. Expression in each spatially-resolved cell (blue color scale) in real and SCHAF-inferred data for training gene *Epcam* in Mouse Pup corpus (**b**), hold-out gene *LUM* in ER+ MBC (**c**) and hold-out gene *TACSTD2* in ER-MBC (**d**).

The predictive quality score helps identify reliable spatial expression patterns in Paired SCHAF predictions, as seen when comparing ExpSCR measurements with Paired SCHAF inferences for genes with a variety of quality scores (**Fig. 3b-d**, **Extended Data Fig. 1-2**). For example, in the Mouse Pup, for both the epithelial marker gene *Epcam* (PQS=0.337 to 1.458, **Fig. 3b**) and the liver hepatocyte marker *Hmgcs2* (**Extended Data Fig. 1a**, PQS = 0.15 to 1.409), higher PQS is associated with better agreement in complex visual patterns between ExpSCR and Paired SCHAF, and lower PQS with noisier gene expression across the sample. Neuronatin (*Nnat*) (**Extended Data Fig. 1b**), an important gene in brain development, has a broader PQS range (0.206, .238, .763, and 2.661), but even with regions with lower PQS and lower spatial correlation between SCHAF prediction and ExpSCR measurement, Paired-SCHAF captured some highly subtle spatial patterns correctly (**Extended Data Fig. 1b, bottom right**). We observed similar patterns in ER+ MBC for the Lumican gene (*LUM*, PQS = 0.696 to 1.002, **Fig. 3c**), *CD3E* (PQS = 0.626 to 0.809, **Extended Data Fig. 2a**), and lipoprotein lipase (*LPL*) (PQS = 0.268, 0.453, 0.477, and 0.697, **Extended Data Fig. 2b**). Similarly to Neuronatin, *LPL* has sparse punctate expression, which is nevertheless qualitatively predicted by Paired SCHAF (**Extended Data Fig. 2b, bottom right**). In ER-MBC, Paired SCHAF performed better for *TACSTD2*, with strong patterned expression (**Fig. 3d**), less well for *CCDC80* with a somewhat weaker but patterned expression (**Extended Data Fig. 2c**), and worst for *CD68*, with weak and sparse expression (**Extended Data Fig. 2d**). Notably, the PQS is not simply a reflection of expression level *per se*. Thus, a lowly expressed gene that has a distinct spatial pattern, such as *CD3E* (PQS = 0.626 to 0.809, **Extended Data Fig. 2a**), can be predicted well and have a high PQS.

### Paired SCHAF generates high quality whole transcriptome predictions from signature spatial training data

Next, we assessed how well Paired SCHAF predicts expression profiles over the entire transcriptome by comparing its predictions to lower resolution Visium spatial transcriptomics (ExpLR) for ER^+^ MBC. Because these low-resolution data measure aggregate transcriptomes in 55 micron spots, we ‘coarsed-up’ the single cell resolution predictions from Paired SCHAF to a comparable map, summing each predicted transcript expression within a spot.

There was a strong agreement between a gene’s predictive quality score and the spatial correlation between the ExpLR maps and low-resolution Paired SCHAF expression profiles, for genes that were not available for the supervised spatial pretraining phase of Paired SCHAF (**Fig. 4a, Extended Data Fig. 3a**; Pearson’s r= 0.88 over all 17,742 genes measured by ExpLR/Visium). For example, low-resolution expression profiles for Tetraspanin 1 (*TSPAN1)* have a range of predictive quality scores over the four folds (0.62, 0.694, 0.876, 1.038) which correlate with the qualitative agreement between Paired SCHAF low-resolution maps and Visium, with lower spatial correlation in lower PQS folds driven by loss of lower signal by Paired SCHAF predictions (**Fig. 4b**). We observed similar patterns for additional genes (e.g., *ATP2C2* and *CENPL*, **Extended Data Fig. 3b,c**). Notably, in all these cases, the original (pre-coarsing) single cell resolution predictions from Paired SCHAF add substantial definition compared to the low-resolution Visium maps (**Fig. 4b,d**, **Extended Data Fig. 3b,c**).

**Figure 4.**
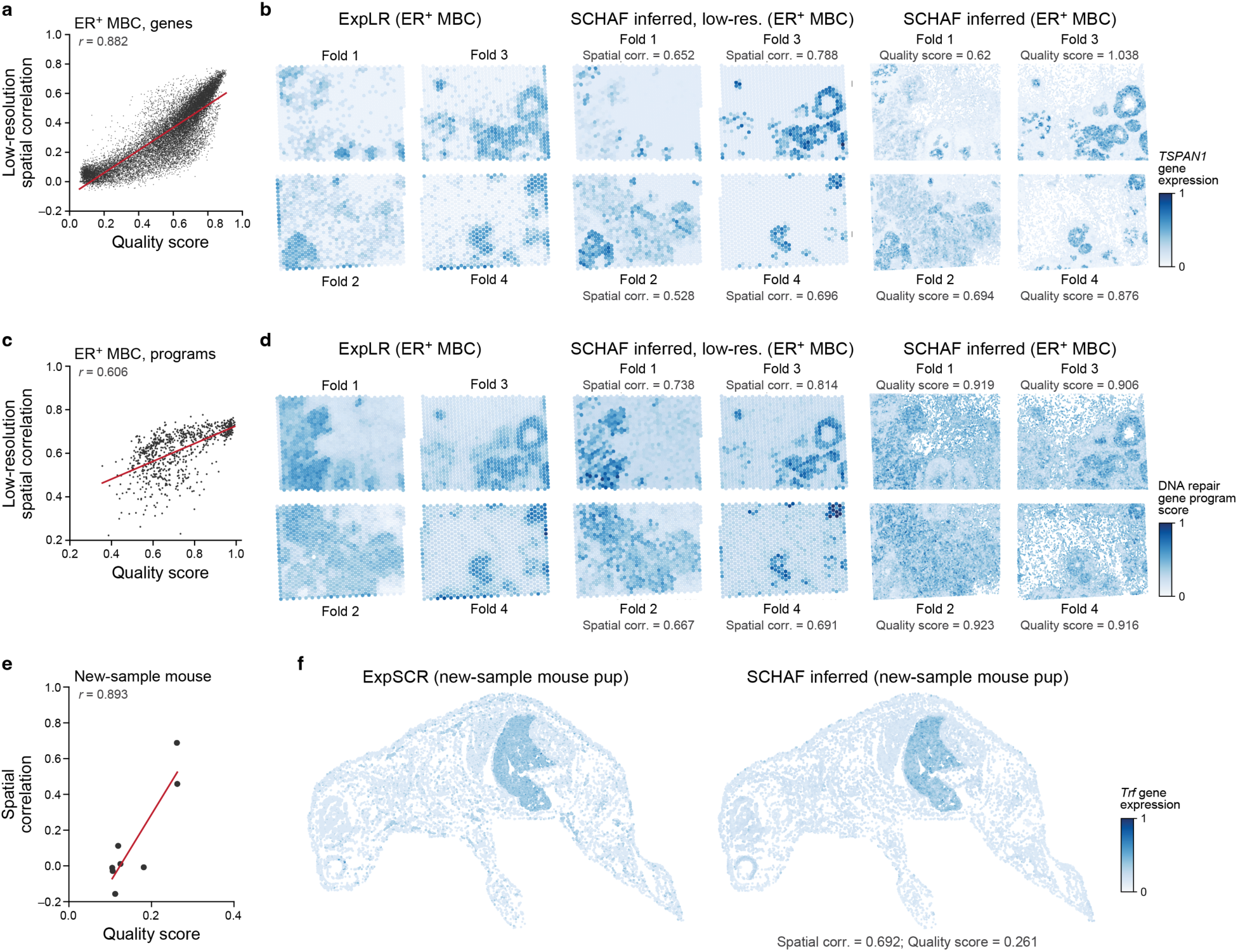
Paired SCHAF infers the whole transcriptome accurately. **a,c.e.** Quality scores agree with the degree of spatial correlation. Quality scores (x axis) and extent of spatial correlation between measured and Paired SCHAF values (y axis) for the ER+ MBC dataset genes (a), the ER+ MBC gene programs (c), or for each gene measured by ExpSCR (ISH) in a new mouse pup sample (e). Low-resolution spatial correlation is Pearson’s *r* across ExpLR spots. Dashed red line: best fit. **b,d,f**. Illustrative examples of SCHAF-inferred spatial measurements. **b,d**. ExpLR-measured (left), low-resolution (middle), and single cell resolution (right) SCHAF inferred spatial levels (color bar) for *TSPAN1* (b) and the DNA Repair gene program (d) for each of four folds. Quality scores and low-resolution spatial correlations are noted. **f.** Measured (ISH, left) and SCHAF-inferred (right) spatial expression levels for *Trf*.

Because lowly and sparsely expressed individual genes may be captured more poorly by spatial transcriptomics and predicted less robustly by Paired SCHAF, we reasoned that the spatial expression of co-functional or co-regulated gene programs in transcriptome-wide data would be more robust and better predicted. To this end, we analyzed 797 cancer gene programs^37,38^, whose genes are well-covered by Paired SCHAF predictions (>90% gene overlap, **Methods**), scoring the overall expression of each gene program (**Methods**) in each spot in the measured and predicted maps. Notably, the spatial correlation values between measured and predicted values increased (compare **Extended Data Fig. 3a and 4a**, P=0.0, Mann-Whitney U test), with the vast majority showing good positive correlation (772 (96.6%) programs *vs*. 6,858 (38.7%) of the constituent program genes with spatial correlation r>0.4). We further defined the program’s predictive quality score as the standard deviation of its program score over the cells in the Paired SCHAF generated data. There was a moderately strong relationship between a program’s PQS and the spatial correlation (Pearson’s r=0.606, **Fig. 4c**). This was also qualitatively reflected in the spatial patterns of multiple programs, including a DNA Repair program (**Fig. 4d**), a Tumor Epithelial gene program (**Extended Data Fig. 4b**) and a Serine Proteases program (**Extended Data Fig. 4c**).

### *In situ* hybridization experiments validate Paired SCHAF predictions

To determine whether Paired SCHAF can accurately predict high resolution expression profiles for genes that are not available for model training, we selected eight genes that were neither training nor hold-out genes and that had a wide range of predictive quality scores across the mouse pup. We then obtained a new tissue slice for *in situ* hybridization (ISH), spanning the entirety of another (new) one-day old mouse pup. We developed a new protocol to perform both H&E and ISH from the same slice (**Methods**) and performed this for the eight genes in the same slice. We trained a Paired SCHAF model on all data from the original mouse pup and used it to predict the expression values of these eight genes from the new pup’s H&E slice. Because the H&E and ISH assays were performed on the same slice, we could calculate exact, single cell resolution spatial correlations between Paired SCHAF predictions and ExpSCR (ISH) measurements.

The spatial correlation was in strong agreement with the predictive quality score for the eight genes, such that genes with higher predictive quality scores indeed had much higher spatial correlation between Paired SCHAF predictions and experimental measurements (**Fig. 4e**). These were also supported by qualitative assessment of images of Paired SCHAF predicted gene expression and the experimental maps from ISH experiments for genes from different tissues (**Fig. 4f, Extended Data Fig. 5**). For example, *Trf*, a liver marker with a relatively high quality score (0.261) and spatial correlation (0.692) also showed a stronger qualitative agreement than the cardiomyocyte and skeletal muscle marker *Csrp3* (0.262 and 0.462, respectively) and the sparsely expressed neural marker *Rtn1* (0.124, 0.015) (**Fig. 4f, Extended Data Fig. 5**). Nevertheless, for *Csrp3*, there was substantial qualitative similarity between ISH measurement and Paired SCHAF predicted patterns, suggesting that Paired SCHAF captured the subtler patterns. *Rtn1*, with a low PQS, had a low spatial correlation. Both ExpCSR and SCHAF show sparse, scattered expression but not in the same specific cells (**Extended Data Fig. 5**).

### Unpaired SCHAF infers single cell distributions of gene expression

Unlike Paired SCHAF, which requires spatial transcriptomics for training, Unpaired SCHAF is trained only with H&E images and sc/snRNA-seq data (**Fig. 1a**). Unpaired SCHAF training can optionally incorporate cell type annotations when available for both scRNA-seq data (from expression) and H&E data (via pathologist annotations), to encourage correspondence between input tiles and their corresponding prediction (**Supplementary Fig. 3**). Thus, Unpaired SCHAF uses an H&E image to infer the distribution of RNA transcripts in a given tissue sample.

To assess Unpaired SCHAF, we relied on three human tissue datasets, where both sc/snRNA-Seq and H&E histology were available for training. Two datasets were from human tumors. As non-canonical tissues, tumors are a challenging use case for prediction on new specimens (new tumor from a new patient). The **small cell lung cancer (SCLC)** dataset consisted of 24 tumors profiled by snRNA-seq and matching H&E stains from an unpublished study from Memorial Sloan Kettering Cancer Center (MSKCC). The snRNA-seq data were collected from a portion of the same slice used in H&E staining. No cell type annotations were performed on these H&E data. The **metastatic breast cancer (MBC)** dataset consisted of nine patient samples from the Human Tumor Atlas Project Pilot (HTAPP)^39^, where each pair of H&E and snRNA-seq/scRNA-seq data came from different biopsies of the one metastasis. We also included previously collected spatial profiling by MERFISH^13^ (used for validation), on another adjacent section. In addition, we generated coarse cell type annotations by an expert pathologist for the H&E sections, broadly delineating large regions that contain mostly cells of the same type. The third dataset was from four **human placenta** samples, which we previously profiled by scRNA-seq^40^ and added here matching whole-slide H&E stains from an adjacent section with detailed cell type annotations at nearly single cell resolution, newly generated by a clinical pathologist with expertise in placental histopathology. We also had previously collected spatial profiling by STARmap (used for validation)^40^, using a portion of another adjacent section. For each tissue sample, we evaluated Unpaired SCHAF via leave-one-out cross validation (LOOCV), where we trained on all but one of the samples in the dataset and evaluated on the sample excluded from training. The reported results are the average across all patient samples. We incorporated cell types into training Unpaired SCHAF for the two datasets (MBC/HTAPP and placenta) with expert histopathologist annotations.

Unpaired SCHAF generates adequate single cell profiles from H&E based on its ability to correctly estimate (**1**) gene expression distributions across single cells, (**2**) distribution (composition) of cell types and psuedobulk profiles; (**3**) heterogeneity of expression profiles within each cell type; and (**4**) appropriate correlation structure and clustering. First, we quantified the difference between Unpaired SCHAF predicted expression distributions and the ground truth distributions for a given gene using the Earth Mover’s Distance (EMD), a widely used quantitative measure of similarity between two continuous, nonparametric distributions (**Methods**; low values indicate high similarity). The average gene EMD between Unpaired-SCHAF predictions and real sc/snRNA-Seq was very low in each of the three datasets (**Fig. 5a**; EMD=0.09 (MBC), 0.054 (Placenta), 0.062 (SCLC), with many genes’ EMD nearly at zero, showing that most genes had predicted distributions very similar to their measured scRNA-seq distributions. Consistently, the gene-gene correlation structures were relatively preserved, with some variation between datasets (Placenta performing best) and between samples within a dataset (**Supplementary Fig. 4**). Second, the cell type distributions of Unpaired SCHAF predictions were similar to those from measured sc/snRNA-seq, based on the Jensen-Shannon Divergence (JSD, **Methods**), with cell types assigned by a previously trained neural network classifier to the Unpaired SCHAF generated data (**Extended Data Fig. 6a, b**). The pseudobulk profiles across different cell types were typically well correlated between Unpaired SCHAF inferred data and measured scRNA-seq (**Supplementary Fig. 5**), with some notable exceptions (e.g., HTAPP 4531). Third, the extent of gene expression heterogeneity within each cell type using the same heterogeneity score as in Paired SCHAF (**Methods**) was comparable between generated and real sc/snRNA-seq, with more disparity in the Placenta dataset than MBC or SCLC (**Supplementary Fig. 6**), and the distribution of EMD for genes was typically low at a cell type specific level (**Supplementary Fig. 7, 8**). Fourth, overall, the relationship between generated profiles is preserved in terms of the extent of clustering, both locally (coherence of clusters) and globally (cluster separation) and as reflected visually in low dimensional embeddings (**Fig. 5b, Extended Data Fig. 6c, d, e**). Heatmaps summarizing these metrics highlight varying degrees of consistency in different samples (**Supplementary Fig. 9**).

**Figure 5.**
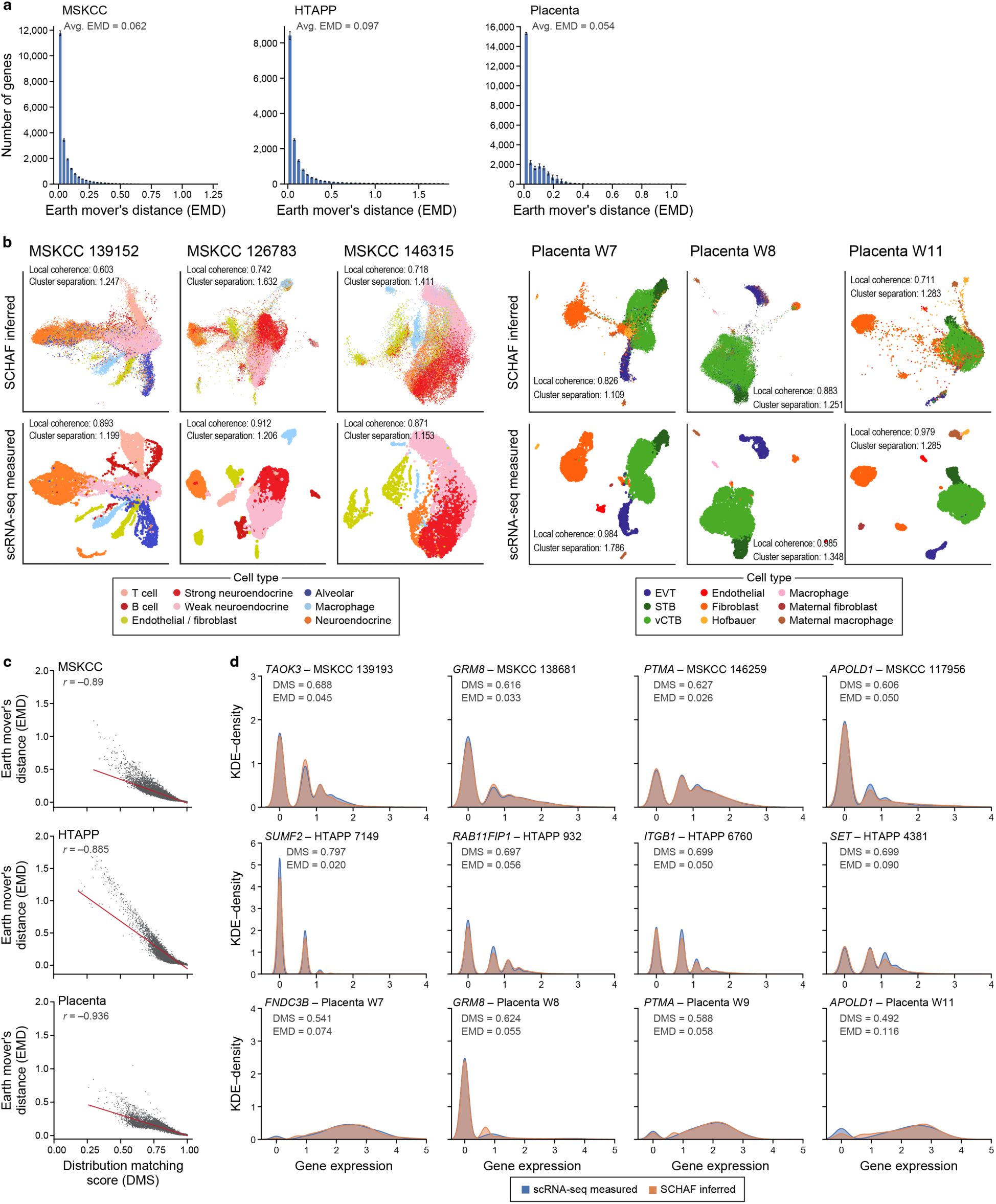
Unpaired SCHAF inferences are similar to measured scRNA-seq. **a.** Most genes have similar expression distributions across cells in measured and SCHAF-inferred scRNA-Seq. Distribution of Earth Mover’s Distance (EMD) between the measured and SCHAF-inferred expression distributions of genes in the SCLC (MSKCC, left), breast cancer (HTAPP, middle) and placenta (right) datasets, in the cross validation (holding out one sample at a time). Error bars: standard error of samples assessed by cross validation. **b.** Representative UMAPs. UMAP embedding of measured (bottom) or Unpaired SCHAF-inferred (top) scRNA-seq profiles (dots) colored by cell types (legends), for three representative samples from each of the placenta (right) or SCLC (MSKCC, left) datasets. The measured and inferred scRNA-seq are projected in a common space. **c,d.** Agreement of expression distributions. **c.** Distribution Matching Score (DMS, x-axis) and Earth Mover’s Distance (EMD, y-axis) between the measured and SCHAF-inferred expression distribution of each gene (dot). Red line: best fit. **d.** KDE plots for measured (blue) and SCHAF inferred (orange) scRNA-seq gene expression for selected genes (labeled on top) from the SCLC (MSKCC, top), breast cancer (HTAPP, middle) and placenta (bottom) datasets. EMD and DMS values are denoted.

While no other methods explicitly solve the problem addressed by Unpaired SCHAF (to the best of our current knowledge), we compared it to several state-of-the-art methods for generalized domain translation. Because none can exploit cell type annotation in their training, we compared them to Unpaired SCHAF trained without cell type information. Across both gene level Earth Mover’s Distance and cell type distribution Jensen-Shannon Divergence criteria, Unpaired SCHAF outperformed competing methods in all corpora (**Supplementary Fig. 10a**). On other criteria (pseudobulk profiles, gene-gene correlation structure, etc.), unpaired SCHAF performed on par with or better than benchmarks (**Supplementary Fig. 10b**).

### A gene-specific distribution matching score identifies gene distributions that Unpaired SCHAF will accurately predict

We next developed a dedicated per-gene prediction quality score to *a priori* anticipate when the generated expression distribution from Unpaired SCHAF is similar to the measured sc/snRNA-seq. Unlike Paired SCHAF, where a low standard deviation was empirically associated with poor predictions (**Fig. 3a**), in Unpaired SCHAF, we reasoned that a high standard deviation (more than mean expression or CV), typically associated with more lowly expressed (and less reliably measured) genes in scRNA-seq^41^, is associated with lower prediction accuracy. Accordingly, we defined a Gene Distribution Matching Score (GDMS) as one minus one-half the standard deviation of that gene’s Unpaired SCHAF inferred expression, which ensures, given that the maximum standard deviation over inferred values is 2, that the score is always non-negative.

Indeed, in all three datasets, the GDMS is highly negatively correlated with EMD distance between Unpaired SCHAF predictions and measured sc/snRNA-seq (**Fig. 5c**; Pearson’s r=-0.885 to - 0.936), such that a high GDMS is associated with a successful prediction of expression distributions. This is also supported when examining expression distributions of individual genes across all three datasets and multiple GDMS scores (**Fig. 5d**, **Extended Data Fig. 7**). Genes with complex multimodal distributions but high GDMS (>0.49) were well predicted by SCHAF (EMD=0.02-0.12) (**Fig. 5d**, **Extended Data Fig. 7** top row), whereas those with low GDMS often resulted in poor prediction (**Extended Data Fig. 7**, bottom four rows, e.g., *LRMDA* in SCLC).

### Unpaired SCHAF inferences spatially agree with pathologist annotations and marker genes

Finally, we assessed the spatial accuracy of Unpaired SCHAF predictions – trained with no spatial molecular data, focusing on its ability to generate spatially accurate cell type assignments. We generated single cell RNA expression data using Unpaired SCHAF for each cell in the H&E section, assigned a cell type to the generated profiles and compared these to expert pathologist annotations.

We found strong quantitative agreement in the placenta dataset and a weaker – but still marked – agreement in the less canonical HTAPP tumor metastases (**Fig. 6**, **Supplementary Fig. 11**). In the placenta, there was also strong qualitative agreement between spatial cell type annotations from Unpaired SCHAF inferences and from the expert pathologist (**Fig. 6a**), especially if pathologist cell type annotations were included in the training (**Supplementary Fig. 12**). In all four samples, inferences of major cell types in the placenta, such as extravillous trophoblasts (EVTs), syncytiotrophoblasts (STBs), villous cytotrophoblast (vCTBs), fibroblasts, and decidua, are spatially accurate (**Fig. 6a**), as also reflected in confusion matrices (**Supplementary Fig. 11b**), albeit with some specific challenges for particular cell types in some samples. In particular, in the W9 and W11 samples, idiosyncratic regions populated by mostly decidua, fibrin, and maternal cells were well captured by Unpaired SCHAF. In W7, W8, and W9 samples, small but dense regions of EVTs were also captured well. In the W8 and W11 samples, the sparse spatial patterning of fetal blood and endothelial cells is correctly present. The two main cell groups across all samples, one consisting of fetal fibroblasts and Hofbauer cells, and the other of STBs and vCTBs, are both inferred well by Unpaired SCHAF (**Fig. 6a**). Ablating cell type information from training lowered this spatial agreement (**Supplementary Fig. 12**). In the tumor dataset, where samples are far less canonical, there was still notable but weaker qualitative agreement with pathologist annotations in three complex samples (**Supplementary Fig. 11**a,c), with similar reduced performance when cell type information is ablated (**Supplementary Fig. 13**). Note that elementary image analyses show only a weak, if any, correspondence between the number of cells per tile and Unpaired SCHAF’s inferred spatial cell type maps (**Supplementary Fig. 14-17**), suggesting that Unpaired SCHAF is learning high complexity features, rather than merely cell type dependent density.

**Figure 6.**
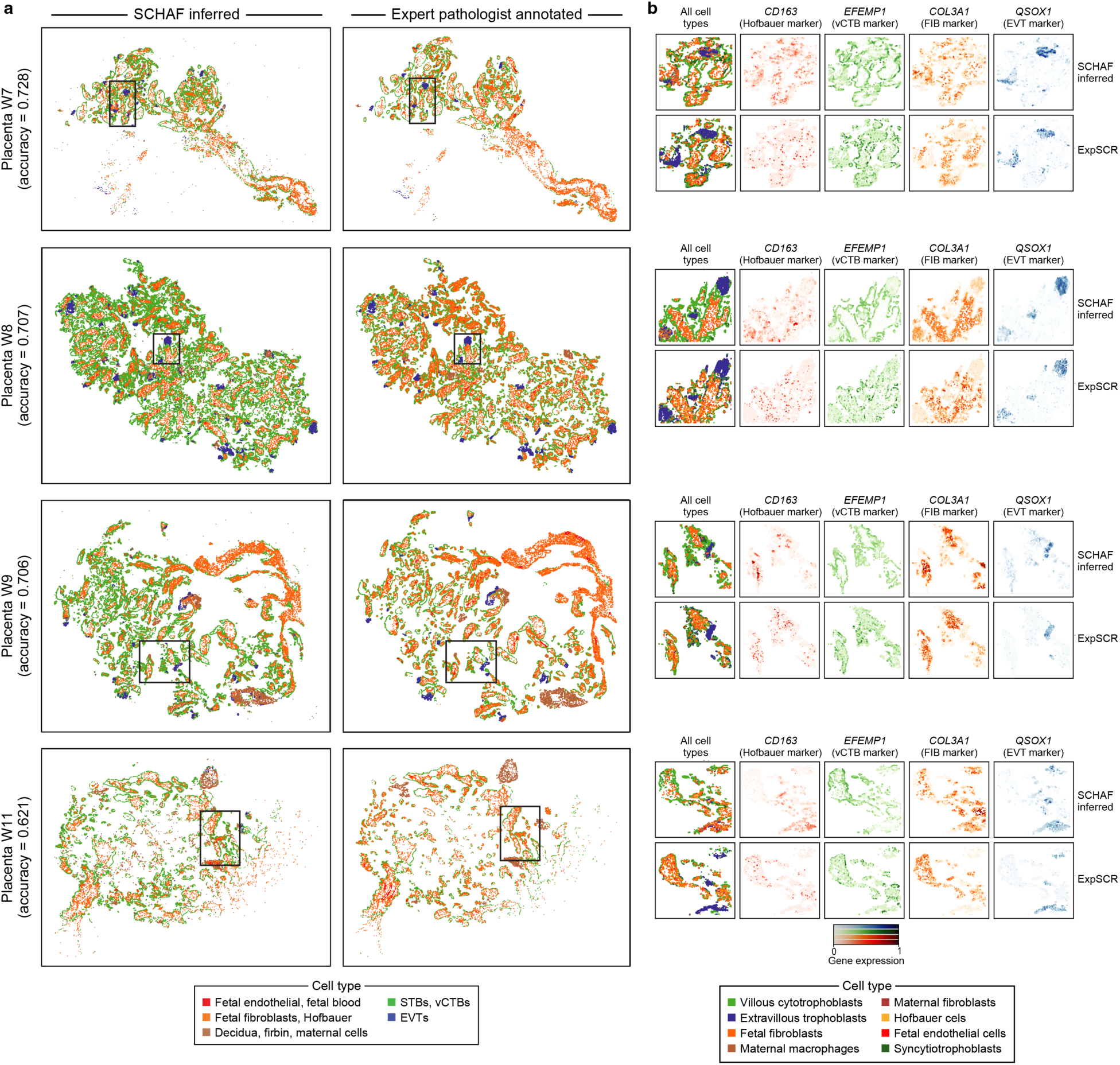
Unpaired SCHAF inferences spatially match expert pathologist annotations and STARmap spatial measurements in the human placenta. **a.** Agreement with pathologist annotations. Four human placenta samples (rows) colored by annotations from a cell type classifier of Unpaired SCHAF inferences (left) or an expert pathologist (right). **b.** Agreement with STARmap measurements. Fragments from four human placenta samples (as boxed in **a**), measured by ExpSCR (STARmap), colored by cell type annotations and marker gene expression, in both Unpaired-SCHAF-inferred (top) and ExpSCR (STARmap, bottom) measured profiles.

We further compared SCHAF predictions to cell type annotations determined by ExpSCR (MERFISH for HTAPP, STARmap for placenta) and found relatively compelling qualitative and quantitative results in the placenta and HTAPP datasets (**Supplementary Fig. 18, 19**). Since the H&E and ExpSCR data for each sample came from adjacent, but not the same, slices, we could perform cell-to-cell mapping between the two. To address this, we aligned lower-resolution spots (∼5 cells on average), handling our ExpSCR data as ExpLR data and comparing to coarsed-up (low-resolution) SCHAF predictions. In the Placenta, qualitative and quantitative comparison show substantial, but imperfect agreement (including due to the loss of signal due to coarsing up) (**Supplementary Fig. 18c**). In the HTAPP data, the agreement for three complex samples with multiple cell types was weaker, especially struggling with immune cells, although still notable (**Supplementary Fig. 19c**).

Unpaired SCHAF also generated correct spatial patterns for more specific cell types derived from ExpSCR (STARmap) (**Fig. 6b**) in addition to four marker genes associated with various annotation categories: *QSOX1* (EVT), *COL3A1* (fibroblasts), *EFEMP1* (vCTB) and *CD163* (Hofbauer), by qualitative comparison to STARmap^21^ measurements in adjacent portions (**Fig. 6b**; sections could not be fully matched up at exact single cell granularity). Overall, more than 100 genes were well captured in the placenta as assessed by comparison to ExpLR measurements (low resolution STARmap) (**Supplementary Fig. 20**). In HTAPP, results were weaker and very variable across tumors (**Supplementary Fig. 21**a-c). Notably, Unpaired SCHAF trained without cell types inferred few (in HTAPP) to no (in placenta) genes with a correlation over 0.4 (**Supplementary Fig. 21d**). Ablating cell type information from training also lowered spatial agreement with pathologist annotations (**Supplementary Fig. 12, 13**).

## DISCUSSION

We describe SCHAF, a computational framework to predict spatially resolved single cell omics profiles from H&E images. SCHAF is the first method to infer high gene throughput, single cell resolution, spatially resolved (in the case of Paired SCHAF) transcriptomic data using only histology images, a relatively low-cost technique in wide clinical practice. To address the range of opportunities for training, we present two versions of SCHAF: Paired SCHAF, which requires limited molecular spatial data in training, but has spatial accuracy guarantees for its predictions, and Unpaired SCHAF, which, does not require any spatial transcriptomic data during training, but at the cost of making predictions with weaker spatial guarantees. We demonstrate SCHAF across both mouse and human data, from both healthy, canonical developmental tissues and from non-canonical human tumors. We assess its predictions in terms of genes, gene programs, and cell types in single cell and spatial patterns with experimental validation by sc/snRNA-seq, ISH, Xenium, STARmap, MERFISH, and Visium.

Paired SCHAF leverages a state-of-the-art foundation^32^ model based on a vision transformer architecture to match the spatial transcriptomic data. Instead of supervised training using a small number of spatial marker transcripts, Paired SCHAF augments supervised training with information obtained by mapping scRNA-seq data to spatial positions in the histology image. This allowed the model to generate highly quality spatial, transcriptome-wide profiles. While many spatial transcriptomics methods only measure a small portion of tissue, H&E can be found in a much larger tissue view. Accordingly, spatial transcriptomics from a small part of tissue can be used to train Paired SCHAF, which can then be used to extrapolate spatial data for the entire tissue during inference. Indeed, Paired SCHAF was able to accurately infer large parts of a tissue sample not seen in training and generalize to entirely new tissue samples. SCHAF produces data that is higher resolution and more realistic than extant methods trained to infer low-resolution Visium data from H&E. Our prediction quality score helps users identify *a priori* genes and gene programs that are likely well-predicted by Paired SCHAF.

Unpaired SCHAF leverages adversarial deep learning and foundation models and does not require spatial transcriptomic data to train. It also performed quite well in generating accurate gene distributions and cell type compositions, producing high gene throughput, single cell resolution expression data from an H&E stain. The Gene Distribution Matching Score allows a user to *a priori* determine the genes that are predicted well. By incorporating expert histopathologist annotations into our training, when including expert pathologist cell annotations in the training, Unpaired SCHAF was able to spatially replicate both expert pathologist annotations and marker gene expression measured via STARmap on unseen tissues.

SCHAF was successful on corpora of matched H&E and transcriptomics data from a wide variety of tissue types. As diseased tissues are highly variable and idiosyncratic, predicting a new sample’s profile, even of the same type, is a challenging task. At present, the paucity of organized multi-modal datasets limits the generalizability of models like SCHAF across new tissue types. Future studies are required to assess the ability to train a single model across a single corpus of multiple types of tissues, tumors, or technologies (e.g., snRNA-Seq, scATAC-Seq, and spatial multiomics). Doing so would require additional organized spatial datasets with matched single cell profiling and histology imaging, which are still surprisingly scarce, but are growing thanks to efforts, such as the Human Tumor Atlas Network^42^, HubMAP^43^, CZI Virtual Cell initiative^44^, and the Human Cell Atlas^4,2^. To encourage this, we have released the “SCHAF-1M” data repository with this study, encapsulating all the data analyzed here and used in training, as a resource to the broader community.

The main experimental limitation of Paired SCHAF lies in the requirement of expensive spatial transcriptomic data in training and/or to test the spatial accuracy of its predictions. To guarantee spatial accuracy, spatial transcriptomic data must be incorporated in training Paired SCHAF. Even in Unpaired SCHAF, spatial accuracy of cell type level predictions required expert histopathologist annotations in the training. Future work should aim to minimize the number of genes required in training models like Paired SCHAF.

SCHAF’s success motivates the use of a modality-agnostic, lower-dimensional latent space in computational techniques for a multitude of applications in data integration and modality transfer. The underlying principles defining SCHAF can be extended to other cases in biology, both with similar data modalities (*e.g.*, cell profiles and microscopy images measured in cultured cells^45^, single cell and spatial multi-omics) in different biological modalities (such as temporal tracing^46^), leading to models producing multimodal outputs from one readily-accessible input. More fundamentally, interpreting SCHAF’s predictions could have implications for our understanding of tissue biology, including how molecular information leads to tissue structures and vice versa.

Finally, SCHAF’s ability to produce high quality transcriptomic data *in silico* from histological imaging would significantly increase the accessibility of high-quality molecular profiles. Beyond the savings in time, effort, and expense, SCHAF could be used to unlock molecular information for the vast number of clinical samples archived over decades and allow scientists to benefit from the insights provided by high-resolution methods in low resource settings. Furthermore, clinical biopsies, yielding minimal patient measurements, could leverage SCHAF to glean more useful clinical information in minimal time and sample access. SCHAF, being the first method to generate high gene throughput, single cell resolution, spatially resolved (in the case of Paired SCHAF) transcriptomic data at the relatively low cost of a histology stain, is well poised to do such.

## Supplementary Figure Legends

**Supplementary Figure 1. Paired SCHAF preserves expression heterogeneity within cell types.**

Heterogeneity (y axis, **Methods**) of expression profiles within each cell type (*x* axis) in measured (ExpSCR, blue) and SCHAF-inferred (orange) profiles in the mouse pup (left), ER+ MBC (middle) and ER-MBC (right) datasets. Error bars: standard error of folds assessed by cross validation.

**Supplementary Figure 2. Benchmarking Paired SCHAF inferences with other methods**

Average spatial correlation across spatially-measured genes (**a,** y axis) and cell type accuracy (**b**, y axis; defined as proportion of cells in each fold assigned to the correct cell type label) for inferences by Paired SCHAF (red), SPiRiT^35^ (dark blue), ST-Net^31^ (light blue), DeepPT^36^ (purple), and HE2RNA^30^ (green) in each dataset (x axis) for the mouse pup (left), ER+ MBC (middle) and ER-MBC (right) datasets. Error bars: standard error of folds assessed by cross validation.

**Supplementary Figure 3. Unpaired SCHAF learns to predict a tissue’s single cell omics data from its H&E image**

Detailed overview of Unpaired SCHAF training (top and middle) and inference (bottom).

**Supplementary Figure 4. Characteristics of scRNA-seq profiles inferred by Unpaired SCHAF**

**a.** Agreement with pseudobulk RNA-Seq. Correlation (Pearson’s r, y axis) between pseudobulk profiles from the measured and Unpaired SCHAF inferred scRNA-Seq for each sample (x axis) in the placenta (right), MBC (HTAPP, middle) and SCLC (MSKCC, left) datasets. **b.** Preservation of heatmaps of order-ranked entries of gene-gene correlation matrices in inferred profiles. Heatmaps, as described above (and in **Methods**), were formed from the 100 most highly variable genes in each sample’s scRNA-seq data, from SCHAF inferred (top) and scRNA-seq (bottom) from three samples from each of the Placenta (bottom), MBC (HTAPP, top right), and SCLC (MSKCC, top left) datasets. Measured heatmaps are clustered and the same order is imposed on the SCHAF heatmaps. Samples’ heatmap meta correlations (Pearson’s *r* between corresponding flattened entries of the SCHAF and scRNA-seq heatmaps) are noted, **c.** Agreement of meta correlations. Heatmap meta correlations (y axis, **Methods**, described above) for each sample (x axis) in the placenta (right), MBC (HTAPP, middle) and SCLC (MSKCC, left) datasets.

**Supplementary Figure 5. Cell type pseudobulk profiles inferred by Unpaired SCHAF**

**a.** Cell type pseudobulk agreement across samples. Cell type pseudobulk correlation score (y axis, **Methods**), with dots representing each cell type’s pseudobulk correlation and error bars capturing standard error across cell types, between cell type specific pseudobulk profiles from measured and SCHAF inferred scRNA-seq in each sample (x axis) in the Placenta (right), MBC (HTAPP, middle), and SCLC (MSKCC, left) datasets. **b.** Cell type pseudobulk agreement in representative samples. Correlation (Pearson’s *r*, y axis) between pseudobulk profiles from measured and SCHAF inferred scRNA-seq in each cell type (x axis) in three representative samples from the Placenta (bottom), MBC (HTAPP, middle), and SCLC (MSKCC, top).

**Supplementary Figure 6. Unpaired SCHAF preserves the extent of expression heterogeneity within cell types**

Mean heterogeneity (x axis, **Methods**; across samples from cross validation) of expression profiles within a cell type (*x* axis) in measured (blue) and Unpaired SCHAF-inferred (orange) scRNA-seq for SCLC (MSKCC, left), MBC (HTAPP, middle), and placenta (right). Error bars: standard error of samples assessed by cross validation.

**Supplementary Figure 7. Agreement of gene expression distributions in each cell type in Unpaired SCHAF**

Distribution of Earth Mover’s Distance (EMD, x axis) between the measured and Unpaired SCHAF-inferred expression distributions of each gene in each cell type (label on top) in the breast cancer (HTAPP, left), placenta (middle) and SCLC (MSKCC, right) datasets, in the cross validation (holding out one sample at a time). Error bars: standard error of samples assessed by cross validation.

**Supplementary Figure 8. Agreement of highly variable gene expression distributions in each cell type in Unpaired SCHAF**

Distribution of Earth Mover’s Distance (EMD, x axis) between the measured and Unpaired SCHAF-inferred expression distributions of each of highly variable genes in each cell type (label on top) in the breast cancer (HTAPP, left), placenta (middle) and SCLC (MSKCC, right) datasets, in the cross validation setting (holding out one sample at a time). Error bars: standard error of samples assessed by cross validation. Highly variable genes: a gene that is found to be one of the 500 most highly variable in any sample of a dataset.

**Supplementary Figure 9. Aggregation of evaluation metrics for Unpaired SCHAF**

Heat maps displaying values of metrics used in evaluating Unpaired SCHAF across samples in breast cancer (HTAPP, left), SCLC (MSKCC, middle), and placenta (right) datasets.

**Supplementary Figure 10. Benchmarking Unpaired SCHAF inferences**

Mean Earth Mover’s Distance from measured scRNA-seq (EMD, y axis, **a**, left), Jensen-Shannon Divergence from measured scRNA-Seq (JSD, **a**, y axis, right), mean correlation to measured pseudobulk RNA-seq (Pearson’s r, y axis, **b**, left), mean cell type pseudobulk correlation (Pearson’s r, y axis, **b**, middle), and gene-gene correlation heatmap meta correlation (Pearson’s r, y axis, **b**, right), for scRNA-seq profiles inferred by Unpaired SCHAF (without cell types in training) (dark blue) and other methods (a-c, color legends), in each data set (x axis). Error bars: standard error of samples assessed by cross validation.

**Supplementary Figure 11. Comparison of Unpaired SCHAF spatial inferences to expert pathologist annotations**

**a.** Global spatial accuracy. Proportion of cells (y axis) with cell types assigned to SCHAF inferences that agree with expert pathologist annotations in each sample (x axis) from the Placenta (left) and MBC (HTAPP, right) datasets. **b**. Cell type specific accuracy in the placenta. Number of cells with each pathologist annotation (rows) and SCHAF-inferred annotation (columns) in each placenta sample. **c.** Regional accuracy in relatively complex MBC samples. Left: Sample sections with cells (left) or regions (right) colored by one of five region categories based on SCHAF-inferred (left) or coarse pathologist (right) annotations. Right: Number of cells with each pathologist annotation (by region; rows) and SCHAF-inferred annotation (columns) in the corresponding MBC sample.

**Supplementary Figure 12. Unpaired SCHAF spatial annotations match expert pathologist annotations in the placenta better when trained with cell type information**

**a.** Global spatial accuracy. Proportion of cells (y axis) with cell types assigned to SCHAF inferences that agree with expert pathologist annotations in each sample (x axis) from the Placenta, when SCHAF is trained with (blue) and without (orange) cell types in training. **b.** Human placenta samples (rows, labeled on left) colored by annotations from an expert pathologist (right) or based on Unpaired SCHAF data, with models trained with (middle) and without (left) cell type information. **c.** Cell type specific accuracy in the placenta. Number of cells with each pathologist annotation (rows) and SCHAF-inferred (without cell types in training) annotation (columns) in each placenta sample.

**Supplementary Figure 13. Unpaired SCHAF spatial annotations match expert pathologist annotations in MBC (HTAPP) better when trained with cell type information**

**a.** Global spatial accuracy. Proportion of cells (y axis) with cell types assigned to SCHAF inferences that agree with expert pathologist annotations in each sample (x axis) from MBC (HTAPP) samples, when SCHAF is trained with (blue) and without (orange) cell types in training.

**b.** MBC (HTAPP) samples (rows, labeled on left) colored by annotations from an expert pathologist (right) or based on Unpaired SCHAF data, with models trained with (middle) and without (left) cell type information. **c.** Cell type specific accuracy in MBC (HTAPP). Number of cells with each pathologist annotation (rows) and SCHAF-inferred (without cell types in training) annotation (columns) in each placenta sample.

**Supplementary Figure 14. Comparison of spatial cell types from Unpaired SCHAF inferences to cell neighborhood density in the placenta**

**a.** Placenta sections from each sample (rows, labeled on left) colored by SCHAF inferred cell types (left) or the number of cells in each cell’s H&E tile (right, color bar at bottom). **b**. Distribution of number of cells per H&E tile (y axis) in cells annotated by Unpaired SCHAF to each category (x axis, color legend). **c**. Distribution of number of cells per H&E tile (y axis) in cells annotated by a pathologist to each category (x axis, color legend).

**Supplementary Figure 15. Comparison of spatial cell types from Unpaired SCHAF inferences to RGB value intensity in the placenta**

**a.** Placenta sections from each sample (rows, labeled on left) colored by SCHAF inferred cell types (left) or by mean RGB value intensity in each cell’s H&E tile (right, color bar at bottom). **b**. Distribution of mean RGB value intensity per H&E tile (y axis) in cells annotated by Unpaired SCHAF to each category (x axis, color legend). **c**. Distribution of mean RGB value intensity per H&E tile (y axis) in cells annotated by a pathologist to each category (x axis, color legend).

**Supplementary Figure 16. Comparison of spatial cell types from Unpaired SCHAF inferences to cell neighborhood density in MBC (HTAPP)**

**a.** MBC (HTAPP) sections from each sample (rows, labeled on left) colored by SCHAF inferred cell types (left) or the number of cells in each cell’s H&E tile (right, color bar at bottom). **b**. Distribution of number of cells per H&E tile (y axis) in cells annotated by Unpaired SCHAF to each category (x axis, color legend). **c**. Distribution of number of cells per H&E tile (y axis) in cells annotated by a pathologist to each category (x axis, color legend).

**Supplementary Figure 17. Comparison of spatial cell types from Unpaired SCHAF inferences to RGB value intensity in HTAPP**

**a.** MBC (HTAPP) sections from each sample (rows, labeled on left) colored by SCHAF inferred cell types (left) or by mean RGB value intensity in each cell’s H&E tile (right, color bar at bottom).

**b.** Distribution of mean RGB value intensity per H&E tile (y axis) in cells annotated by Unpaired SCHAF to each category (x axis, color legend). **c**. Distribution of mean RGB value intensity per H&E tile (y axis) in cells annotated by a pathologist to each category (x axis, color legend).

**Supplementary Figure 18. Comparison of Unpaired SCHAF inferences to spatial STARmap cell type annotations in the placenta.**

**a.** Global spatial cell type accuracy. Proportion of ExpLR (low-resolution (coarsed up) STARmap) spots (y axis) with cell types that match those on (coarsed up) SCHAF inferences (x axis) for the portion of the placenta data measured by STARmap and H&E. **b,c**. High and low resolution cell type agreement. **b, and c top**, Placenta portion measured by both ExpSCR (STARmap) and H&E (b, and c top) colored by cell type annotations (legend) on SCHAF-inferred (top) and measured (bottom) data at high (ExpSCR) resolution (b) and at coarsed-up low resolution (c, top). **c, bottom**, Number of cells of each cell type in ExpLR (coarsed-up STARmap) measured (rows) and SCHAF-inferred (column) data for each sample.

**Supplementary Figure 19. Comparison of Unpaired SCHAF inferences to spatially MERFISH cell type annotations in MBC (HTAPP)**

**a.** Spatial cell type accuracy. Proportion of ExpLR (low-resolution (coarsed up) MERFISH) spots (y axis) with cell types that match those on (coarsed up) SCHAF inferences in each sample (x axis) in the MBC (HTAPP) dataset. **b,c**. Comparison of spatial cell annotations in individual sections. **b, and c top**, Tumor sections from different samples (columns) colored by cell type (color legend) from SCHAF inferred (top) and MERFISH measured (bottom) data at cellular resolution (**b,** ExpSCR) or coarsed-up low resolution (**c top,** ExpLR). **c,bottom,** Number of cells of each type in SCHAF inferred (columns) and coarsed-up MERFISH (ExpLR, rows) spatial data for each of the samples in b,c.

**Supplementary Figure 20. Comparison of Unpaired SCHAF spatial expression to STARmap measured genes in the placenta**

**a.** Spatial gene expression accuracy. Average spatial correlation (y axis, left), total number of genes with higher (>0.4) spatial correlation (y axis, middle) and fraction of genes with higher (>0.4) spatial correlation (y axis, right) between ExpLR measurements (coarsed-up, low resolution STARmap) and Unpaired-SCHAF inferred spatial expression in each placenta sample (x axis). **b,c,** Comparison of individual genes. Normalized expression level (color bar) in each placenta sample (major rows, labeled on left) of each of four genes (labeled on top) in SCHAF inferred (top panels) and STARmap measured (bottom panels) data at either single cell resolution (b) or coarsed-up low resolution (c).

**Supplementary Figure 21. Comparison of Unpaired SCHAF spatial expression to MERFISH measured genes in MBC (HTAPP)**

**a.** Spatial gene expression accuracy. Average spatial correlation (y axis, left), total number of genes with higher (>0.4) spatial correlation (y axis, middle) and fraction of genes with higher (>0.4) spatial correlation (y axis, right) between ExpLR measurements (coarsed-up, low resolution MERFISH) and Unpaired-SCHAF inferred spatial expression in each MBC (HTAPP) sample (x axis). **b,c,** Comparison of individual genes. Gene expression level (color bar) in each MBC sample (major rows, labeled on right) of each of four genes (labeled on top) in MERFISH measured (top panels) or SCHAF inferred (bottom panels) data at either single cell resolution (b) or coarsed-up low resolution (c). **d.** Comparison to other inference methods. Mean number (across samples) of coarsed-up, low resolution MERFISH (ExpLR) measured genes with spatial correlation>0.4 (y axis) with each inference method (x axis, color legend). Dots represent individual samples.

## Supplementary Table Legends

**Supplementary Table 1.** The number of cells and genes measured by each modality in each fold/sample.

**Supplementary Table 2.** For each gene in the predicted original mouse pup data, in each fold and the cross validation average, the spatial correlation measurement (when the gene was measured by ExpSCR), predictive quality score, and whether or not the gene was training or hold-out (when the gene was measured by ExpSCR).

**Supplementary Table 3.** For each gene in the predicted new mouse pup data, the spatial correlation measurement (when the gene was measured by ExpSCR) and predictive quality score.

**Supplementary Table 4.** For each gene in the predicted ER+ MBC data, in each fold and in the cross validation average, the spatial correlation measurement (when the gene was measured by ExpSCR), low resolution spatial correlation measurement (when the gene was measured by ExpLR), predictive quality score, and whether or not the gene was training or hold-out (when the gene was measured by ExpSCR).

**Supplementary Table 5.** For each program analyzed in the predicted ER+ MBC data, in each fold and the cross validation average, the low resolution spatial correlation measurement, predictive quality score, and, for the genes in the program that were also measured by ExpLR (Visium), the number of said genes, proportion of said genes from the original program, and the list of genes.

**Supplementary Table 6.** For each gene in the predicted ER-MBC data, the spatial correlation measurement (when the gene was measured by ExpSCR) and predictive quality score, and whether or not the gene was training or hold-out (when the gene was measured by ExpSCR).

**Supplementary Table 7.** For each gene predicted in the HTAPP data, in each sample and in the cross validation average, the Earth Mover’s Distance, Gene Distribution Matching Score, and the spot-level spatial correlation measurement (when the gene was measured by ExpSCR).

**Supplementary Table 8.** For each gene predicted in the Placenta data, in each sample and in the cross validation average, the Earth Mover’s Distance, Gene Distribution Matching Score, and the spot-level spatial correlation measurement (when the gene was measured by ExpSCR in the portion of the tissue measured by ExpSCR)

**Supplementary Table 9.** For each gene predicted in the MSKCC data, in each sample and in the cross validation average, the Earth Mover’s Distance and Gene Distribution Matching Score.

## METHODS

### Human subjects

All samples used in this study were collected under protocols approved by the relevant Institutional Review Boards. All HTAPP MBC samples used in this study underwent IRB review and approval at Dana Farber Cancer Institute (DFCI) (protocol 05-246), as well as at the Broad Institute (protocol #15-370B). All SCLC samples were from patients who underwent a surgical resection or tissue biopsy at Memorial Sloan Kettering Cancer Center (MSKCC), with patients identified and biospecimens collected prospectively from 2005 to 2013 (MSKCC protocol #06-107 A(13)). All patients provided written informed consent through the corresponding Institutional Review Board-approved biospecimen collection and analysis protocol. Tumor specimens were collected intraoperatively, immediately flash-frozen in liquid nitrogen in optimal cutting temperature (OCT) compound, and stored at –80°C until further processing.

### Matched transcriptomic and H&E images data

#### Several different tissue corpora were used

One tissue slice spanning a whole, one day old mouse pup that was profiled by ExpSCR (Xenium) and H&E stained was obtained from a public repository^16^. scRNA-seq from another one day old mouse pup was obtained from a different public repository^34^. One tissue slice spanning all of another one day old mouse pup was assayed by both ISH and H&E in this study (as described below). Cell types of ExpSCR profiles were manually annotated based on k-means (k=10) clustering performed by curators of the publicly available data^16^.

One tissue section from a primary ER+ metastatic breast cancer (MBC) tumor profiled by both Xenium and H&E, scRNA-seq from an adjacent section, and Visium (ExpLR) from another adjacent section as well as one primary ER-MBC tumor section profiled by both Xenium and H&E were all obtained from a single public repository^16^.

Twenty-four (24) small cell lung cancer (SCLC) samples profiled by snRNA-seq and matching H&E stains were obtained from an unpublished study (as described below) from Memorial Sloan Kettering Cancer Center (MSKCC). The snRNA-seq data came from a portion of the same specimen used in H&E staining. Cell types of snRNA-seq profiles were manually annotated by the curators of the data.

Nine other MBC tissue samples, with H&E and snRNA-seq/scRNA-seq and MERFISH data, were obtained from the Human Tumor Atlas Project Pilot (HTAPP)^39^. The H&E, sc/sn-RNA-seq and MERFISH data of each patient sample were obtained from different biopsies of the same metastasis. Cell types of scRNA-seq profiles were manually annotated. Coarse annotations were manually performed on H&E images by an expert histopathologist. Each scRNA-seq profile had two annotations: a specific cell type annotation, and a coarser one with the histopathology annotation categories. Each MERFISH profile had a specific cell type annotation.

Four human placenta samples profiled by scRNA-seq, STARmap, and matching H&E stains were obtained from a study from the Massachusetts General Hospital^40^. scRNA-seq was collected from a section adjacent to that used in H&E staining. STARmap data came from a portion of another section adjacent to the H&E section. Cell types of scRNA-seq profiles were manually annotated by the curators of the data^40^. The H&E stains were collected for the purpose of this study, and precise annotations were manually performed on H&E images by an expert placental histopathologist (as described below). Each scRNA-seq profile had two annotations: a specific cell type annotation, and a coarser one to match up with one of the histopathology annotations. Each STARmap profile had a specific cell type annotation.

Further details on these data sets are in **Supplementary Tables 1-9**.

### New mouse pup ISH data for this study

#### Sample preparation

Glasses were coated with poly(d-lysine) solution. An OCT-embedded P1 mouse (C57BL/6J) was cut into 10-μm, sections then fixed with 4% paraformaldehyde in PBS for 15 min and permeabilized by prechilled methanol at −20 °C for 20 min before hybridization.

#### Library construction

STARmap-ISH genes were selected as described in a below section. SNAIL probes were designed and the probe library was constructed according to previous studies^21^. The probes were dissolved in ultrapure RNase-free water and pooled to the final concentration of 10 nM per probe. The probe mixture was heated at 40 °C for 15 min and then equilibrated to 37 °C. Tissue samples were removed from −20 °C tissues, equilibrated to RT and treated with 10 mM Tris, pH 7.5 for 10 min. The samples were then washed by PBS-TR (0.1% Tween-20, 0.1 U μl−1 of SUPERase•In in PBS) and incubated in hybridization buffer (2× saline– sodium citrate (SSC), 10% formamide, 1% Tween-20, 20 mM ribonucleoside vanadyl complex, 0.1 mg ml−1 of yeast transfer RNA, 0.1 U ul−1 of SUPERase•In, SNAIL probes with 5 nM per probe) at 40 °C with gentle shaking for 48 h. The samples were washed by PBS-TR twice for 20 min at 37 °C and then by 4× SSC in PBS-TR for 20 min at 37 °C, following a rinse by PBS-TR at RT. The SNAIL padlock probes annealed to the samples were ligated by incubating with T4 DNA ligation mixture (1:10 dilution of T4 DNA ligase, 0.2 mg ml−1 of BSA, 0.5 U ul−1 of SUPERase•In) for 2 h at RT with gentle agitation, followed by a 5-min wash of PBS-TR twice.

The samples were then incubated in the RCA mixture (1:50 dilution of Phi29 DNA polymerase, 250 μM dNTP, 20 μM 5-(3-aminoallyl)-dUTP, 0.2 mg ml−1 of BSA, 0.2 U μl−1 of SUPERase•In) for 2 h at 30 °C with gentle agitation, followed by a 5-min wash of PBS-TR twice. Subsequently, the samples were treated with 25 mM acrylic acid N-hydroxysuccinimide ester for 2 h at RT with agitation, rinsed by PBS-T (0.1% Tween-20 in PBS) once and incubated with monomer buffer (4% acrylamide, 0.2% bis-acrylamide, 2× SSC in H2O) for 15 min for polymerization pretreatment. The buffer was removed and 30 μl of monomer mixture (0.1% ammonium persulfate, 0.1% tetra-methylethylenediamine in monomer buffer) was directly added to the center of each sample, which was immediately covered by a coverslip (no. 2 coverslip was coated with Gel-Slick Solution according to the manufacturer’s instructions) and allowed to polymerize in ambiance for 1 h. The tissue–gel hybrid was washed with PBS-T twice and cleared by proteinase K digestion mixture (50 mM Tris, pH 7.5, 100 mM NaCl, 1% SDS, 0.2 mg ml−1 of proteinase K in H2O) at 37 °C overnight. On the next day, the samples were treated with a dephosphorylation mixture (1:100 dilution of shrimp alkaline phosphatase, 0.2 mg ml−1 of BSA, 1:10 dilution of CutSmart buffer in H2O) and rinsed with PBS-T.

#### Imaging and sequencing

The 19-nt fluorescent oligo complementary to DNA amplicon was diluted at 100 nM in 1× SSC dissolved in PBS-T and samples were incubated at RT for 60 min, then washed by PBS-T 3× for 5 min each, DAPI staining was performed for cell segmentation. Images were acquired by Leica Stellaris 5 confocal microscope with a 405 diode, white light laser and 20x air objective.

#### In situ hybridization image processing

For STARmap-ISH images, the process was initiated by detecting spots within the raw images and generating multiplexed images, encompassing multiple genes within a single tissue. The initial step involved aligning images from two rounds of acquisition using the 2D Fourier transform. Subsequently, a method was employed to identify regions with local maxima, which serve as signal spots for visualization. Specifically, maximum and minimum filters were applied from the *scipy.ndimage.filters* python library to locate potential local maxima and minimal values within neighborhoods sized at 15 pixels. Regions where the difference between local maxima and minimal values exceeded a defined threshold of 100 were identified and marked as local maxima. Connected regions were delineated as a result of this identification process. Subsequently, cell segmentation was performed on DAPI channel images using the pretrained *2D_versatile_fluo* model from StarDist^47^. After segmentation, the centroids and areas of the identified cells were extracted. These centroids served as markers for further refinement using a watershed algorithm. The resulting segmented image was then used to assign decoded transcripts to individual cells and to generate the count matrix used for downstream analysis.

### Staining procedure for new mouse pup and placenta H&E data for this study

For Hematoxylin and Eosin (H&E) staining, slides containing 10 μm sections were loaded onto a Histo-Tek® SL Slide Stainer (Sakura Finetek, USA) and stained with the appropriate H&E staining program. H&E-stained slides were mounted with Richard-Allan Scientific™ Cytoseal™ XYL (Thermo Fisher Scientific 8312-4) using glass coverslips (VWR 48393-059). Slide images were obtained by an Axio Scan Z1 Slide Scanner and analyzed with the ZEN Blue Software (Zeiss, Germany).

### Small cell lung cancer H&E stains and snRNA-seq data for this study

#### Histology (H&E Staining)

Formalin-fixed, paraffin-embedded (FFPE) tissue sections were prepared from a portion of each tumor and stained with hematoxylin and eosin (H&E) using standard protocols at the MSKCC pathology core facility. Stained slides were reviewed by a board-certified pathologist to confirm tumor content and histological features.

#### Sample Processing and snRNA-seq Library Preparation

Frozen SCLC tumor samples (n = 24) were used for single-nucleus RNA sequencing (snRNA-seq). Nuclei were isolated from each sample using a Dounce homogenizer in an ice-cold lysis buffer, following standard protocols for frozen tissue. The resulting nuclear suspensions were filtered through a 40 μm cell strainer to remove debris. Nuclei were stained with DAPI and sorted by fluorescence-activated cell sorting (FACS) to enrich for intact, singlet nuclei. snRNA-seq libraries were prepared using the Chromium Single Cell 3’ Reagent Kits (10x Genomics), with 23 samples processed using v3 chemistry and one sample using v2 chemistry, according to the manufacturer’s instructions. Approximately 10,000 nuclei per sample were loaded onto the Chromium instrument. Libraries were sequenced on an Illumina platform with a minimum target depth of 50,000 reads per nucleus.

#### snRNA-seq Data Processing

The snRNA-seq data processing pipeline used in this study is similar to another popular approach^48^. Demultiplexed FASTQ files were generated from raw sequencing reads using Cell Ranger mkfastq (v2.0 and v3.0, 10x Genomics). Reads were aligned to the human GRCh38 genome and gene counts were quantified as unique molecular identifiers (UMIs) using Cell Ranger count (v3.1.0, 10x Genomics). For snRNA-seq data, reads mapping to both introns and exons were counted to increase gene detection sensitivity, following a conventional approach^49^. A pre-mRNA reference was built using Cell Ranger mkref (v3.0) and a modified GTF file, as described in the referenced protocol.

Sequencing data were processed using the Cumulus Cellranger workflow on the Terra cloud platform. CellBender was used to remove ambient RNA contamination from each sample. Count matrices were aggregated across all 24 SCLC samples using the Cumulus workflow, which wraps Pegasus for cloud-based computation. Quality control filtering retained nuclei expressing 200– 6,000 genes and with less than 20% of UMIs mapping to mitochondrial genes. Robust genes were selected at a rate of 0.05% (i.e., expressed in at least 3 out of 6,000 cells). Counts were normalized to TP100K and log-transformed. The top 2,000 highly variable genes were selected in a batch-aware manner, and principal component analysis (PCA) was performed using 50 principal components. Data integration was conducted using *harmony-pytorch*, and a k-nearest neighbor (kNN) graph (k = 100) was constructed using hnswlib. Clustering was performed using the Leiden algorithm (resolution = 2.0), and UMAP was used for visualization. Doublets were identified and removed using Pegasus’ doublet detection algorithm. After doublet removal, the analysis steps from highly variable gene selection to UMAP generation were repeated. Differential expression analysis was performed using the Mann-Whitney U test (FDR = 5%) to identify cluster-specific markers, and putative cell types were annotated based on these results and canonical cell type markers. Annotated epithelial cell clusters were further subclustered (Leiden resolution = 1.0) to distinguish tumor cells from AT1, AT2, and secretory cells.

### Placenta expert histopathologist annotations for this study

Annotations were manually performed on H&E images by a physician pathologist with fellowship training in placental and perinatal pathology. Digital scans of the frozen placenta tissue were obtained and annotated by manually “drawing” over the tissue using photo editing software. Annotations were performed by region and groups/clusters of cell types rather than by individual cells, due to the processing artifact inherent to freezing fresh tissue, with each group of cell types being assigned a different color. Cell types were grouped based on tissue processing artifact, image resolution, and correlation with known placental physiologic relevance. Syncytiotrophoblast and villous cytotrophoblast were marked green, extravillous trophoblast (villous growth columns) were marked blue, decidua and maternal fibrin and blood marked brown. Villous parenchyma (including fibroblasts, stromal cells, Hofbauer cells, and villous vessels including endothelial cells, vascular smooth muscle, and fetal red blood cells) were marked orange. Areas inside the villous parenchyma where fetal vessels were identified with a high degree of likelihood (probable or definitive vessel lumen, with or without intraluminal nucleated red blood cells) were marked red. Foci of intravillous hemorrhage (villi filled with fetal red blood cells), likely a physical artifact caused by the clinical procedure of collecting tissue from the patient, were also marked orange. Software was then used to assign each nucleus from the original H&E image to a cell type group based on the overlying color annotation, so that cell-less empty white space was not annotated with a cell identity.

### Transcriptomics data pre-processing

All ExpSCR data and their corresponding H&E images were aligned at single cell resolution with the Xenium Explorer 2 software program. Cell types of ExpSCR profiles in the original datasets were manually annotated based on kmeans (k=10) clustering performed by curators of the publicly available data.

Cell profiles with less than eight genes were removed (except for ISH profiles, where only eight genes were measured), followed by the removal of genes expressed in no cells. Each cell’s counts *x* were transformed by a *log1p* (*f(x) = log(x + 1)*) transformation. Next, within each tissue dataset, all genes shared across all samples in the dataset were identified and retained for further analysis. The above quality-control steps were performed with python’s *scanpy* package^50^ with default parameters except for the following: *sc.pp.filter_cells*, *sc.pp.filter_genes*, and *sc.pp.log1p*.

### Cell segmenting and tiling histology images

Cells in H&E images from sections also profiled by ExpSCR were segmented according to the location of the single cell transcriptomic measurements.

StarDist^47^ was used to segment cells in the H&E images in all samples used in Unpaired SCHAF.

Histology images were tiled into smaller, potentially-overlapping tiles, each tile centered around a segmented cell’s location, with tile size selected via hyperparameter tuning.

### Paired SCHAF training

All machine learning models were built and trained using the python deep learning library *pytorch*^51^.

Given a training dataset of tissue samples, each with a corresponding H&E image, spatial transcriptomics from the same section as the H&E, and sc/snRNA-seq data, Paired SCHAF was trained to infer an entire spatially resolved, single cell resolution, transcriptome-scale dataset from a histology image.

First, the Supervised Spatial Pretraining model was trained to infer a cell’s training genes from its corresponding H&E tile. The model follows an encoder decoder architecture, with the encoder initialized with weights from UNI^32^, a state-of-the-art foundation model for computational pathology, currently trained on the most data, with the largest number of parameters, and best performance^32^. UNI follows a large vision transformer architecture, with an H&E encoder initialized via the python deep learning library *timm*, via the function *timm.create_model(“vit_large_patch16_224”, img_size=224, patch_size=16, init_values=1e-5, num_classes=0, dynamic_img_size=True)*. A 1024-dimensional embedding of each tile was output by the encoder. The decoder consisted of two fully connected layers: one from 1024 to 512 dimensions, and one from 512 dimensions to the number of training genes. After the first linear layer but before the second was a batch normalization layer followed by a ReLU layer.

An expression profile was predicted by resizing the tile to have both length and width 224 pixels, normalizing it to a RGB mean of (0.485, 0.456, 0.406) and an RGB standard deviation of (0.229, 0.224, 0.225), passing the input tile to the encoder, and then the outputs of the encoder to the decoder. Before training, the expression data for training were z-score normalized so each gene had a mean value of 0 and a standard deviation of 1. The encoder decoder model was then trained on all the spatial transcriptomic profiles of all training samples, with a mean-squared-error (MSE) loss, optimized via stochastic gradient descent, using the Adam optimizer^52^, for up to 20 epochs, with a batch size of 64, at a learning rate of .000005.

After training Supervised Spatial Pretraining, and prior to training Paired SCHAF, Tangram was used to map scRNA-seq/snRNA-seq data onto the spatial transcriptomics data with the *tangram* software library from python. To this end, after filtering out cells in both scRNA-seq and spatial transcriptomics data with fewer than 8 genes with non-zero expression (via *scanpy*’s *sc.pp.filter_cells* function with *min_genes*=8), all spatial transcriptomics data and all scRNA-seq data were separately z-scored. Due to memory constraints of *tangram* software, if there were more than 45,000 scRNA-seq profiles in the sample, 45,000 cells were randomly sampled and retained. The intersection of the genes measured by scRNA-seq and the spatial transcriptomics training genes was termed *tangram_genes*. Spatial transcriptomics data were then partitioned into 25,000 cell chunks to accommodate *tangram* memory constraints. The following three functions were run in succession on each chunk: *tg.pp_adatas(sc_rna, st_chunk, genes=tangram_genes)*, *ad_map = tg.map_cells_to_space(sc_rna, st_chunk,mode=“cells”, density_prior=’uniform’,num_epochs=125,), projected_sc = tg.project_genes(ad_map, sc_rna)*, where *sc_rna* was the processed scRNA-seq data and *st_chunk* was a chunk of the spatial transcriptomics data. All *projected_sc* were then concatenated together to represent the final Tangram-mapped scRNA-seq data onto the spatial transcriptomics. Mouse scRNA-seq data were mapped to ExpSCR data, and cancer scRNA-seq data were mapped to ER+ ExpSCR data.

Finally, Paired SCHAF was trained. Paired SCHAF is an encoder decoder model. The encoder had the same architecture as Supervised Spatial Pretraining, with a large vision transformer, where an H&E tile encoder initialized with the python deep learning library *timm*, via the function *timm.create_model(“vit_large_patch16_224”, img_size=224, patch_size=16, init_values=1e-5, num_classes=0, dynamic_img_size=True)*. The encoder was initialized with weights from the encoder at the end of the training of Supervised Spatial Pretraining. The decoder consisted of four fully*-*connected layers in the following order (1) from 1024 to 2048 dimensions, (2) from 2048 to 4096 dimensions, (3) from 4096 to 8192 dimensions, and (4) from 8192 dimensions to the number of genes in the Tangram-mapped expression data. After each of the first three linear layers (but before the linear layer following each one) was a batch normalization layer followed by a ReLU layer.

An expression profile was predicted by resizing the tile to have both length and width 224 pixels, normalizing it to a RGB mean of (0.485, 0.456, 0.406) and an RGB standard deviation of (0.229, 0.224, 0.225), passing the input tile to the encoder, and then the outputs of the encoder to the decoder. Before training, spatial transcriptomics data for training was z-score normalized for each gene to a mean value of 0 and a standard deviation of 1. The encoder decoder model was then trained on all the spatial transcriptomics profiles of all training samples, with a mean-squared-error (MSE) loss, optimized via stochastic gradient descent, using the Adam optimizer, for up to 20 epochs, with a batch size of 64, at a learning rate of .000005.

### Unpaired SCHAF training

All machine learning models were built and trained using the python deep learning library *pytorch*^51^.

For Unpaired SCHAF, a model was trained to learn a single cell’s expression profile from a single histology-image tile, using an adversarial encoder-decoder^53^ based framework for domain translation applied to image-tile and gene-expression domains.

First, histology images were segmented and tiled (as described above) and the foundation model UNI, as described above, was used to create representations of every histology image tile. The scRNA-seq foundation model scGPT was used to create representations of every scRNA-seq/snRNA-seq profile. scGPT was chosen at the time of the study as the largest in the number of parameters and training data, and a top performer^33^. The resulting H&E embeddings represented each tile as a 1,024-dimensional vector and each scRNA-seq profile as a 512-dimensional vector.

An encoder-decoder, *G*, was trained to reconstruct an scRNA-seq profile on all the expression profiles of all training samples, with a mean-squared-error (MSE) loss, optimized via stochastic gradient descent, for up to 15 epochs, with a batch size of 128, at a learning rate of .0001, with the Adam optimizer. Specifically, let *L* be the latent space of scRNA-seq encodings. If cell type annotations were available, a cell type discriminator, *C,* was trained on all encodings by *G* of all training scRNA-seq profiles. *C* was trained to map an scRNA-seq profile’s encoding to its cell type annotation. Lastly, an adversarial autoencoder *T* was trained on all image tiles of all training samples’ histology tiles, to simultaneously reconstruct image tile, encode to a latent space indistinguishable from *L* via an adversarial training regime of the tile network against an adversarial discriminator, and to encode tiles and scRNA-seq profiles with the same cell type annotation to the same part of the shared latent space.

*C* was trained according to a binary-cross-entropy loss with-logits (BCE), optimized via stochastic gradient descent for 15 epochs, with a batch size of 32, at a learning rate of .001. The adversarial discriminator was trained according to a binary-cross-entropy loss with-logits (BCE), optimized via stochastic gradient descent for up to 50 epochs, with a batch size of 128, at a learning rate of .004. *T* was trained according to a regularized mean-squared-error loss, given by the loss function *f(i’, i, z) = MSE(i’, i) + beta*BCE(z, t) + gamma*BCE(y, a)*, where *i* is the input tile’s embedding by UNI*, i’* is the reconstruction of *i* by *T, t* are the labels signifying that a latent space comes from the gene-expression latent space, *z* is the output of the adversarial discriminator on the encoding of *i* by *T, y* is the output of *C* on the encoding of *i* by *T,* and *a* is the cell type annotation corresponding to *i*. *L* was trained with *beta=*.001 and *gamma=*5, optimized via stochastic gradient descent, for up to 50 epochs, with a batch size of 128, at a learning rate of .001. The adversarial discriminator and *L* were trained in adversarial fashion with alternating gradient updates.

*G* and *T* both had an encoder-decoder architecture. *G*’s encoder was of a frozen scGPT model, and the decoder was composed of four sequential [Linear, BatchNorm, ReLU] blocks, followed by one [Linear, ReLU] block. The dimensions of the five linear layers of the decoder were, in order, 512, 2048, 2048, 2048, 2048, and *number_of_genes* neurons, where *number_of_genes* was the number of genes in each dataset being retained for analysis. *T*’s encoder was of a frozen UNI model, followed by four sequential [Linear, BatchNorm, ReLU] blocks, followed by one [Linear, ReLU] block. The dimensions of these five linear layers were, in order, 1024, 1024, 1024, 1024, 512, and 512 neurons. *T*’s decoder was of four sequential [Linear, BatchNorm, ReLU] blocks, followed by one [Linear, ReLU] block. The dimensions of these five linear layers were, in order, 512, 512, 1024, 1024, 1024, and 1024 neurons.

The adversarial discriminator had an architecture consisting of four [Linear, SpectralNorm, ReLU] blocks, followed by one [Linear, SpectralNorm] block. The dimensions of the five linear layers were, in order, 512, 256, 128, 64, 32, and 2 neurons. *C* had an architecture consisting of four [Linear, SpectralNorm, ReLU] blocks, followed by one [Linear, SpectralNorm] block. The dimensions of the five linear layers were, in order, 512, 256, 128, 64, 32, and *num_cell_types* neurons, with *num_cell_types* as the number of cell types common to both H&E and scRNA-seq data. The Linear, BatchNorm, ReLU, and SpectralNorm layers were implemented via pytorch’s *nn.Linear, nn.BatchNorm1D, nn.ReLU,* and *nn.utils.spectral_norm* functions, respectively.

To translate from image-tile to a single cell expression profile, after training *G* and *T*, the tile was first encoded with *T’s* encoder and then decoded with *G*’s decoder. For a full inference pipeline from H&E-stain to single cell dataset, the input histology image was tiled with the same parameters used for training images, and a single cell expression profile was inferred for each tile.

### Train/test dataset splits

In all datasets, for all models, for both Paired and Unpaired SCHAF, training data were partitioned to a training set (90%) and a validation set (10%), and the latter was used for hyperparameter tuning. Hyperparameter tuning was performed in standard fashion: training 10% of the data over a grid search of possible hyperparameters until a metric computed from the validation data was minimized, at which time the corresponding set of hyperparameters would be used to train the final model. For Paired SCHAF, the average gene spatial correlation of the validation data was minimized. For Unpaired SCHAF, the average gene Earth Mover’s Distance was minimized. Ten percent of the training data, in each case, were used as validation data for hyperparameter tuning in each training corpus. Hyperparameter values found via tuning included, but were not limited to, learning rate (swept over a wide range of values from .01 to .0000001), batch size (swept over a wide range of values from 32 to 256), H&E tile radius (swept over a wide range of values from 16 to 1024 pixels), and the number of hidden layers in each model’s decoder (swept from 1 to 5).

### Evaluating SCHAF via cross validation criteria

To evaluate SCHAF, leave-one-out cross validation was used for each data corpus. For each fold/sample *s* in the corpus, SCHAF was trained on all other folds/samples and evaluated on *s*. Each evaluation benchmark was computed for each sample *s*, using the model trained without *s*. A weighted average was computed across all samples for the final result, with each sample weighted by the number of cells in its ground truth transcriptomic dataset. Corresponding error bars represent one standard error measurement above and below the related cross validation result. In histogram settings, histograms were found by finding histograms for each individual fold/sample and constructing a new cross validation histogram, with each bar having the height of the cross validation averaged height of each fold/sample’s corresponding bar, with error bars representing the standard error of the bar height.

### Paired SCHAF evaluation by spatial cell type annotations

To compare the spatial organization of cell type profiles in measured ExpSCR and Paired SCHAF inferred data, for each sample in a dataset, a cell type classifier was first trained on labeled (cell type annotated) profiles and then applied to predict a cell type label for each Paired SCHAF inferred profile. The classifier had a linear neural network architecture, composed of three sequential [Linear, BatchNorm, ReLU] blocks, followed by one [Linear, SoftMax] block. The dimensions of the four linear layers were, in order, *num_genes*, 1024, 256, 64, and *num_cell_types* neurons, where *num_genes* is the number of genes predicted by SCHAF, and *num_cell_types* is the number of different cell types in the dataset. The Linear, BatchNorm, ReLU, and SoftMax layers were implemented via pytorch’s *nn.Linear, nn.BatchNorm1D, nn.ReLU,* and *nn.Softmax* functions, respectively. The model was trained with a standard cross-entropy loss. Input transcriptomic profiles were z-scored, such that each gene had mean zero and standard deviation 1.

Each ExpSCR and Paired SCHAF inferred profile was plotted spatially and colored by cell type. Accuracy was computed as the proportion SCHAF inferred profiles with a given cell type annotation that matched the annotation of the ExpSCR profile.

Expression profile heterogeneity of cells of each type was computed for cells with each cell type annotation in a sample/fold, in measured (ExpSCR) or Paired SCHAF inferred data. In each case, the standard deviation was computed for the min-max normalized vector of the expression of each gene *g*, and then averaged over all genes as the heterogeneity measure.

### Paired SCHAF evaluation by spatial correlation

For each ExpSCR measured gene, the correlation of measured (ExpSCR) and Paired SCHAF inferred values was computed with the *numpy.corrcoef(a, b)* function in the python *numpy* library, where *a* and *b* were vectors of all SCHAF inferred and ExpSCR measured values. For visualization of individual genes, normalized expression/score were computed by z-scoring the expression vector *x* of gene *g* to a mean 0 and standard deviation 1, followed by min-max normalization to minimum of 0 and maximum of 1. The line of best fit was computed between normalized ExpSCR value and normalized SCHAF inferred values with x-coordinates *numpy.unique(ps)* and y-coordinates *numpy.poly1d(numpy.polyfit(ps, ts, 1))(numpy.unique(ps)),* with python’s *numpy* library, where *ps* and *ts* are SCHAF inferred and ExpSCR measured values, respectively.

### Paired SCHAF evaluation by quality scores

To find the quality score of a gene *g*, the standard deviation was computed for the expression vector of *g*, over all inferred profiles. For genes measured by ExpSCR, the correlation between a gene’s quality score and its spatial correlation value was computed with the *numpy* function *numpy.corrcoef*, as well the line of best fit between these two using *numpy.poly1d*.

Low-resolution SCHAF Inferred data were generated as follows. First, the ExpLR spot containing each ExpSCR inferred profile was identified (if available), and all ExpSCR values in one ExpLR spot were aggregated at spot level and normalized. Spatial correlations between low-resolution SCHAF inferred data and ExpLR measured data were computed for each ExpLR-measured gene expression (line of best fit, as above) and visualized spatially.

Similar analysis was performed for each of the gene programs with >90% genes overlapping with genes inferred by SCHAF, comparing program scores at low-resolution between ExpLR and SCHAF inferences. Following previous studies^54^, the expression score of program *p* was the mean of z-scored expression of program genes. Similar to individual genes, the quality score of a program *p* was found as the standard deviation, over all inferred cells, of the expression score of program *p*.

The eight genes measured by ISH in the new pup placenta sample were randomly selected from the genes that were not measured by ExpSCR and that had quality scores that varied highly across folds (from <0.2 to nearly 2.0). A Paired SCHAF model was then trained on the entire In-Sample Mouse data and used to infer the new mouse sample’s ISH.

### Evaluation of overall properties of RNA profiles inferred by Unpaired SCHAF

‘Pseudobulk’ RNA profiles were calculated for each sample analyzed by Unpaired SCHAF, as the mean expression profiles of either measured or Unpaired SCHAF-inferred scRNA-seq profiles, and Pearson’s correlation coefficient between them was calculated.

For each sample analyzed by Unpaired SCHAF, the 100 most highly variable genes were found in the measured scRNA-seq via *scanpy*’s function *scanpy.pp.highly_variable_genes*, and gene-gene correlation matrices were computed between measured and inferred profiles by converting each to a pandas dataframe and then using the .corr function. Each entry in the matrix was then assigned an integral rank by magnitude, and the matrix was converted into a numpy array using the *scipy* python library’s *stats.rankdata* function. Meta correlation was computed as the Pearson’s correlation between corresponding flattened entries of the measured and SCHAF inferred numpy arrays.

Per-cell-type ‘pseudobulk’ RNA-Seq profiles and their correlations were computed as for pseudobulk above, but separately for all cells of each cell type *c*. In addition, a weighted average of Pearson’s r was calculated, weighted by cell type prevalence, as the “cell type pseudobulk correlation score.”

### Unpaired SCHAF evaluation by gene count distributions

For each gene *g,* the Earth Mover’s Distance (Wasserstein Distance) between the distributions of SCHAF inferred and measured scRNA-seq values (over all profiles) was computed with the function *scipy.stats.wasserstein_distance(a, b)* from the *scipy.stats* python library, where *a* and *b* are the vectors of all counts of gene *g* from all SCHAF inferred and measured profiles, respectively. *a* was un-normalized after prediction (multiplied by the standard deviation from training and added with the mean in training). Each distribution was visualized using the *pandas.plot.kde* function from the python *pandas* library, with default parameters except for *bw_method*, which was set to *n**(-.2)*, where *n* was the number of profiles in the distribution.

### Unpaired SCHAF evaluation by distribution matching scores

The Unpaired SCHAF inferred expression vector of gene *g* was un-normalized (multiplied by the standard deviation from training and added with the mean in training), followed by subtracting half of its standard deviation from 1. This value served as a gene’s distribution matching score. For genes measured by scRNA-seq, the correlation between a gene’s distribution matching score and its Earth Mover’s Distance value was found, plotting the line of best fit (found with the *numpy* function *numpy.poly1d*) between the two, in addition to a scatterplot visualizing the two.

### Unpaired SCHAF evaluation by UMAPs

For each dataset *t* analyzed by Unpaired SCHAF, UMAP embeddings were generated from the measured scRNA-Seq by the following series of *scanpy* functions: *scanpy.pp.highly_variable_genes(t, n_top_genes=4096), scanpy.pp.pca(t, use_highly_variable=1), scanpy.pp.neighbors(t), scanpy.tl.umap(t)*. Next, *scanpy.tl.ingest(p, t, embedding_method=’umap’)* was used to find the embedding of the corresponding Unpaired SCHAF data *p* in the same space as the UMAP of the scRNA-seq measured data *t*. UMAPs were then visualized using the *scanpy.pl.umap* function, coloring by cell type, where Unpaired SCHAF inferred profiles were assigned to cell types by the same classifier-based method used to evaluate Paired SCHAF.

UMAP organization was quantified using a local coherence score, defined as the average, over all cells, of the proportion of each cell’s *k*=10 nearest neighbors in UMAP space that are of its type. Cluster separation was quantized using a cluster separation score, defined as the ratio of an inter-cluster to intra-cluster distance scores. Inter-cluster distance score was defined as the average, over all cells, of each cell’s average distance to cells of its type among its k=15 nearest neighbors. The intra-cluster distance is defined as the average, over all cells, of the cell’s average distance to cells of different types among its k=15 nearest neighbors.

### Unpaired SCHAF evaluation by cell type distributions

Cell types were assigned to Unpaired SCHAF inferred profiles using the same classifier approach as for Paired SCHAF.

Jensen-Shannon Divergence between Unpaired SCHAF inferred and measured scRNA-seq distributions for each sample’s cell type were computed using the *scipy.spatial.distance.jennsenshannon(a, b)* function from the *scipy.spatial* python library, where *a* and *b* are the vectors of cell type proportions in SCHAF inferred and measured scRNA-seq data, respectively.

Expression heterogeneity in Unpaired SCHAF inferred and measured scRNA-seq were computed as for Paired SCHAF, but for Unpaired SCHAF inferences, only cells annotated by the classifier for a given cell type were considered as belonging to that cell type.

### Aggregation of evaluation metrics for Unpaired SCHAF

For each dataset, heatmaps were created to show the value of many Unpaired SCHAF evaluation metrics for each sample. Within each dataset, for each metric, the normalized metric value was computed by z-scoring the vector of all samples’ metric values to a mean 0 and standard deviation 1, followed by min-max normalization to minimum of 0 and maximum of 1. This normalized metric value was then shown for each (sample, metric) pair in the dataset, with samples being noted on the x-axis and metrics on the y-axis.

### Comparison of Unpaired SCHAF to expert pathologist annotations, STARmap or MERFISH

To visually compare to expert pathologist annotations, SCHAF profiles were colored by annotations and regions with two annotations were colored such that a random half of the region’s points had one color and the other half had the other.

H&E image analysis was performed to compare cell type assignments to imaging features. For each H&E image and cell, both the mean pixel intensity and the number of cells were computed from the cell’s H&E tile. The number of cells in each tile was found by placing the coordinates of all cells in each fold/sample in a KDTree (from the python *scipy.spatial* library) and querying the tree for all points that would fall in a tile centered around the cell in question.

The Unpaired SCHAF and STARmap (placenta) or MERFISH (HTAPP MBC) measured spatial expression value of marker genes was also compared for marker genes.

### Spatial accuracy of Unpaired SCHAF predictions

Adjacent slices of H&E and STARmap/MERFISH in parts of the Placenta/HTAPP data were manually aligned. When there was no one-to-one correspondence between individual cells (two separate slices), low-resolution spots (∼five cells / spot) were generated for both inferred (Unpaired SCHAF) and measured (STARmap/MERFISH) data, with average expression of the gene profiles of individual cells in the spot. Spots were assigned a cell type as the most common cell type of the individual cells in the spot. When there was one-to-one spot level alignment, the proportion of correctly annotated cells in each sample was quantified with accuracy and confusion matrix measures, and individual gene correlation measurements across all spots were calculated.

### Ablating cell type information in Unpaired SCHAF training

To assess the importance of cell type information in Unpaired SCHAF, versions of Unpaired SCHAF that did not include the annotation classifier *C* were trained, and their inferred spatial cell type maps were compared to ground truth pathologist annotations as described above.

### SCHAF evaluation against benchmark methods

Paired and Unpaired SCHAF were compared to two sets of benchmark methods.

Paired SCHAF was compared to methods that predict expression profiles from H&E in supervised spatial fashion: SPiRiT^35^, ST-Net^31^, DeepPT^36^, and HE2RNA^30^. Each model was implemented according to the specifications in its study’s Methods section, trained in the same cross validation scheme as SCHAF with hyperparameter tuning, and compared by average spatial correlation and the sample/fold cell type accuracy.

Unpaired SCHAF was compared to Cross-modal Autoencoders^55^, CycleGAN^56^, and DistanceGAN^57^. Cross-modal Autoencoders was implemented according to the image-to-omics setting described in its study’s Methods section^55^. The generator architecture for CycleGAN and DistanceGAN mirrored the final inference model used in Cross-modal Autoencoders, with the same discriminator architecture for all benchmarks as used in the Unpaired SCHAF model. Methods were evaluated by the gene Earth Mover’s Distance, fold/sample cell type distribution Jensen-Shannon Divergence, sample bulk RNA-seq correlation, sample meta correlation score, and sample cell type pseudobulk score, all as defined as described above for Unpaired SCHAF.

## Code availability

Code for comprehensive analysis as described here, and figure generation as shown here can be found at https://github.com/ccomiter1/SCHAF.

## Data availability

All unpublished data, in addition to all other publicly available/published data, have been organized into a single repository named “SCHAF-1M” at the Zenodo link listed on https://github.com/ccomiter1/SCHAF. HTAPP data are part of the HTAN-HTAPP data release on Synapse (syn20834712) as referenced in its own study^39^. MSKCC will be formally released in its own study (MC, leader of that study, can be reached at). Until its publication, its data can be obtained from the “SCHAF-1M” database at the Zenodo link listed on https://github.com/ccomiter1/SCHAF. Placenta transcriptomic data can be found in its original publication^40^, and corresponding H&E images and annotations from the “SCHAF-1M” database at the Zenodo link listed on https://github.com/ccomiter1/SCHAF. Publicly available ExpSCR (Xenium), ExpLR (Visium), and H&E data are at https://www.10xgenomics.com/datasets. New mouse pup ExpSCR (ISH) and H&E data can be found in the “SCHAF-1M” databased at the Zenodo link listed on https://github.com/ccomiter1/SCHAF.

## Author Contributions

CC, EDV, and AR conceived and designed the study. CC designed and executed the study with guidance from JS, CMS, and AR. MC, BL, YY, AK, AQV, and CMR mined and provided SCLC tissue data. JJ, MS, JWu, SV, JWa, TRB, AS, DLA, NW, XZ, and JK mined and provided the HTAPP/MBC tissue data. SJR, MT, and KLP provided expert pathological annotations of HTAPP/MBC tissue data. KZ, JQ, RT, FM, SEB, and SD mined and provided both new mouse data and placenta H&E data in addition to helping process publicly-available ExpSCR (Xenium) data. JM provided expert pathologist annotations of placenta data. XC assisted with software engineering and figure preparation. XC, EDV, CMS, KJKK, and JS provided methodological suggestions. JS, CMS, and AR provided feedback and guidance. CC, JS, CMS, and AR wrote the manuscript, with input from all authors.

## Special Acknowledgement

This work is dedicated to the memory of Minh-Thi Nguyen, the best friend of CC, who, at 24 years of age, on the morning of June 21, 2024, following a horrific bicycle collision, tragically lost her life. This work would not have been possible without her years of unparalleled support, encouragement, camaraderie, care, humor, and loving friendship, for which CC is and will be forever grateful.

## Other Acknowledgements

We thank Ania Hupalowska and Leslie Gaffney for help with graphics and members of the Shu, Stultz, and Regev labs for helpful discussions. Work was supported by the Klarman Cell Observatory and Howard Hughes Medical Institute (AR), NIH New Innovator Award (DP2TR004354, JS), NIH Common Fund (UH3CA275687, UG3CA275687, JS), NICHD (R00HD096049), Massachusetts Life Science Center (JS), Burroughs Wellcome Fund (JS), Additional Ventures (JS), and Massachusetts General Hospital (JS). This project was also funded in part by federal funds from NCI, NIH Task Order HHSN261100039 under Contract HHSN261201500003. CC was supported by a Seibel Scholars Fellowship in addition to a MIT Department of Electrical Engineering and Computer Science Fellowship, Teaching Assistantship, and Research Assistantship. CMR is supported by NCI R35 CA263816 and U24 CA213274. JM is supported by NIH training grant 5T32CA193145. JK was supported by an EMBO Long-Term Fellowship (ALTF 738-2017), an HFSP long-term fellowship (LT000452/2019-L) and an EKFS starting grant (2019_A70).

## Extended Data Figure Legends

**Extended Data Figure 1.**
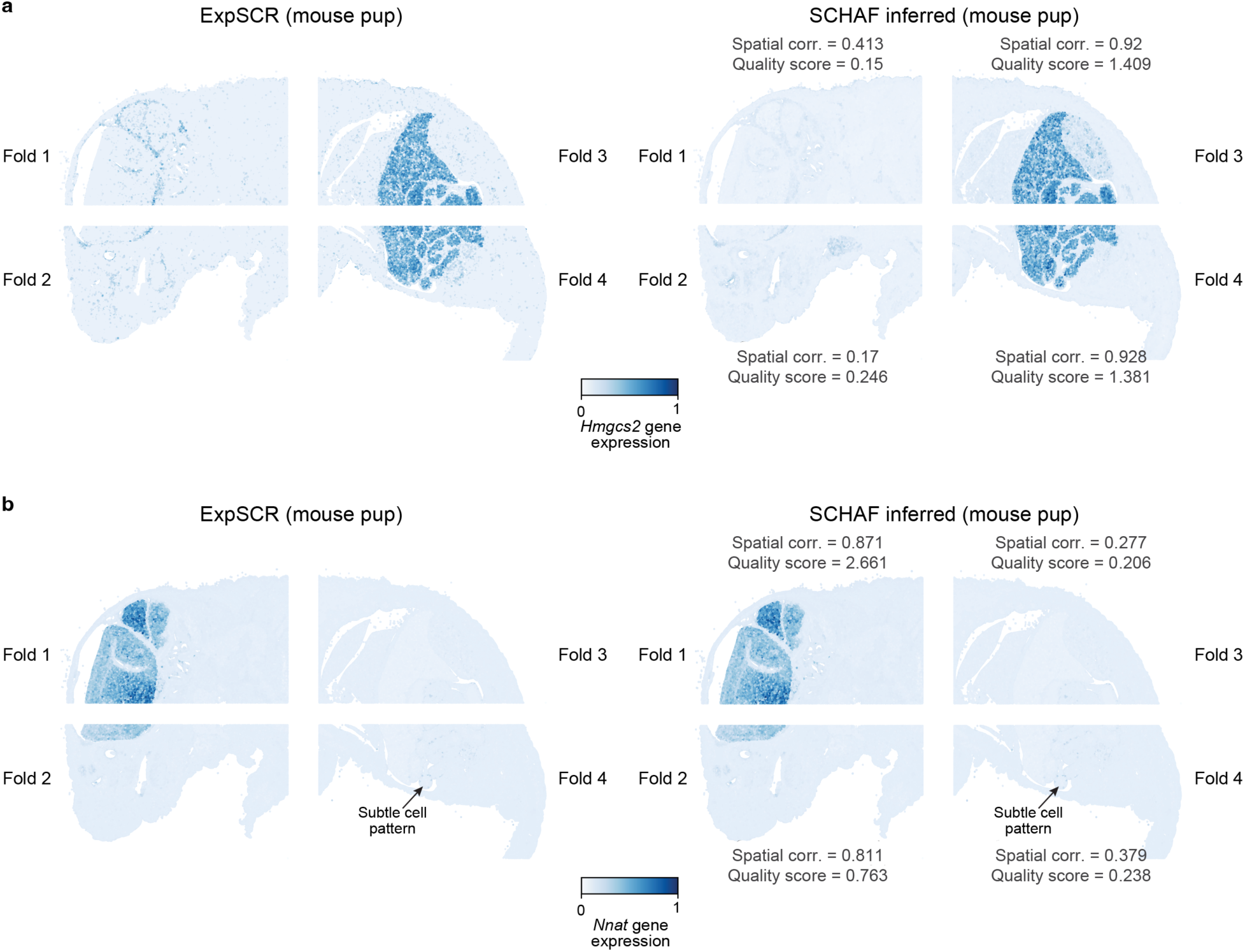
Paired SCHAF predictions of individual gene expression profiles in a healthy mouse pup. Mouse Pup samples, with cells colored by measured (left) or Paired-SCHAF inferred (right) expression of training gene *Hmgcs2* (**a**) or hold-out gene *Nnat* (**b**) in each of four-folds assessed by cross-validation. Quality scores and spatial correlations are noted.

**Extended Data Figure 2.**
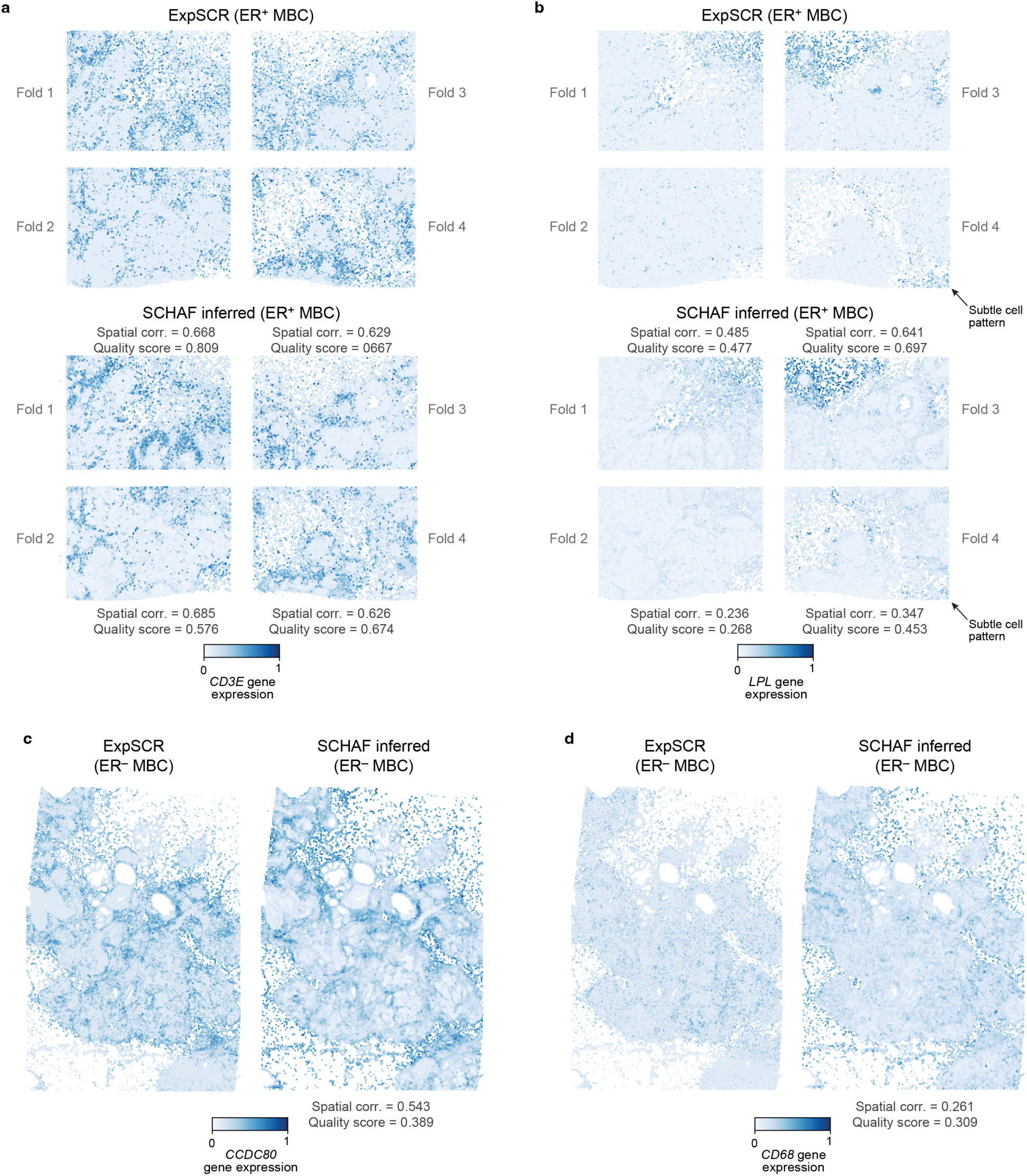
Paired SCHAF predictions of individual gene expression profiles in human metastatic breast cancer. **a,b.** ER+ MBC predictions. ER+ MBC samples with cells colored by measured (ExpSCR, top) or Paired-SCHAF inferred (bottom) expression of training genes *CD3E* (**a**) and *LPL* (**b**) in each of four-folds assessed by cross-validation. Quality scores and spatial correlations are noted. **c-d.** ER-MBC predictions. ER-MBC samples with cells colored by measured (ExpSCR, left) or Paired-SCHAF inferred (right) expression of training genes *CCDC80* (**c**) and *CD68* (**d**). Quality scores and spatial correlations are noted.

**Extended Data Figure 3.**
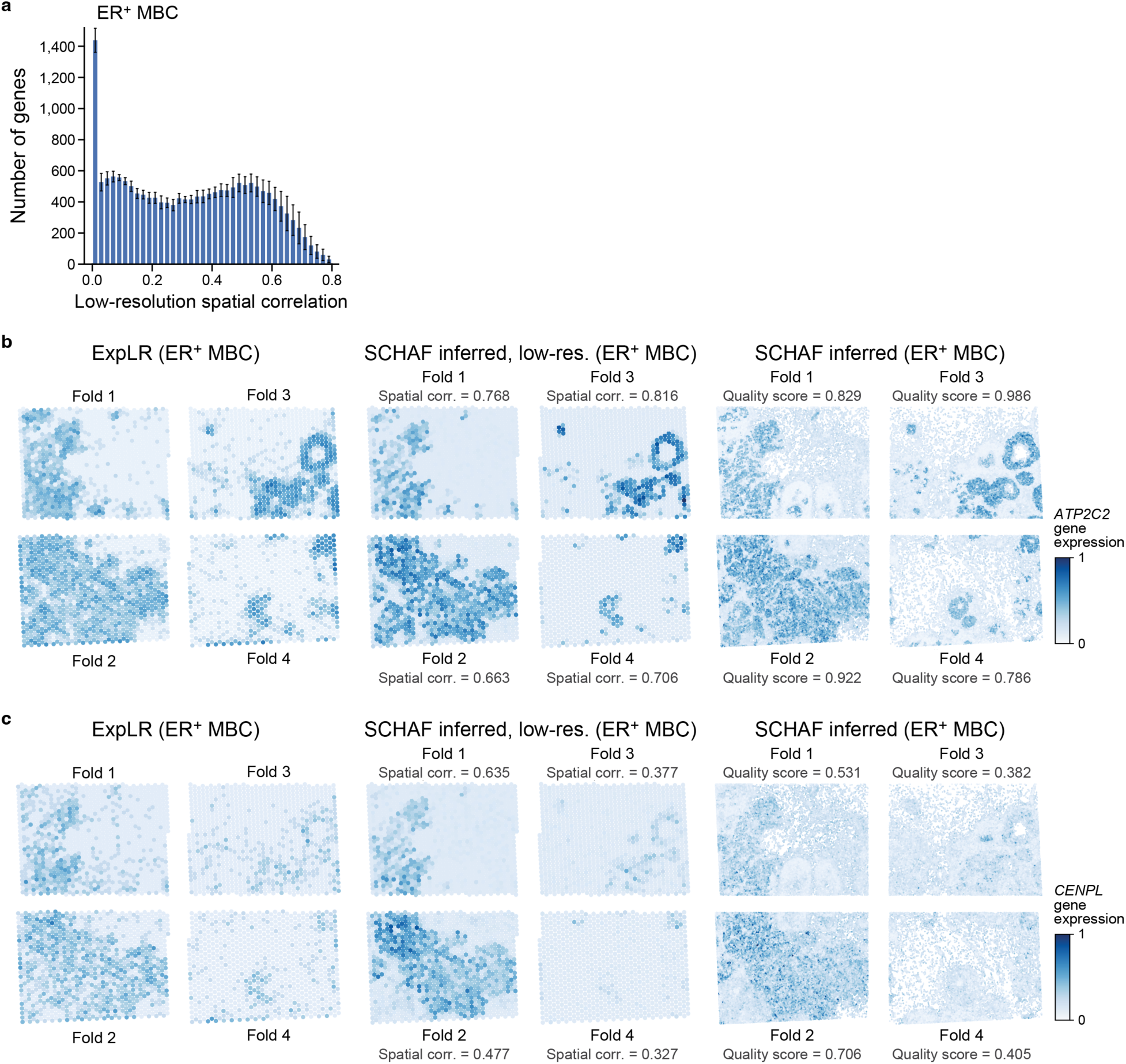
Paired SCHAF whole transcriptome inference in ER+ MBC. **a.** Spatial correlations between measured and SCHAF-inferred expression. Spatial correlations (Pearson’s r, x axis) between low-resolution measured (ExpLR) and the SCHAF inferred (coarsed up) expression values for each measured gene across all ExpLR spots. Error bars: standard error of folds assessed by cross validation. **b-c.** Quality scores capture performance. ER+ MBC samples colored by expression in measured (ExpLR, left, low-resolution spots), coarsed-up SCHAF inferences (middle) and cell-resolution SCHAF inferences (right) for illustrative genes with higher (*ATP2C2*, b) and lower (*CENPL*, c) quality scores (denoted), in each of four folds assessed by cross validation.

**Extended Data Figure 4.**
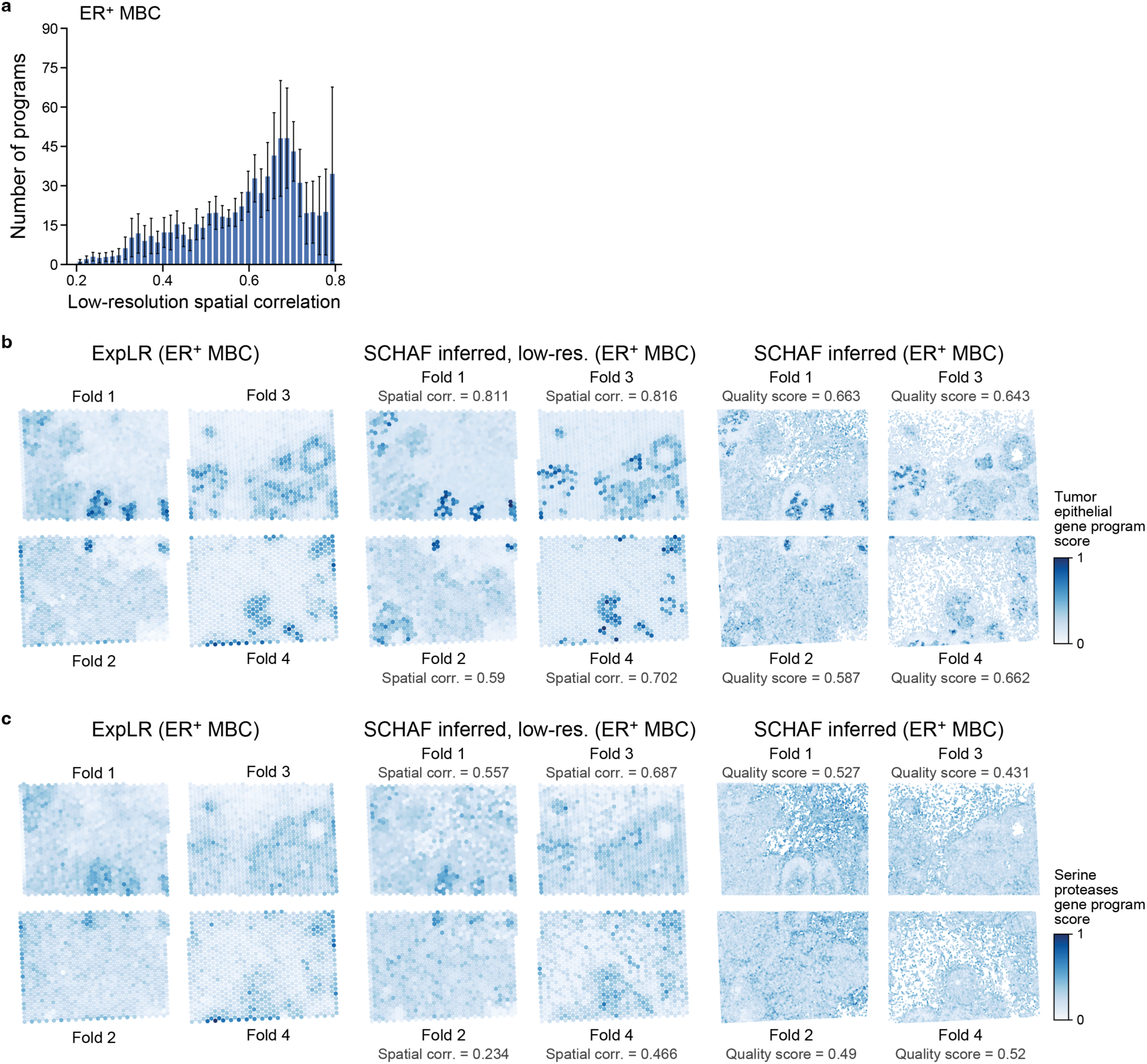
Paired SCHAF gene program inference in ER+ MBC. **a.** Spatial correlations between measured and SCHAF-inferred program scores. Spatial correlations (Pearson’s r, x axis) between low-resolution measured (ExpLR) and the SCHAF inferred (coarsed up) expression scores for each gene program (**Methods**). Error bars: standard error of folds assessed by cross validation. **b-c.** Quality scores capture performance. ER+ MBC samples colored by expression in measured (ExpLR, left, low-resolution spots), coarsed-up SCHAF inferences (middle) and cell-resolution SCHAF inferences (right) for illustrative gene programs with higher (Tumor Epithelial gene program, b) and lower (Serine Proteases gene program, c) quality scores (denoted), in each of four folds assessed by cross validation.

**Extended Data Figure 5.**
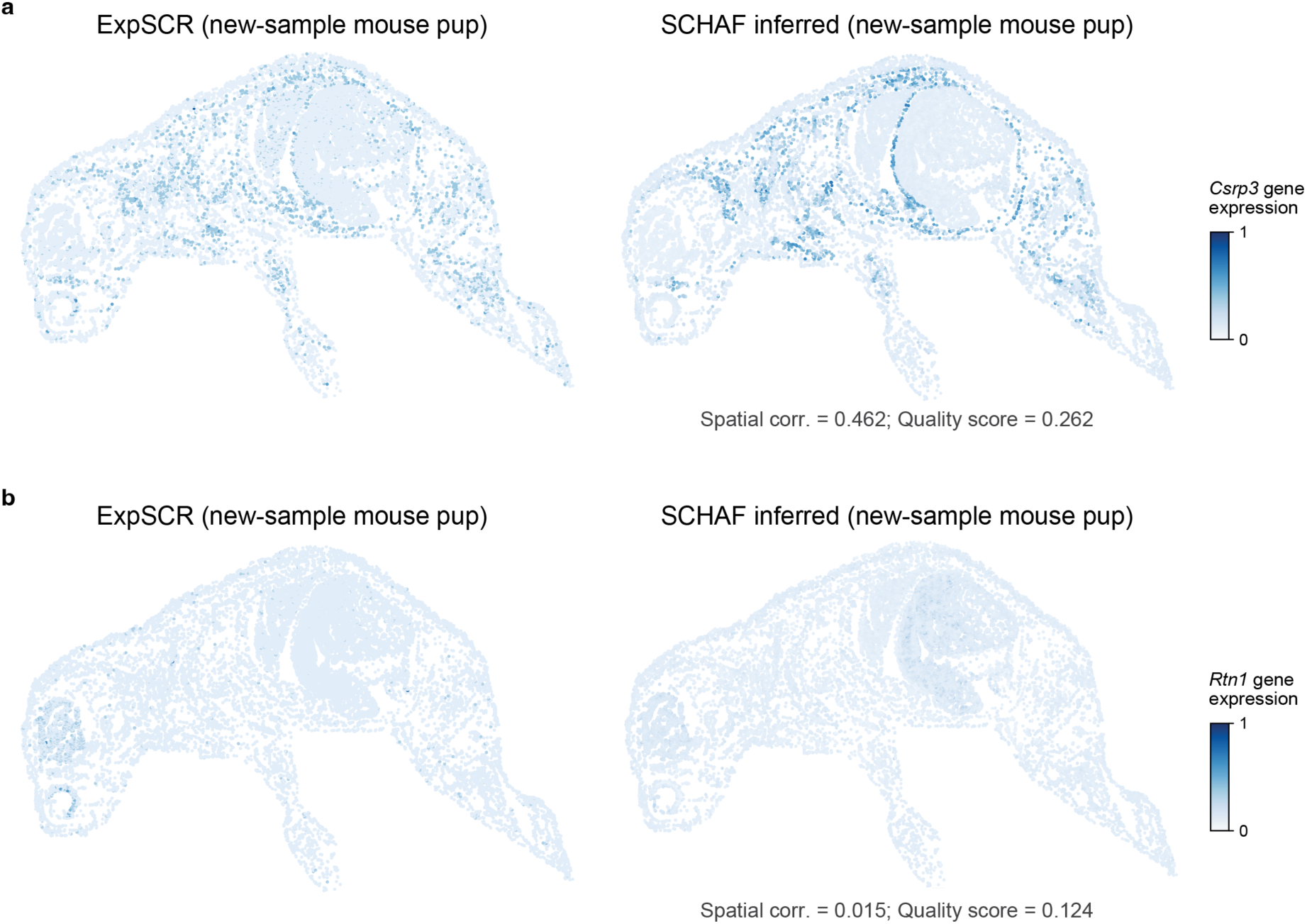
Paired SCHAF whole transcriptome inference in a new mouse pup. New mouse pup samples colored by expression in measured (ExpSCR (ISH), left) and cell-resolution SCHAF inferences (right) for ISH-measured test genes with higher (*Csrp3*, a) and lower (*Rtn1*, b) quality scores (denoted).

**Extended Data Figure 6.**
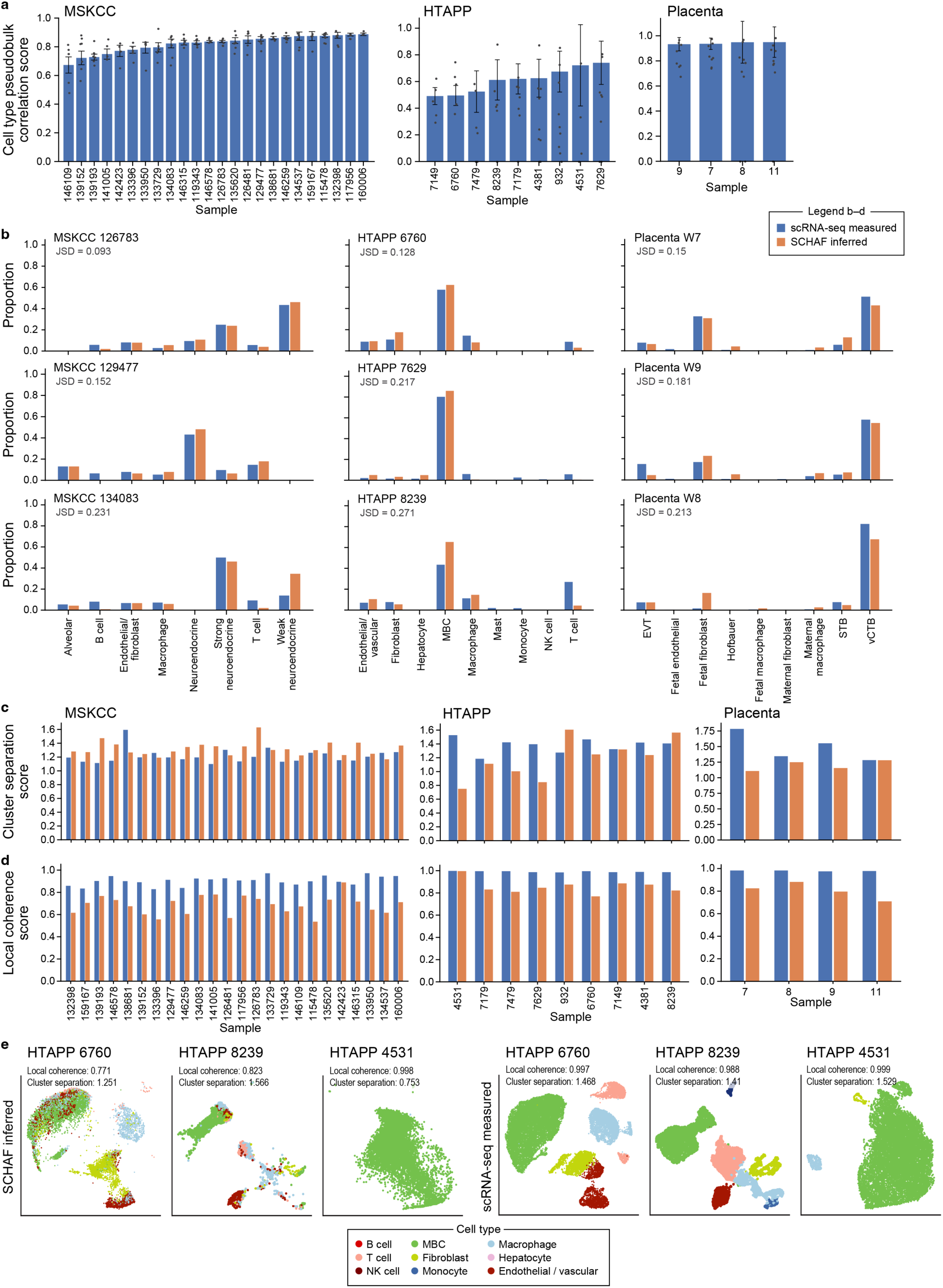
Clustering of unpaired SCHAF inferred profiles resembles measured scRNA-seq. **a.** Jensen-Shannon divergence. Jensen-Shannon Divergence (JSD, x axis) between each sample’s cell type distribution in measured and Unpaired SCHAF-inferred scRNA-seq in SCLC (MSKCC, left), MBC (HTAPP, middle), and placenta (right) samples. **b.** Agreement of cell type proportions between measured and SCHAF-inferred scRNA-Seq. Proportion of cells (y axis) of each type (x axis, as called by a classifier trained on scRNA-seq) in measured (blue) and SCHAF-inferred (orange) scRNA-seq in the SCLC (MSKCC, left), breast cancer (HTAPP, middle) and placenta (right) datasets. **c,d.** Correspondence of clustering metrics. Cluster separation (c, y axis) and local coherence (d, y axis) scores in Uniform Manifold Approximation and Projection (UMAPs) embedding of measured (blue) or Unpaired SCHAF-inferred (orange) scRNA-seq profiles for each sample (x axis, assessed by cross-validation) in the placenta (right), MBC (HTAPP, middle) or SCLC (MSKCC, left) datasets. **e.** Representative UMAPs. UMAP embedding of measured (bottom) or Unpaired SCHAF-inferred (top) scRNA-seq profiles (dots) colored by cell types (legends), for three representative samples from the MBC (HTAPP) dataset. The measured and inferred scRNA-seq are projected in a common space.

**Extended Data Figure 7.**
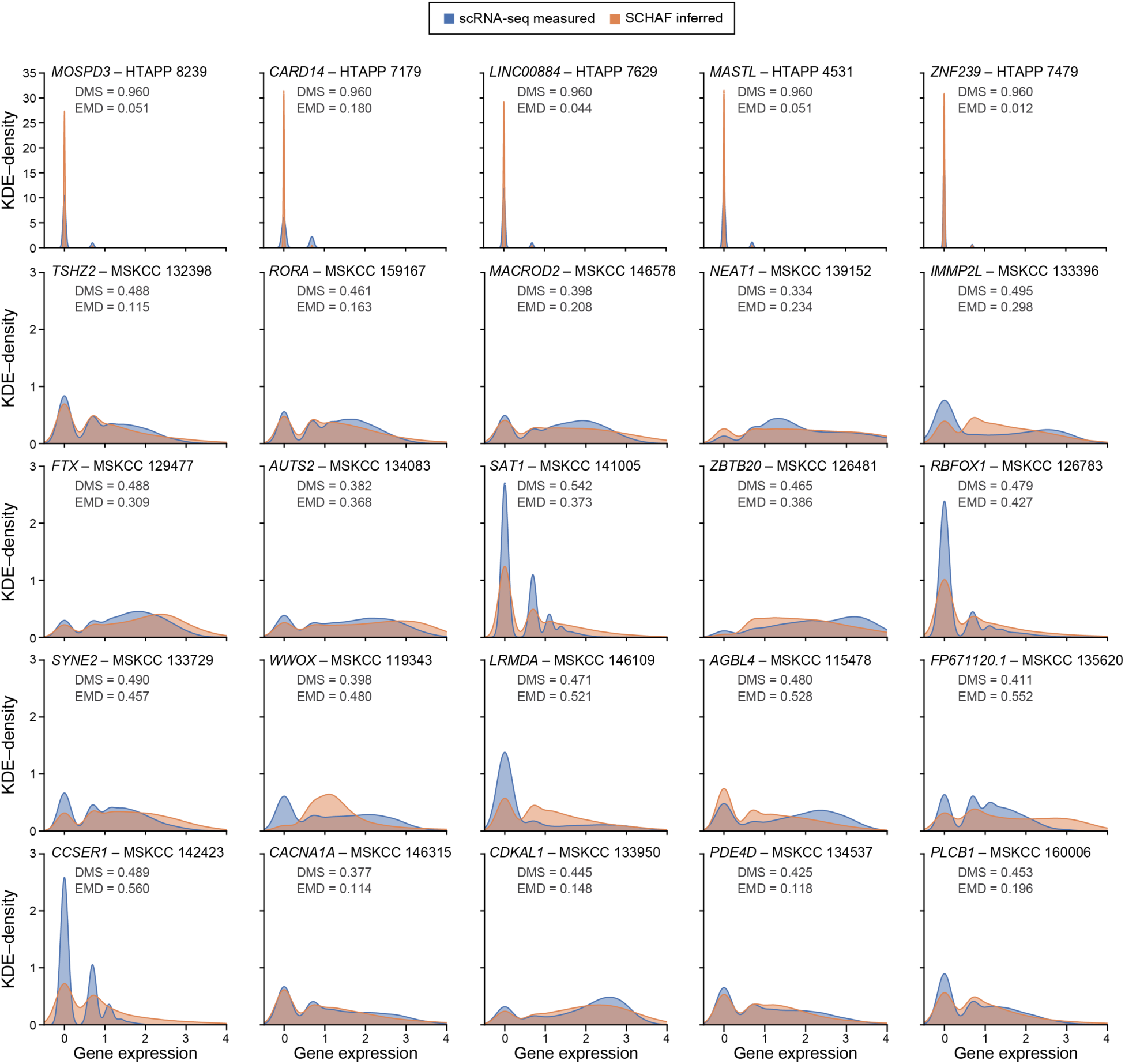
Cell type distributions in Unpaired SCHAF inferences are similar to measured scRNA-seq. KDE plots for measured (blue) and SCHAF inferred (orange) scRNA-seq gene expression for selected genes (labeled on top) from MBC (HTAPP, top row) or SCLC (MSKCC, other rows). EMD and DMS values are denoted.

