## Supplementary Figures for "Inference of single cell profiles from histology stains with the Single Cell omics from Histology Analysis Framework (SCHAF)"

### Supp. Fig. 1

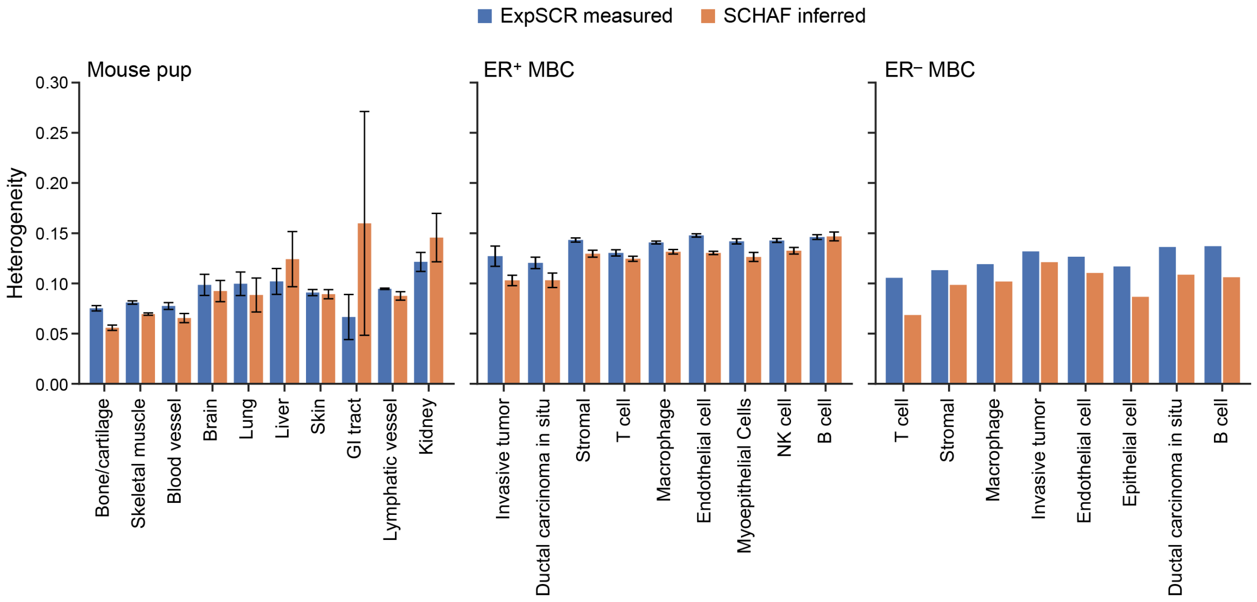

ExpSCR measured
SCHAF inferred
Mouse pup
ER⁺ MBC
ER⁻ MBC
Heterogeneity
0.30
0.25
0.20
0.15
0.10
0.05
0.00
Bone/cartilage
Skeletal muscle
Blood vessel
Brain
Lung
Liver
Skin
GI tract
Lymphatic vessel
Kidney
Invasive tumor
Ductal carcinoma in situ
Stromal
T cell
Macrophage
Endothelial cell
Myoepithelial Cells
NK cell
B cell
T cell
Stromal
Macrophage
Invasive tumor
Endothelial cell
Epithelial cell
Ductal carcinoma in situ
B cell

Supp. Fig. 2

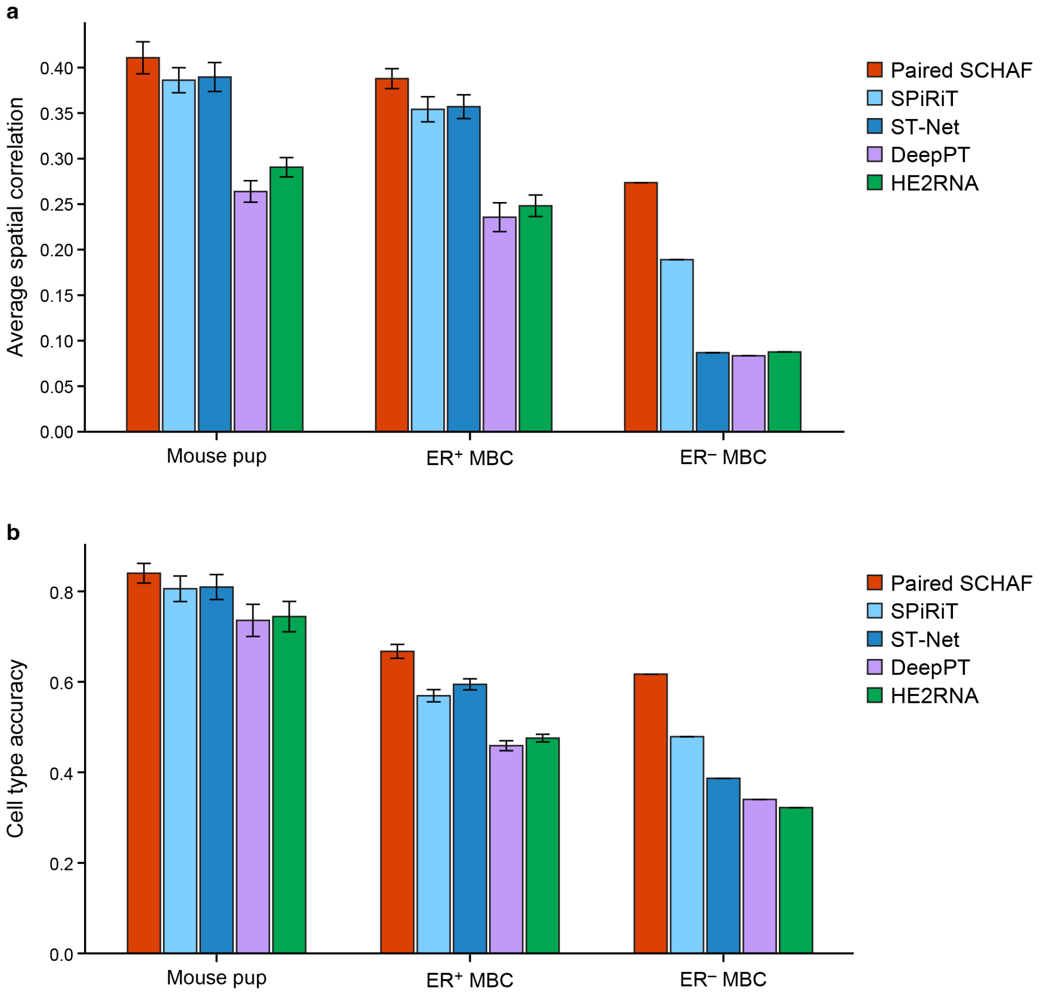
a
Average spatial correlation
0.00
0.05
0.10
0.15
0.20
0.25
0.30
0.35
0.40
Mouse pup
ER+ MBC
ER− MBC
Paired SCHAF
SPiRiT
ST-Net
DeepPT
HE2RNA
b
Cell type accuracy
0.0
0.2
0.4
0.6
0.8
Mouse pup
ER+ MBC
ER− MBC
Paired SCHAF
SPiRiT
ST-Net
DeepPT
HE2RNA

### Supp. Fig. 3

Training:

**Step 1a:**
Train a gene expression decoder to recover scRNA-seq embeddings to the original scRNA-seq gene expression

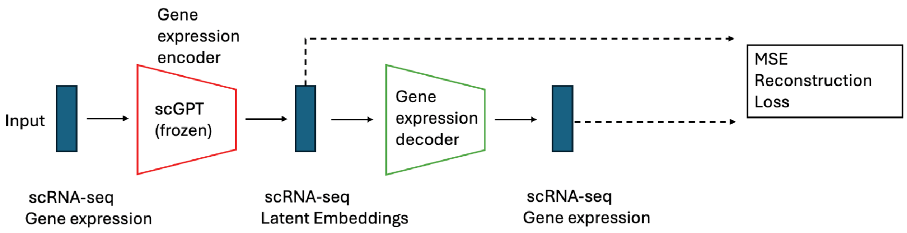


**Step 1b:**
Train a cell type classifier based on scRNA-seq gene expression and its cell type annotation labels
This classifier is used in the **Step 2** to obtain the cell type labels for HE-inferred scRNA-seq

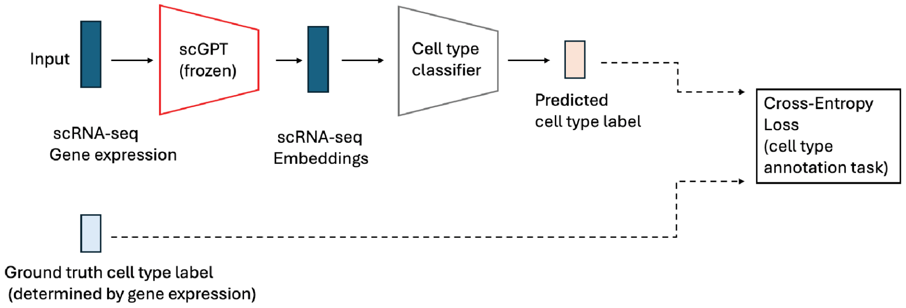


**Step 2:**

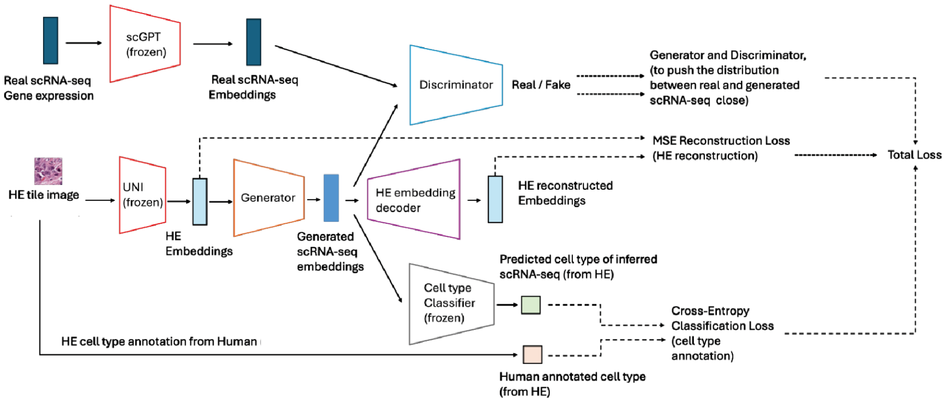


#### Inference:

(1) Input new HE slide, do cell segmentation and crop HE tiles based on cell center coordinates;

(2) Input HE tiles to UNI to get HE embeddings and then sent to Generator to get generated scRNA-seq embeddings, then recover it back to original scRNA-seq gene expression (the the predicted scRNA-seq should have the same cell types as the H&E input)

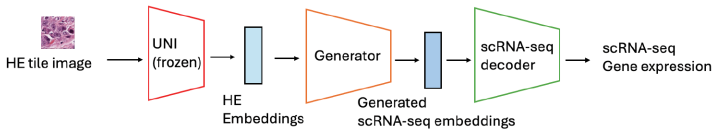

### Supp. Fig. 4

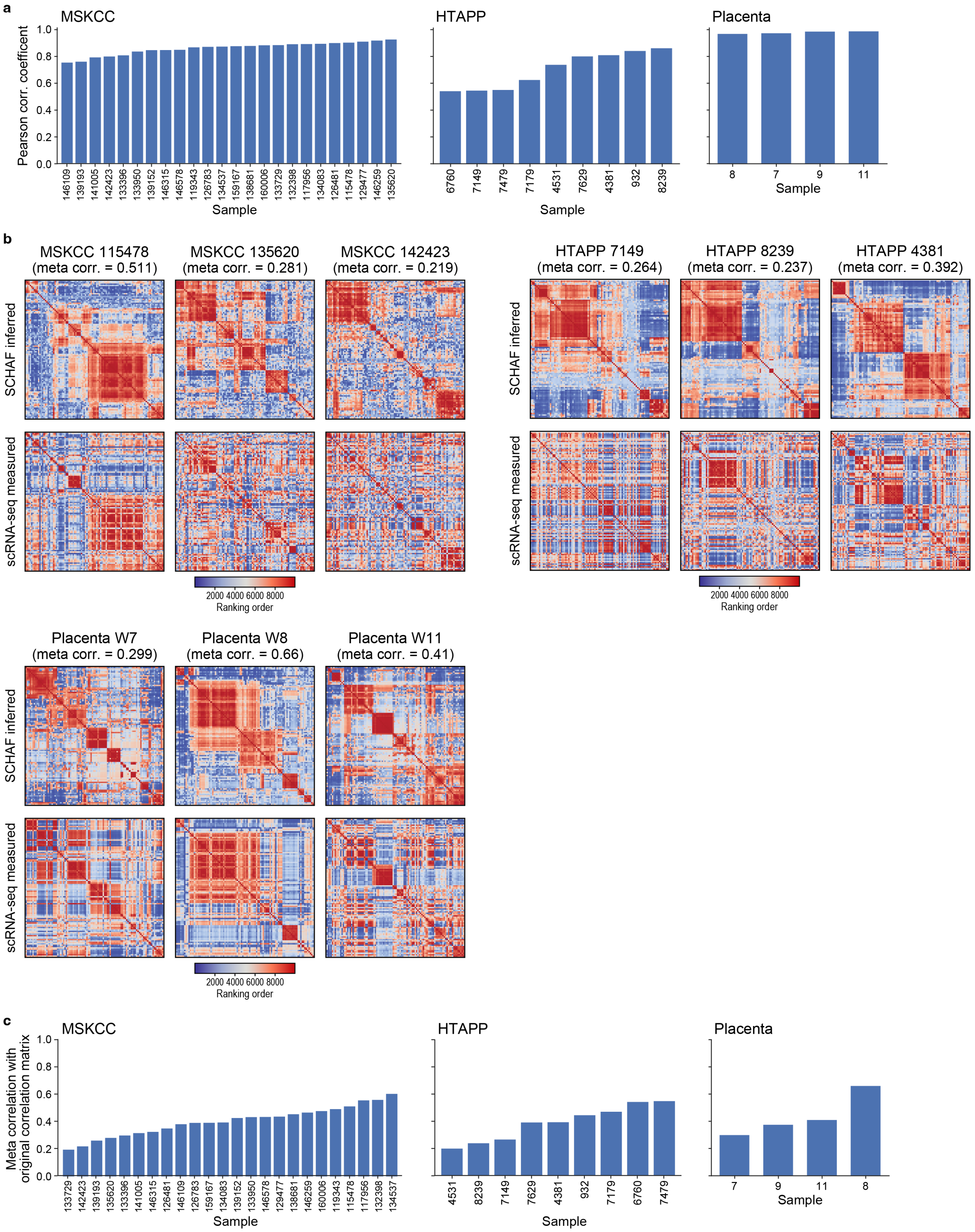
a
MSKCC
HTAPP
Placenta
Pearson corr. coefficent
0.0
0.2
0.4
0.6
0.8
1.0
146109 139193 141005 142423 133396 133950 139152 146315 146578 119343 126783 134537 159167 138681 160006 133729 132398 117956 134083 126481 115478 129477 146259 135620
6760 7149 7479 7179 4531 7629 4381 932 8239
8 7 9 11
Sample
b
MSKCC 115478
(meta corr. = 0.511)
MSKCC 135620
(meta corr. = 0.281)
MSKCC 142423
(meta corr. = 0.219)
HTAPP 7149
(meta corr. = 0.264)
HTAPP 8239
(meta corr. = 0.237)
HTAPP 4381
(meta corr. = 0.392)
SCHAF inferred
scRNA-seq measured
2000 4000 6000 8000
Ranking order
Placenta W7
(meta corr. = 0.299)
Placenta W8
(meta corr. = 0.66)
Placenta W11
(meta corr. = 0.41)
c
MSKCC
HTAPP
Placenta
Meta correlation with
original correlation matrix
133729 142423 139193 135620 133396 141005 146315 126481 146109 126783 159167 134083 139152 133950 146578 129477 138681 146259 160006 119343 115478 117956 132398 134537
4531 8239 7149 7629 4381 932 7179 6760 7479
7 9 11 8
Sample

Supp. Fig. 5

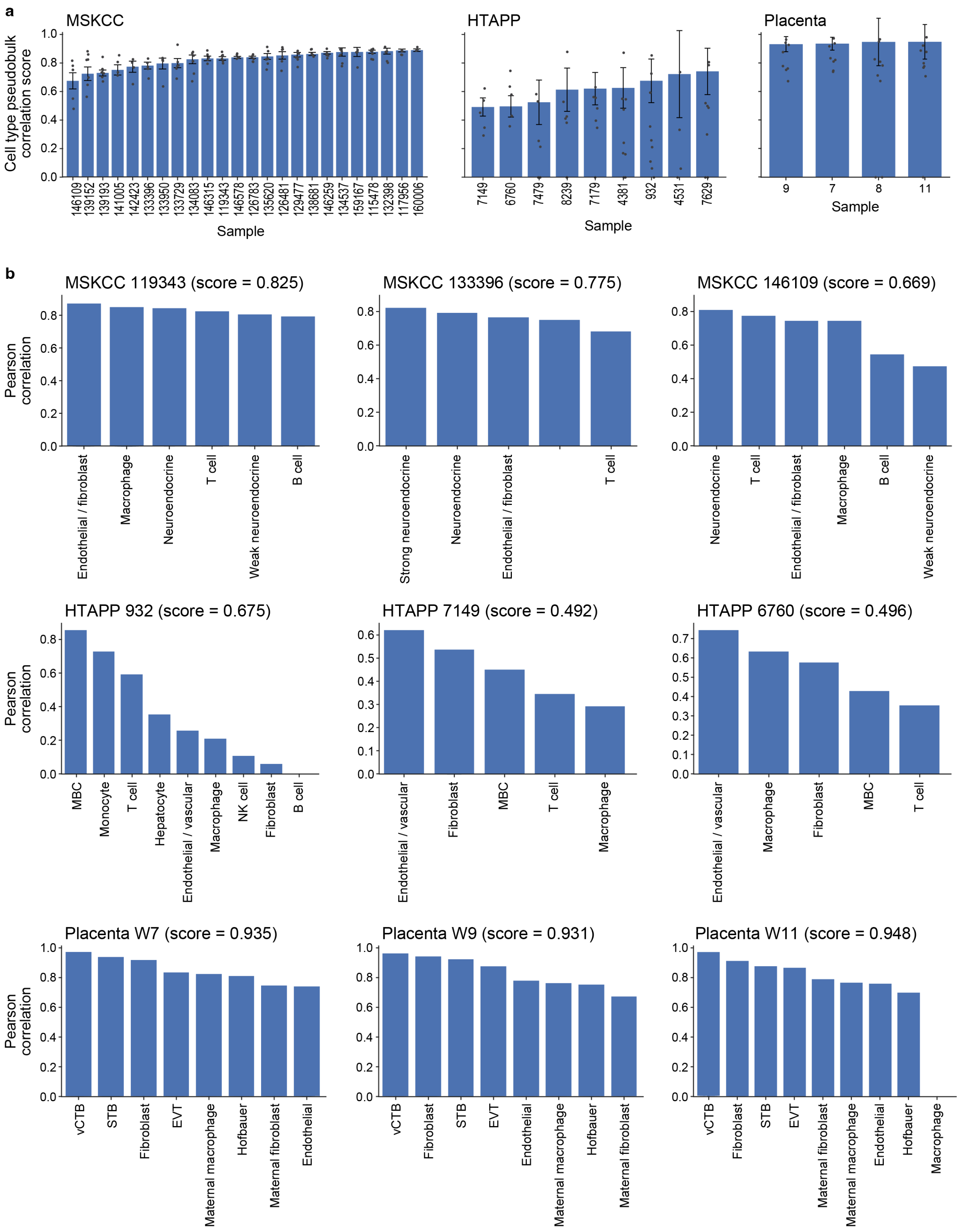
a
MSKCC
HTAPP
Placenta
Cell type pseudobulk correlation score
Sample
146109
139152
139193
141005
142423
133396
133950
133729
134083
146315
119343
146578
126783
135620
126481
129477
138681
146259
134537
159167
115478
132398
117956
160006
7149
6760
7479
8239
7179
4381
932
4531
7629
9
7
8
11
b
MSKCC 119343 (score = 0.825)
MSKCC 133396 (score = 0.775)
MSKCC 146109 (score = 0.669)
Pearson correlation
Endothelial / fibroblast
Macrophage
Neuroendocrine
T cell
Weak neuroendocrine
B cell
Strong neuroendocrine
HTAPP 932 (score = 0.675)
HTAPP 7149 (score = 0.492)
HTAPP 6760 (score = 0.496)
MBC
Monocyte
Hepatocyte
Endothelial / vascular
NK cell
Fibroblast
Placenta W7 (score = 0.935)
Placenta W9 (score = 0.931)
Placenta W11 (score = 0.948)
vCTB
STB
EVT
Maternal macrophage
Hofbauer
Maternal fibroblast
Endothelial
Macrophage

### Supp. Fig. 6

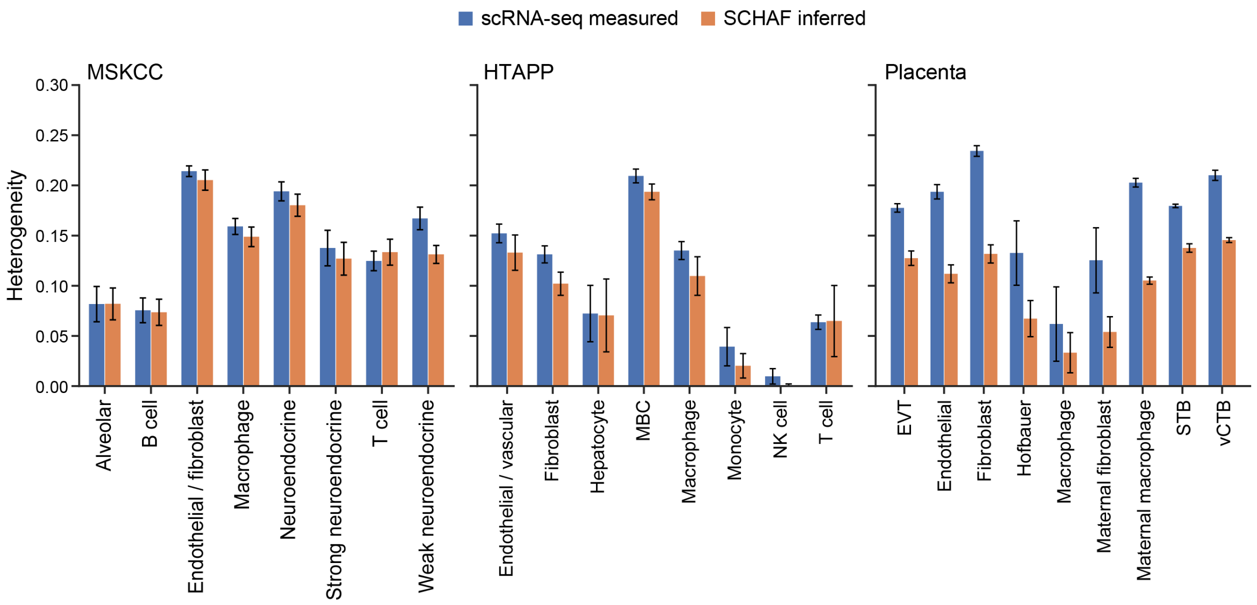

scRNA-seq measured
SCHAF inferred
MSKCC
HTAPP
Placenta
Heterogeneity
0.30
0.25
0.20
0.15
0.10
0.05
0.00
Alveolar
B cell
Endothelial / fibroblast
Macrophage
Neuroendocrine
Strong neuroendocrine
T cell
Weak neuroendocrine
Endothelial / vascular
Fibroblast
Hepatocyte
MBC
Macrophage
Monocyte
NK cell
T cell
EVT
Endothelial
Fibroblast
Hofbauer
Macrophage
Maternal fibroblast
Maternal macrophage
STB
vCTB

### Supp. Fig. 7

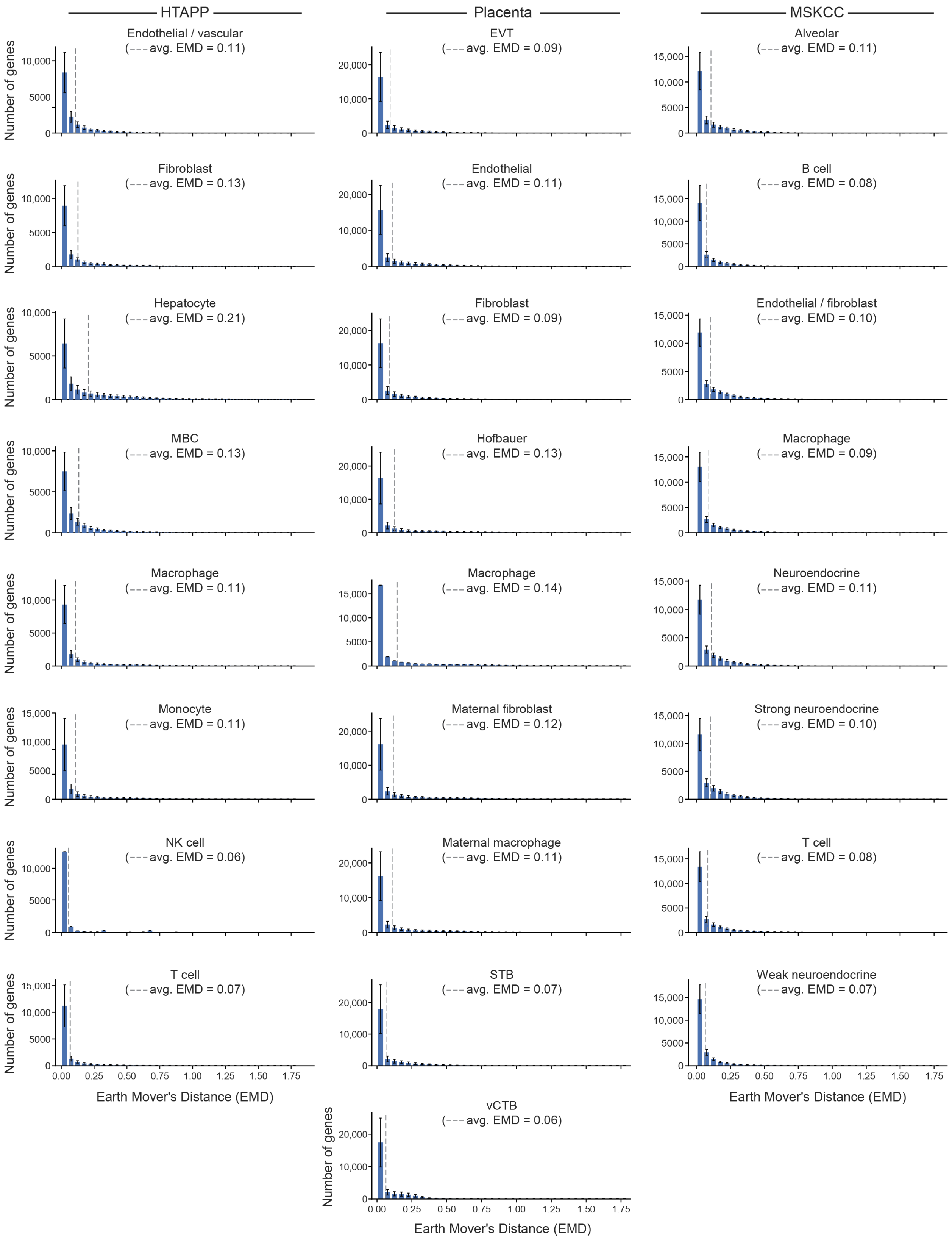
HTAPP
Placenta
MSKCC
Endothelial / vascular
(avg. EMD = 0.11)
EVT
(avg. EMD = 0.09)
Alveolar
(avg. EMD = 0.11)
Fibroblast
(avg. EMD = 0.13)
Endothelial
(avg. EMD = 0.11)
B cell
(avg. EMD = 0.08)
Hepatocyte
(avg. EMD = 0.21)
Fibroblast
(avg. EMD = 0.09)
Endothelial / fibroblast
(avg. EMD = 0.10)
MBC
(avg. EMD = 0.13)
Hofbauer
(avg. EMD = 0.13)
Macrophage
(avg. EMD = 0.09)
Macrophage
(avg. EMD = 0.11)
Macrophage
(avg. EMD = 0.14)
Neuroendocrine
(avg. EMD = 0.11)
Monocyte
(avg. EMD = 0.11)
Maternal fibroblast
(avg. EMD = 0.12)
Strong neuroendocrine
(avg. EMD = 0.10)
NK cell
(avg. EMD = 0.06)
Maternal macrophage
(avg. EMD = 0.11)
T cell
(avg. EMD = 0.08)
T cell
(avg. EMD = 0.07)
STB
(avg. EMD = 0.07)
Weak neuroendocrine
(avg. EMD = 0.07)
vCTB
(avg. EMD = 0.06)
Number of genes
Earth Mover's Distance (EMD)
0.00
0.25
0.50
0.75
1.00
1.25
1.50
1.75

### Supp. Fig. 8

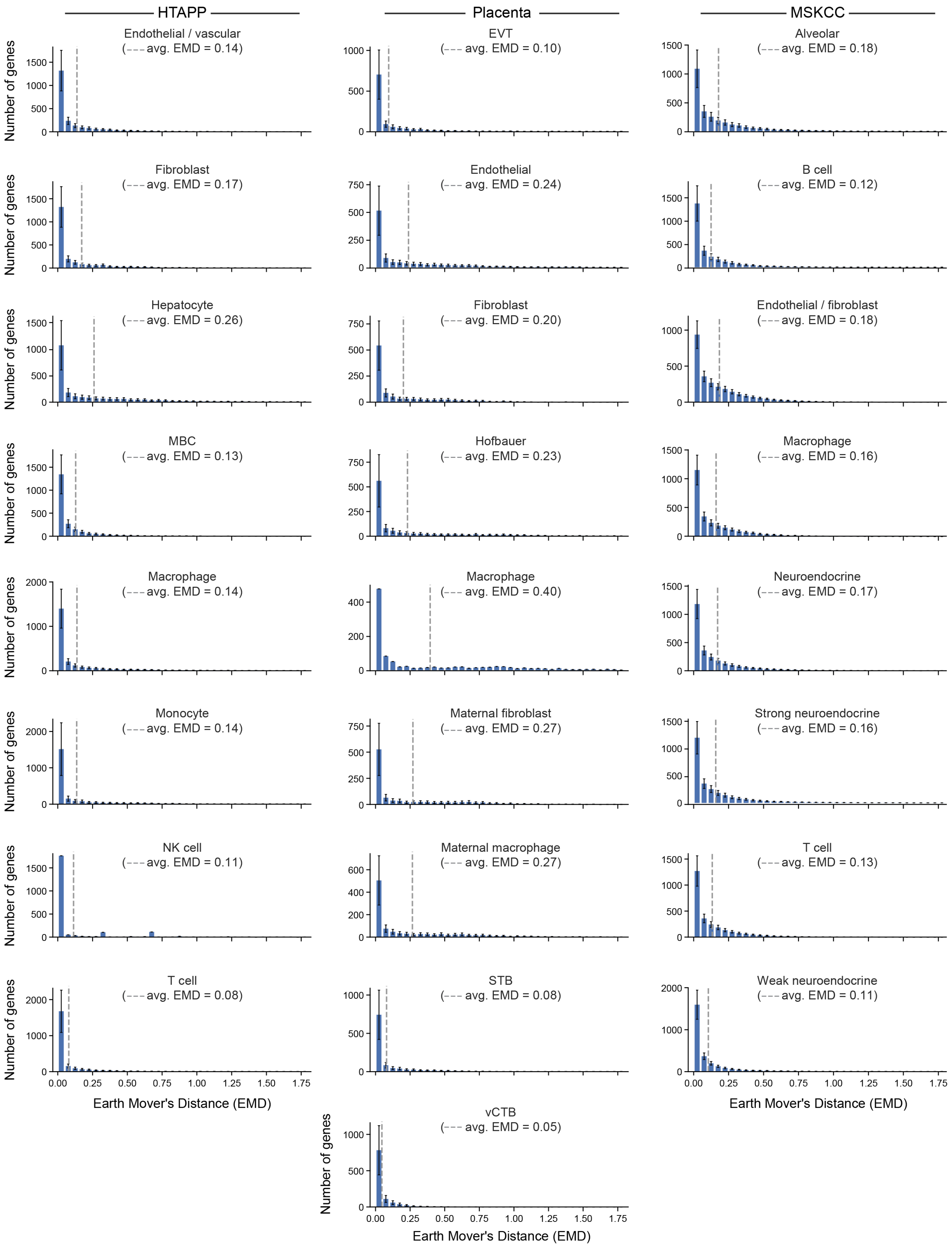
HTAPP
Placenta
MSKCC
Endothelial / vascular (avg. EMD = 0.14)
Fibroblast (avg. EMD = 0.17)
Hepatocyte (avg. EMD = 0.26)
MBC (avg. EMD = 0.13)
Macrophage (avg. EMD = 0.14)
Monocyte (avg. EMD = 0.14)
NK cell (avg. EMD = 0.11)
T cell (avg. EMD = 0.08)
EVT (avg. EMD = 0.10)
Endothelial (avg. EMD = 0.24)
Fibroblast (avg. EMD = 0.20)
Hofbauer (avg. EMD = 0.23)
Macrophage (avg. EMD = 0.40)
Maternal fibroblast (avg. EMD = 0.27)
Maternal macrophage (avg. EMD = 0.27)
STB (avg. EMD = 0.08)
vCTB (avg. EMD = 0.05)
Alveolar (avg. EMD = 0.18)
B cell (avg. EMD = 0.12)
Endothelial / fibroblast (avg. EMD = 0.18)
Macrophage (avg. EMD = 0.16)
Neuroendocrine (avg. EMD = 0.17)
Strong neuroendocrine (avg. EMD = 0.16)
T cell (avg. EMD = 0.13)
Weak neuroendocrine (avg. EMD = 0.11)
Number of genes
Earth Mover's Distance (EMD)
0.00 0.25 0.50 0.75 1.00 1.25 1.50 1.75

Supp. Fig. 9

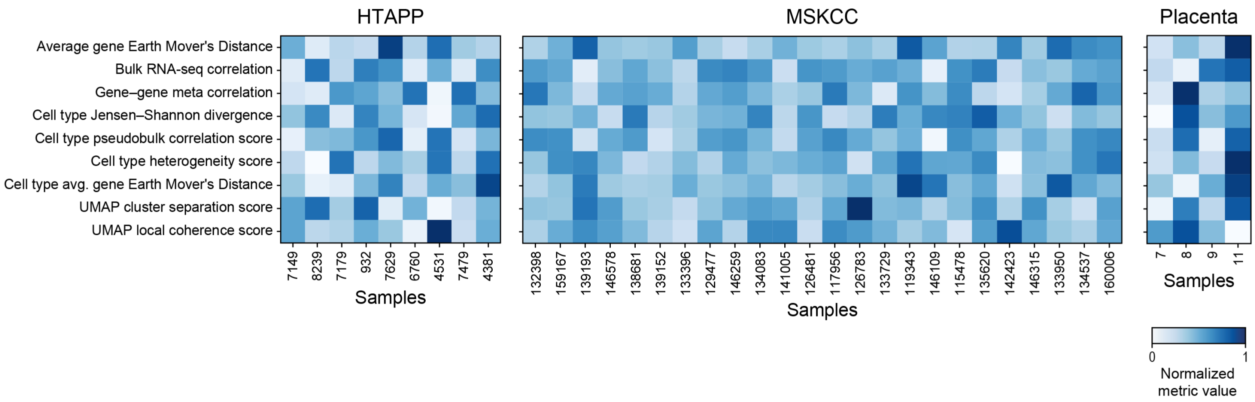
HTAPP
MSKCC
Placenta
Average gene Earth Mover's Distance
Bulk RNA-seq correlation
Gene–gene meta correlation
Cell type Jensen–Shannon divergence
Cell type pseudobulk correlation score
Cell type heterogeneity score
Cell type avg. gene Earth Mover's Distance
UMAP cluster separation score
UMAP local coherence score
7149
8239
7179
932
7629
6760
4531
7479
4381
Samples
132398
159167
139193
146578
138681
139152
133396
129477
146259
134083
141005
126481
117956
126783
133729
119343
146109
115478
135620
142423
146315
133950
134537
160006
Samples
7
8
9
11
Samples
0
1
Normalized
metric value

Supp. Fig. 10

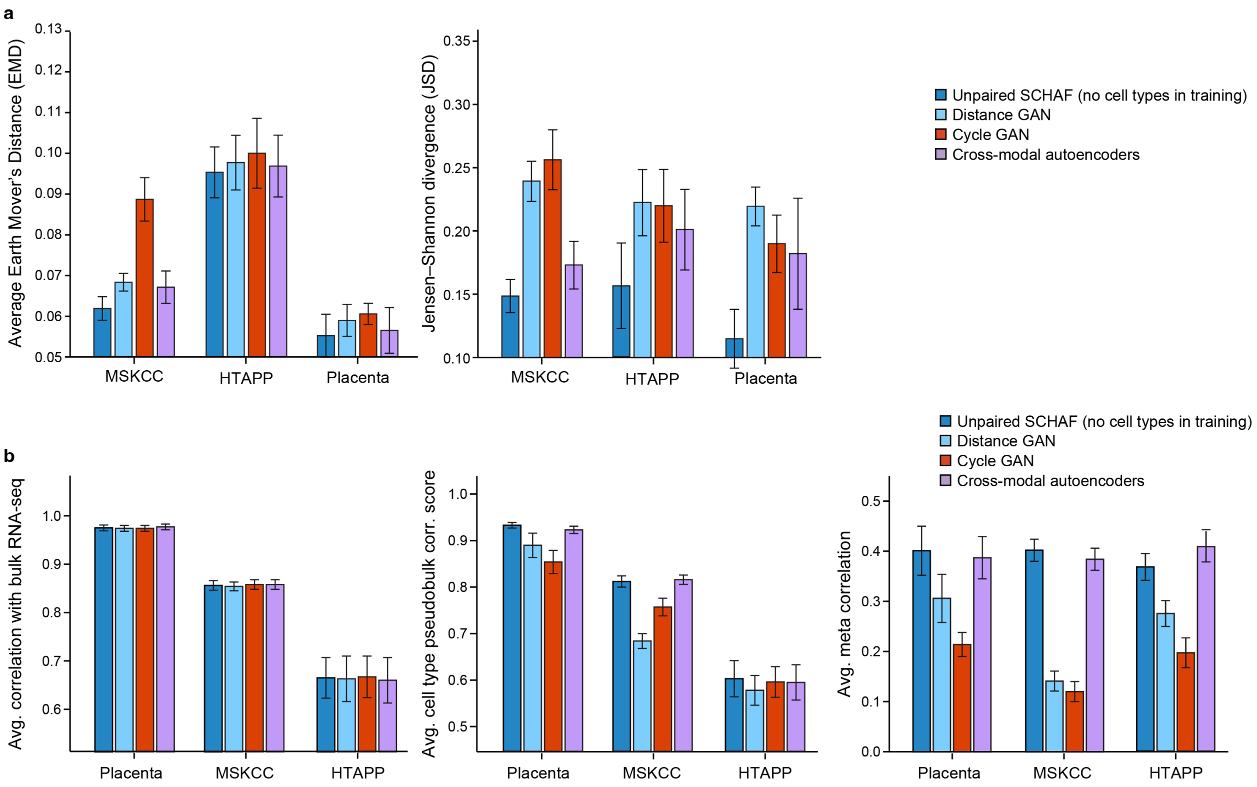
a
Average Earth Mover's Distance (EMD)
MSKCC
HTAPP
Placenta
Jensen–Shannon divergence (JSD)
Unpaired SCHAF (no cell types in training)
Distance GAN
Cycle GAN
Cross-modal autoencoders
b
Avg. correlation with bulk RNA-seq
Avg. cell type pseudobulk corr. score
Avg. meta correlation

### Supp. Fig. 11

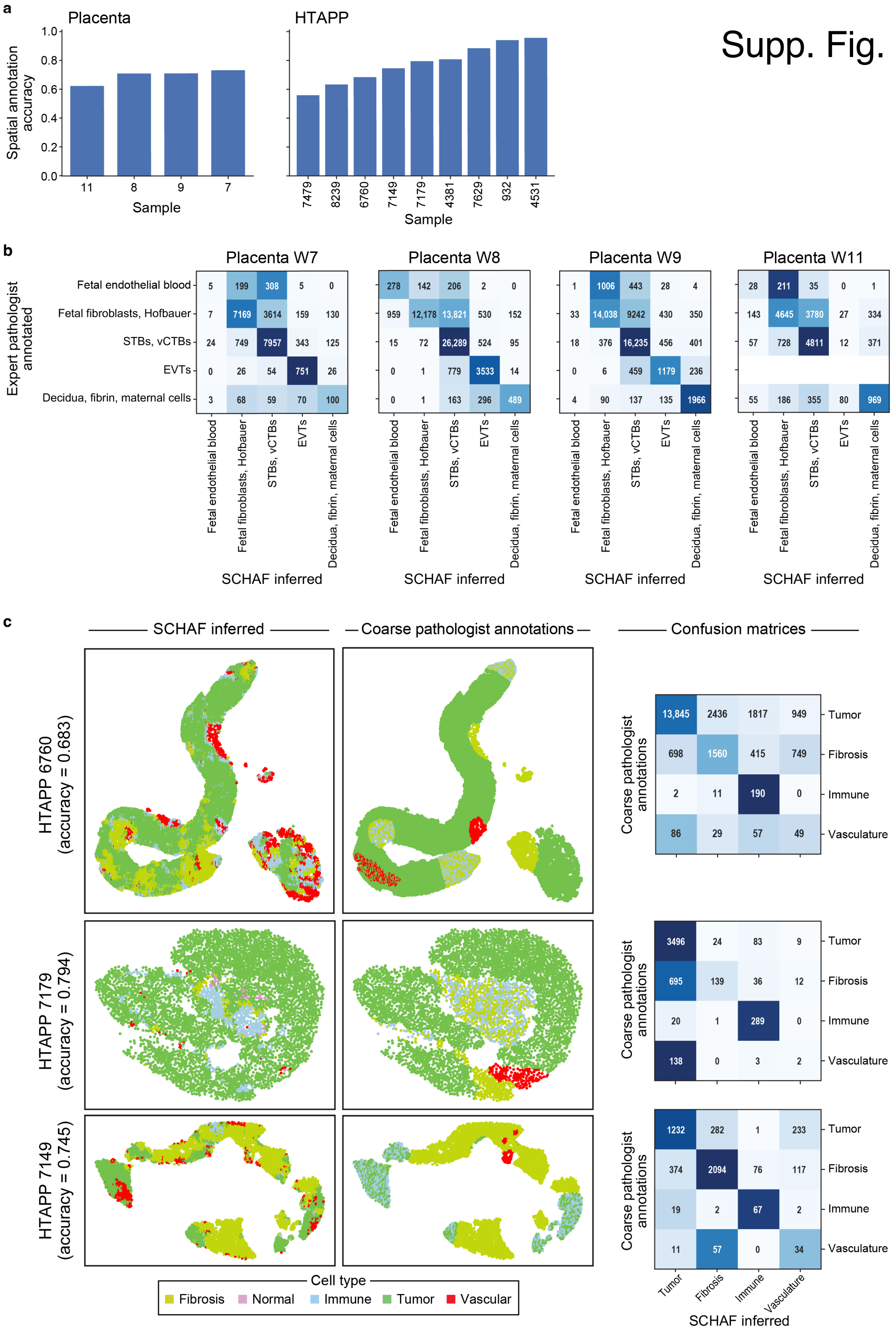

a
Placenta
HTAPP
Spatial annotation accuracy
1.0
0.8
0.6
0.4
0.2
0.0
11
8
9
7
Sample
7479
8239
6760
7149
7179
4381
7629
932
4531
Sample
b
Placenta W7
Placenta W8
Placenta W9
Placenta W11
Expert pathologist annotated
Fetal endothelial blood
Fetal fibroblasts, Hofbauer
STBs, vCTBs
EVTs
Decidua, fibrin, maternal cells
5 199 308 5 0
7 7169 3614 159 130
24 749 7957 343 125
0 26 54 751 26
3 68 59 70 100
278 142 206 2 0
959 12,178 13,821 530 152
15 72 26,289 524 95
0 1 779 3533 14
0 1 163 296 489
1 1006 443 28 4
33 14,038 9242 430 350
18 376 16,235 456 401
0 6 459 1179 236
4 90 137 135 1966
28 211 35 0 1
143 4645 3780 27 334
57 728 4811 12 371
55 186 355 80 969
SCHAF inferred
c
SCHAF inferred
Coarse pathologist annotations
Confusion matrices
HTAPP 6760 (accuracy = 0.683)
HTAPP 7179 (accuracy = 0.794)
HTAPP 7149 (accuracy = 0.745)
Coarse pathologist annotations
13,845 2436 1817 949
698 1560 415 749
2 11 190 0
86 29 57 49
3496 24 83 9
695 139 36 12
20 1 289 0
138 0 3 2
1232 282 1 233
374 2094 76 117
19 2 67 2
11 57 0 34
Tumor
Fibrosis
Immune
Vasculature
SCHAF inferred
Cell type
Fibrosis
Normal
Immune
Tumor
Vascular

Supp. Fig. 12

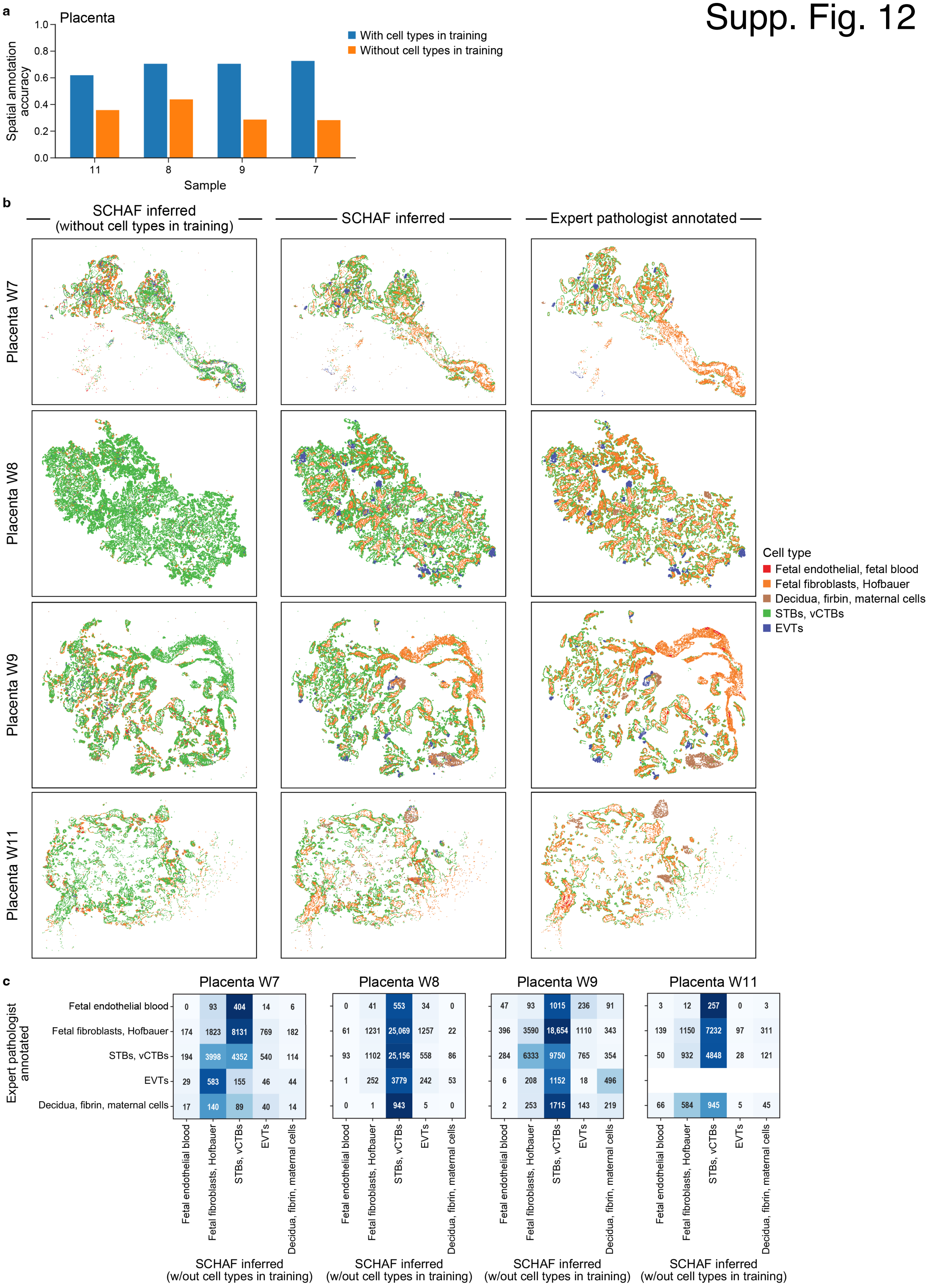
a
Placenta
Spatial annotation accuracy
Sample
With cell types in training
Without cell types in training
b
SCHAF inferred (without cell types in training)
SCHAF inferred
Expert pathologist annotated
Placenta W7
Placenta W8
Placenta W9
Placenta W11
Cell type
Fetal endothelial, fetal blood
Fetal fibroblasts, Hofbauer
Decidua, firbin, maternal cells
STBs, vCTBs
EVTs
c
Placenta W7
Placenta W8
Placenta W9
Placenta W11
Expert pathologist annotated
Fetal endothelial blood
Fetal fibroblasts, Hofbauer
STBs, vCTBs
EVTs
Decidua, fibrin, maternal cells
0 93 404 14 6
174 1823 8131 769 182
194 3998 4352 540 114
29 583 155 46 44
17 140 89 40 14
0 41 553 34 0
61 1231 25,069 1257 22
93 1102 25,156 558 86
1 252 3779 242 53
0 1 943 5 0
47 93 1015 236 91
396 3590 18,654 1110 343
284 6333 9750 765 354
6 208 1152 18 496
2 253 1715 143 219
3 12 257 0 3
139 1150 7232 97 311
50 932 4848 28 121
66 584 945 5 45
SCHAF inferred (w/out cell types in training)

Supp. Fig. 13

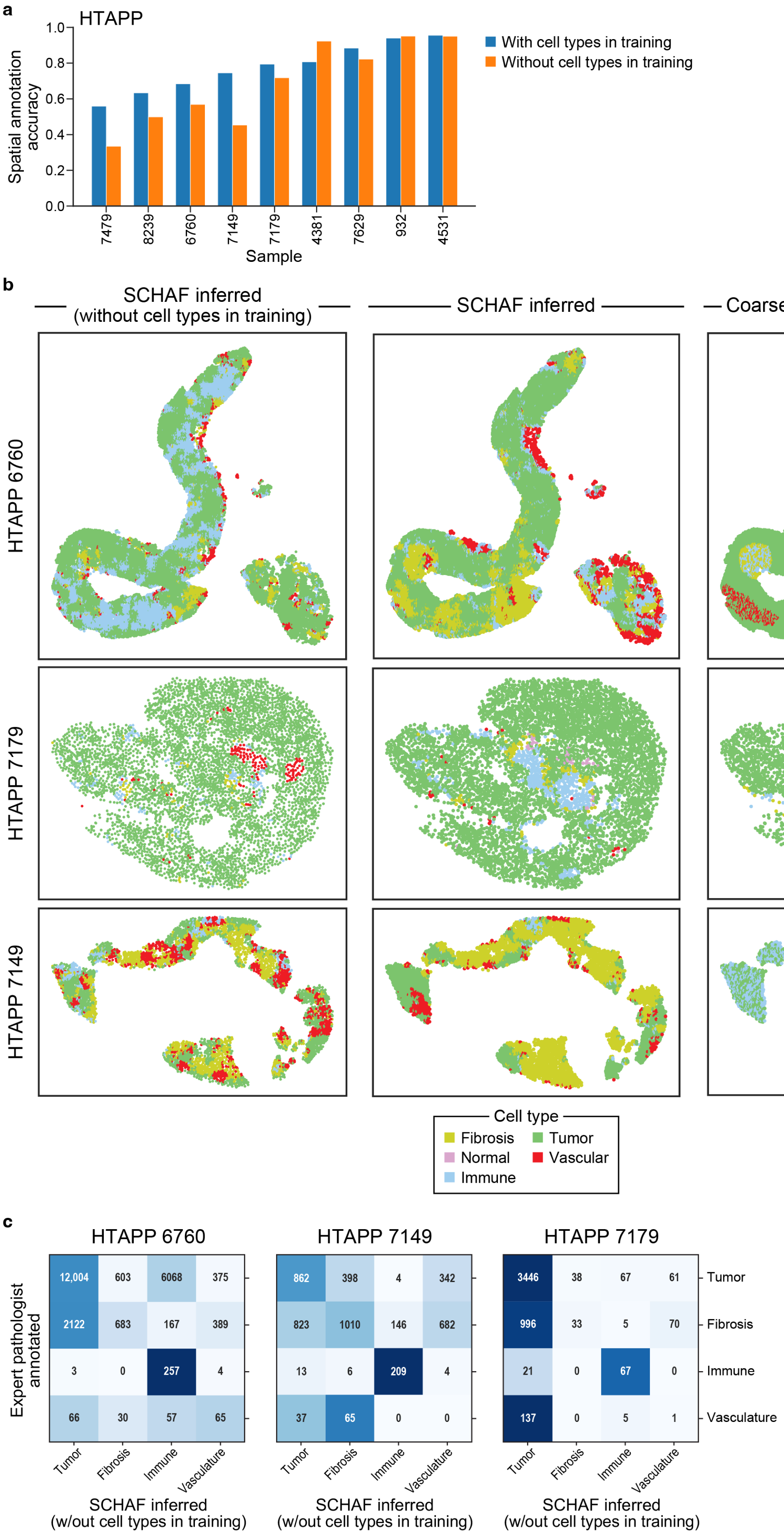

a
HTAPP
Spatial annotation accuracy
1.0
0.8
0.6
0.4
0.2
0.0
With cell types in training
Without cell types in training
7479
8239
6760
7149
7179
4381
7629
932
4531
Sample
b
SCHAF inferred (without cell types in training)
SCHAF inferred
Coarse pathologist annotations
HTAPP 6760
HTAPP 7179
HTAPP 7149
Cell type
Fibrosis
Tumor
Normal
Vascular
Immune
c
HTAPP 6760
HTAPP 7149
HTAPP 7179
Expert pathologist annotated
12,004
603
6068
375
2122
683
167
389
3
0
257
4
66
30
57
65
862
398
4
342
823
1010
146
682
13
6
209
4
37
65
0
0
3446
38
67
61
996
33
5
70
21
0
67
0
137
0
5
1
Tumor
Fibrosis
Immune
Vasculature
SCHAF inferred (w/out cell types in training)

### Supp. Fig. 14

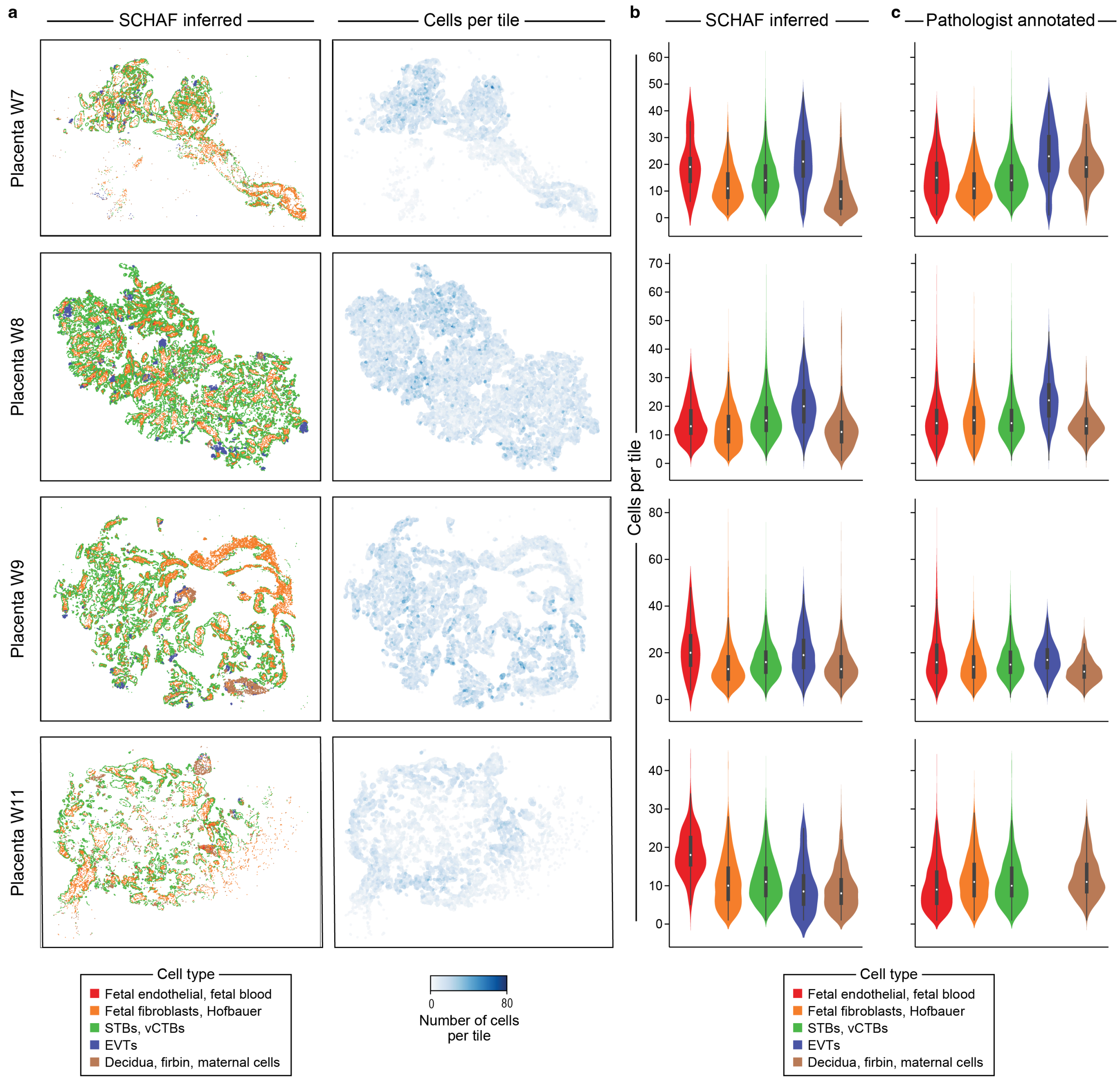

a
SCHAF inferred
Cells per tile
Placenta W7
Placenta W8
Placenta W9
Placenta W11
Cell type
Fetal endothelial, fetal blood
Fetal fibroblasts, Hofbauer
STBs, vCTBs
EVTs
Decidua, firbin, maternal cells
0
80
Number of cells per tile
b
SCHAF inferred
c
Pathologist annotated
Cells per tile
0
10
20
30
40
50
60
70
80
Cell type
Fetal endothelial, fetal blood
Fetal fibroblasts, Hofbauer
STBs, vCTBs
EVTs
Decidua, firbin, maternal cells

### Supp. Fig. 15

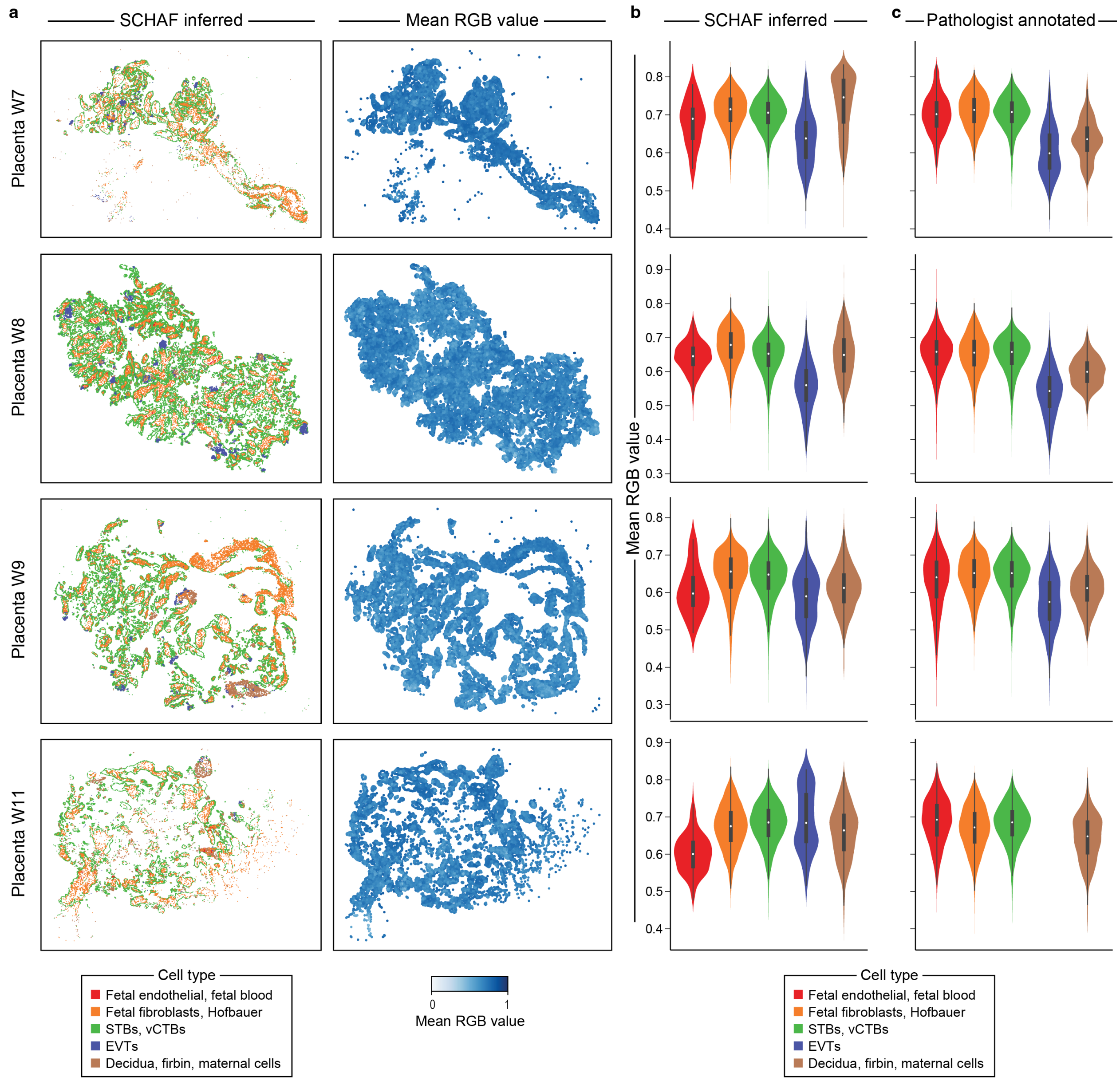

a
SCHAF inferred
Mean RGB value
b
SCHAF inferred
c
Pathologist annotated
Placenta W7
Placenta W8
Placenta W9
Placenta W11
Mean RGB value
Cell type
Fetal endothelial, fetal blood
Fetal fibroblasts, Hofbauer
STBs, vCTBs
EVTs
Decidua, firbin, maternal cells
0
1
Mean RGB value

Supp. Fig. 16

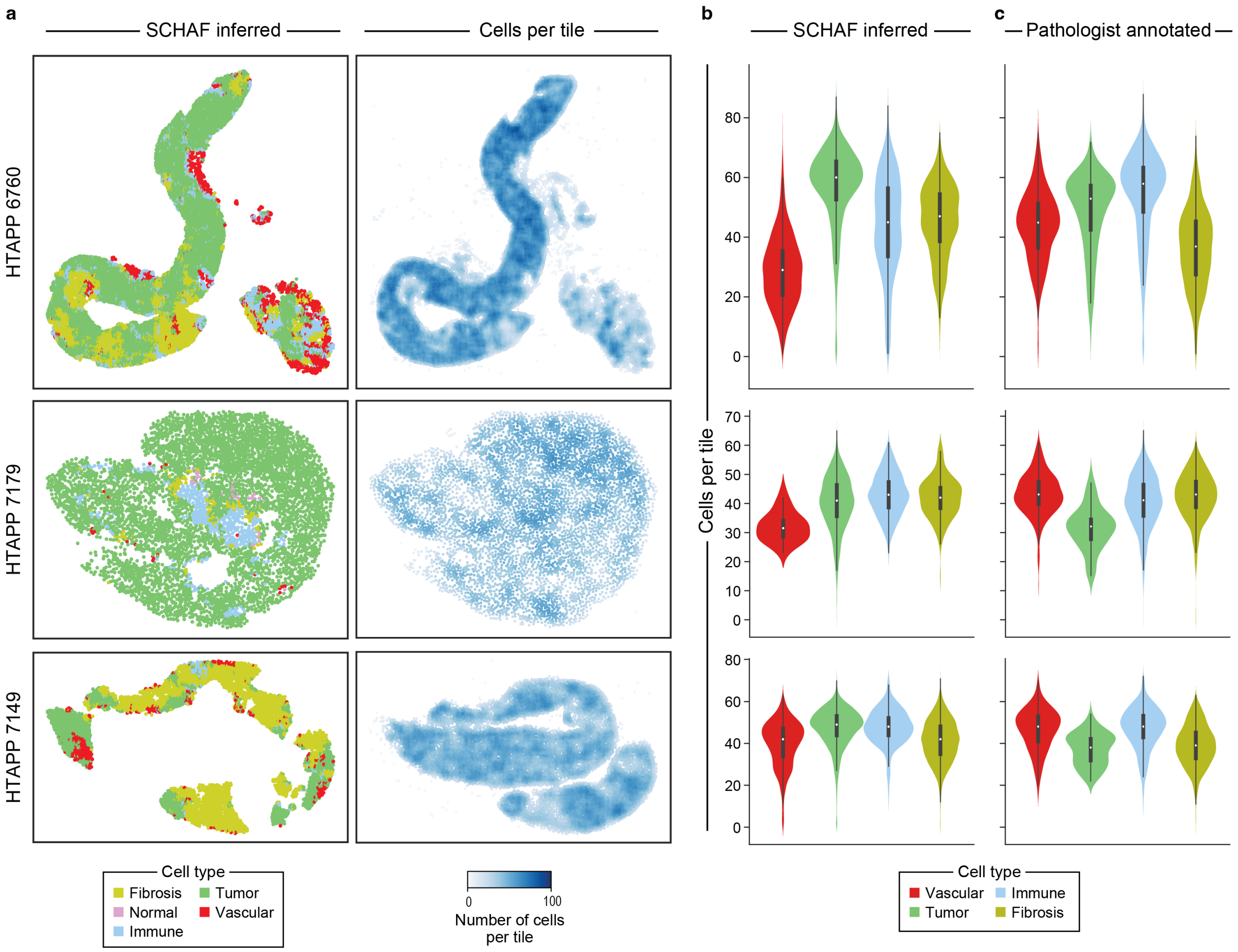
a
SCHAF inferred
Cells per tile
HTAPP 6760
HTAPP 7179
HTAPP 7149
Cell type
Fibrosis
Tumor
Normal
Vascular
Immune
0
100
Number of cells per tile
b
SCHAF inferred
c
Pathologist annotated
Cells per tile
Cell type
Vascular
Immune
Tumor
Fibrosis

Supp. Fig. 17

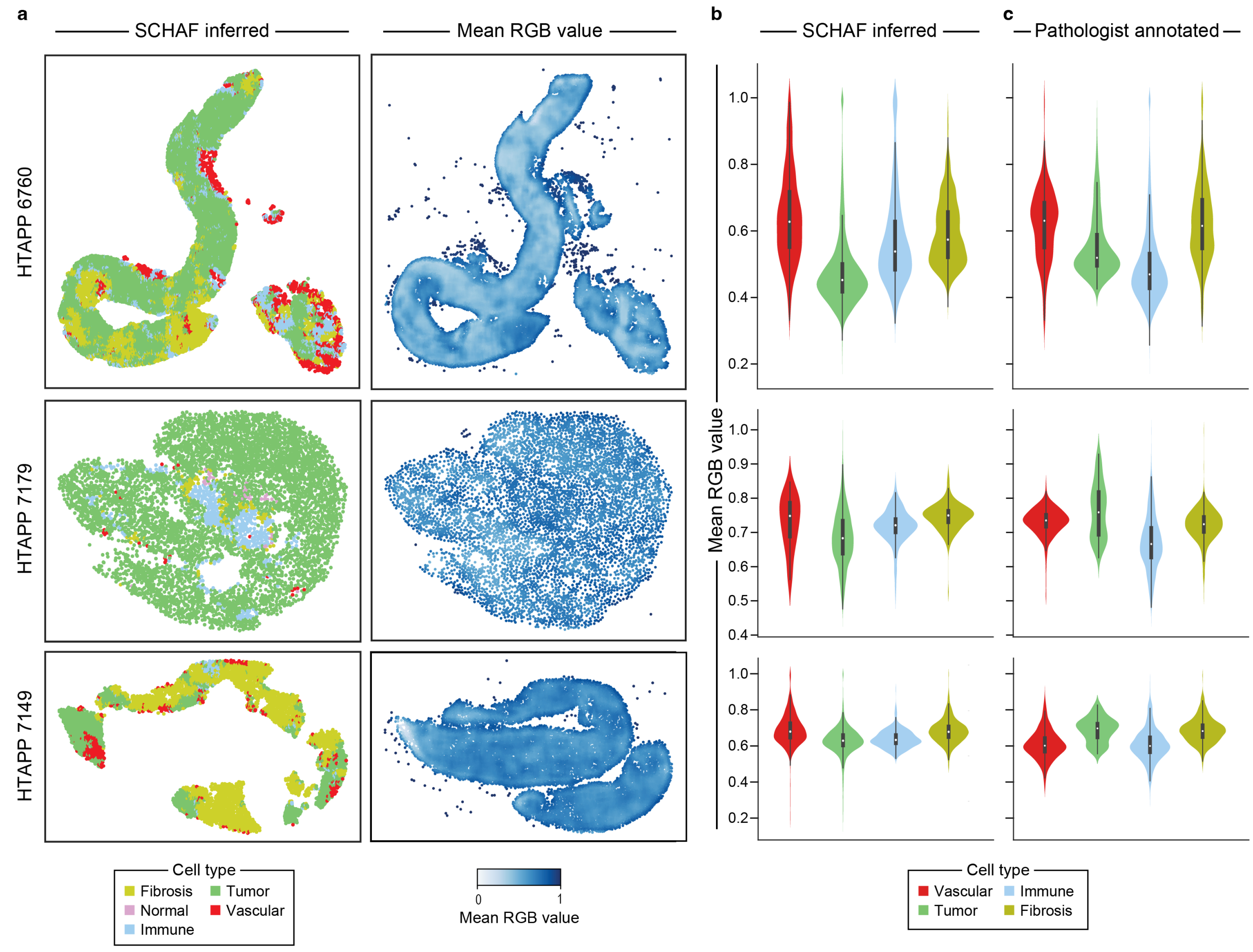
a
SCHAF inferred
Mean RGB value
HTAPP 6760
HTAPP 7179
HTAPP 7149
Cell type
Fibrosis
Tumor
Normal
Vascular
Immune
0
1
Mean RGB value
b
SCHAF inferred
c
Pathologist annotated
Mean RGB value
1.0
0.8
0.6
0.4
0.2
1.0
0.9
0.8
0.7
0.6
0.5
0.4
Cell type
Vascular
Immune
Tumor
Fibrosis

Supp. Fig. 18

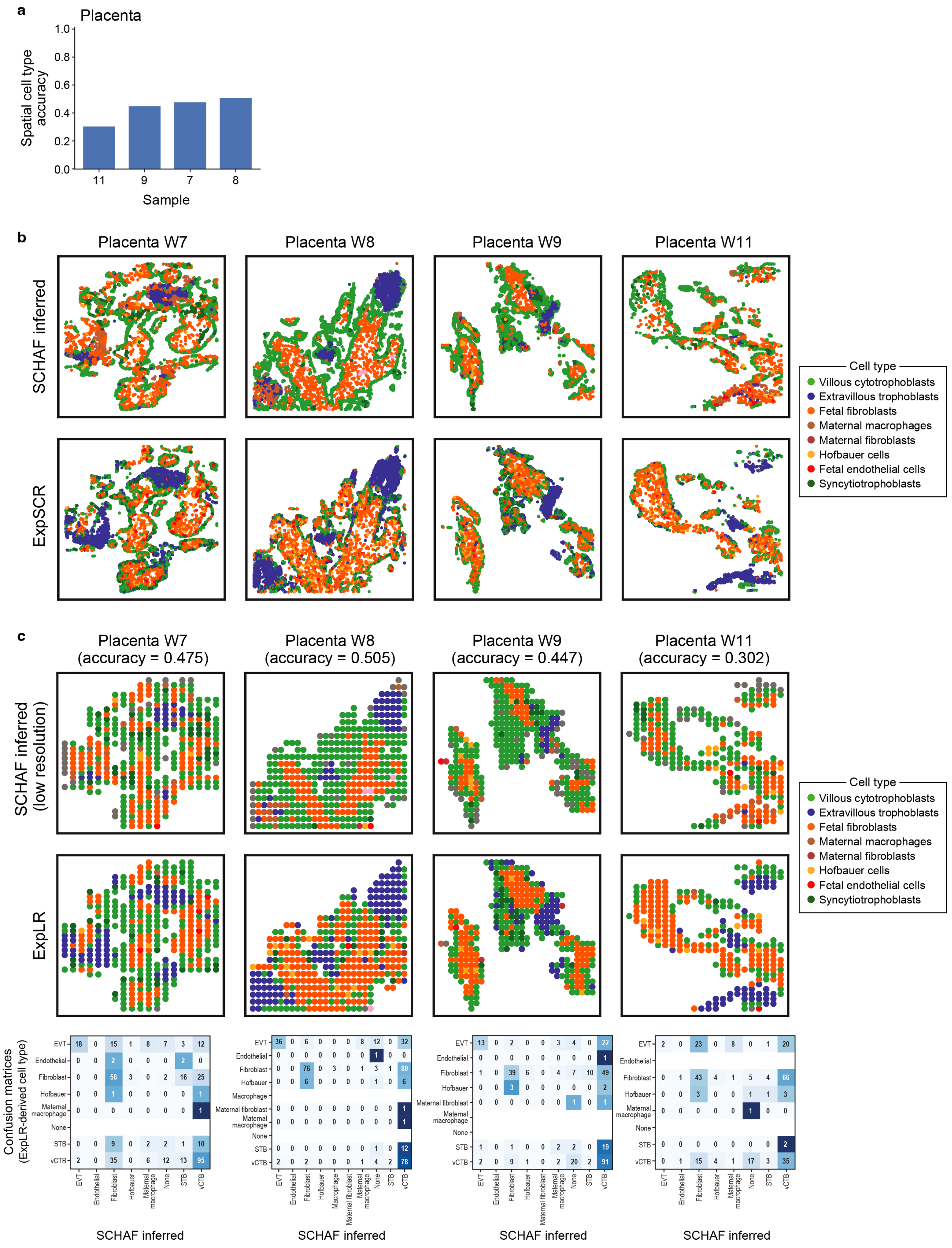
a
Placenta
Spatial cell type accuracy
1.0
0.8
0.6
0.4
0.2
0.0
11
9
7
8
Sample
b
Placenta W7
Placenta W8
Placenta W9
Placenta W11
SCHAF inferred
ExpSCR
Cell type
Villous cytotrophoblasts
Extravillous trophoblasts
Fetal fibroblasts
Maternal macrophages
Maternal fibroblasts
Hofbauer cells
Fetal endothelial cells
Syncytiotrophoblasts
c
Placenta W7 (accuracy = 0.475)
Placenta W8 (accuracy = 0.505)
Placenta W9 (accuracy = 0.447)
Placenta W11 (accuracy = 0.302)
SCHAF inferred (low resolution)
ExpLR
Cell type
Villous cytotrophoblasts
Extravillous trophoblasts
Fetal fibroblasts
Maternal macrophages
Maternal fibroblasts
Hofbauer cells
Fetal endothelial cells
Syncytiotrophoblasts
Confusion matrices (ExpLR-derived cell type)
EVT
Endothelial
Fibroblast
Hofbauer
Macrophage
Maternal fibroblast
Maternal macrophage
None
STB
vCTB
SCHAF inferred
SCHAF inferred
SCHAF inferred
SCHAF inferred

Supp. Fig. 19

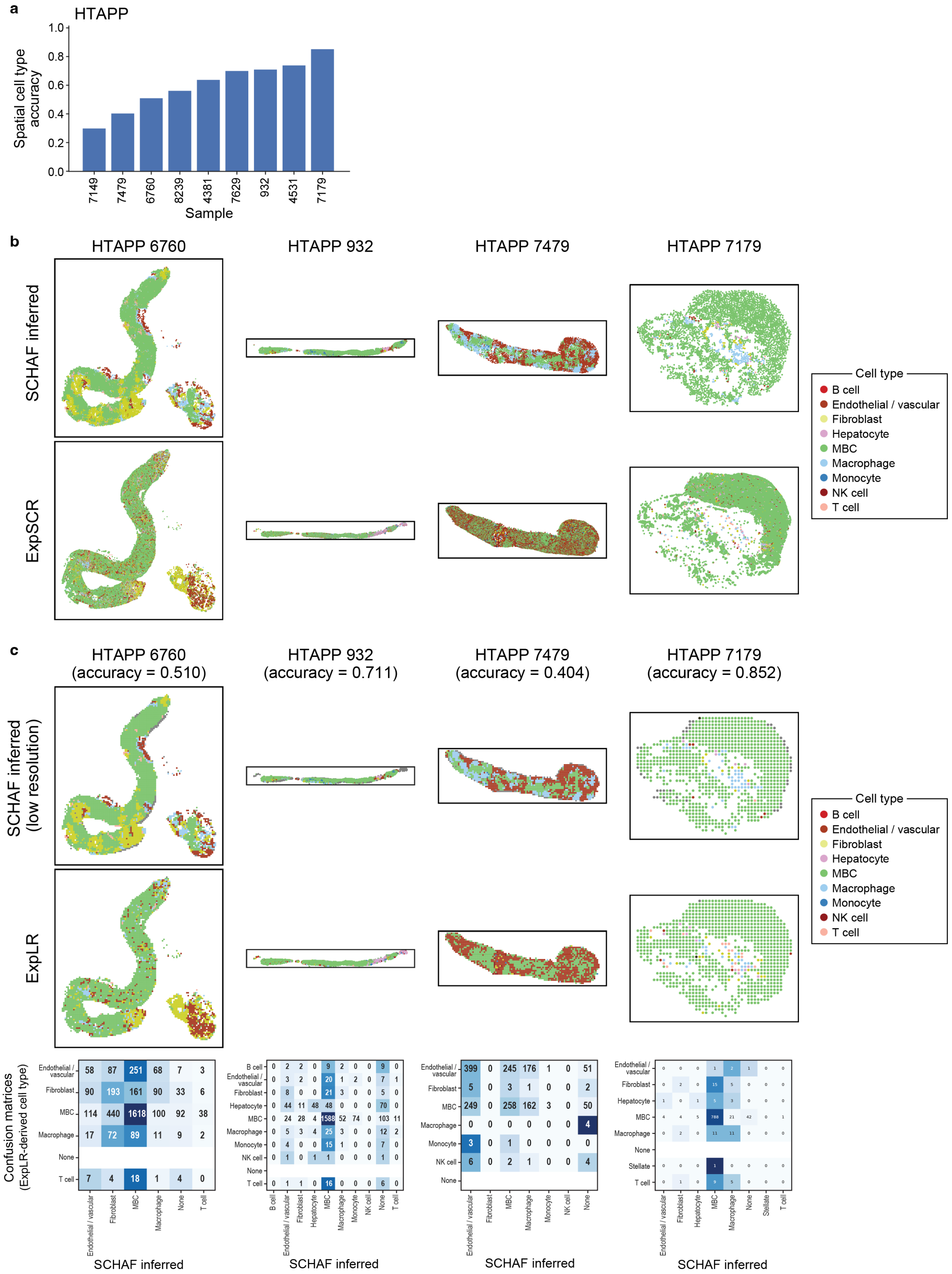
a
HTAPP
Spatial cell type accuracy
Sample
7149
7479
6760
8239
4381
7629
932
4531
7179
b
HTAPP 6760
HTAPP 932
HTAPP 7479
HTAPP 7179
SCHAF inferred
ExpSCR
Cell type
B cell
Endothelial / vascular
Fibroblast
Hepatocyte
MBC
Macrophage
Monocyte
NK cell
T cell
c
HTAPP 6760 (accuracy = 0.510)
HTAPP 932 (accuracy = 0.711)
HTAPP 7479 (accuracy = 0.404)
HTAPP 7179 (accuracy = 0.852)
SCHAF inferred (low resolution)
ExpLR
Confusion matrices (ExpLR-derived cell type)
SCHAF inferred

### Supp. Fig. 20

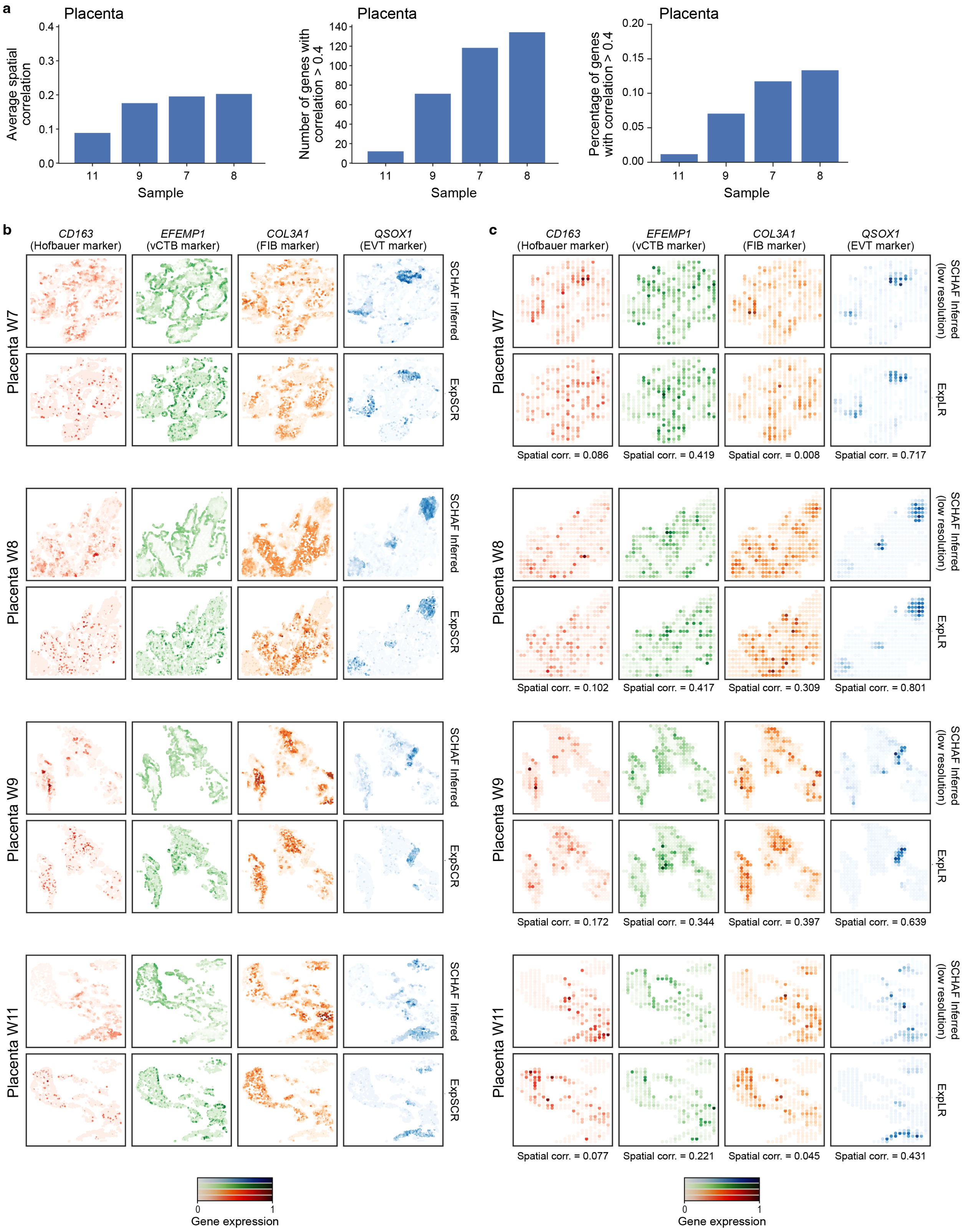
a
Placenta
Average spatial correlation
Number of genes with correlation > 0.4
Percentage of genes with correlation > 0.4
Sample
11
9
7
8
b
c
CD163 (Hofbauer marker)
EFEMP1 (vCTB marker)
COL3A1 (FIB marker)
QSOX1 (EVT marker)
Placenta W7
Placenta W8
Placenta W9
Placenta W11
SCHAF Inferred
ExpSCR
SCHAF Inferred (low resolution)
ExpLR
Spatial corr. = 0.086
Spatial corr. = 0.419
Spatial corr. = 0.008
Spatial corr. = 0.717
Spatial corr. = 0.102
Spatial corr. = 0.417
Spatial corr. = 0.309
Spatial corr. = 0.801
Spatial corr. = 0.172
Spatial corr. = 0.344
Spatial corr. = 0.397
Spatial corr. = 0.639
Spatial corr. = 0.077
Spatial corr. = 0.221
Spatial corr. = 0.045
Spatial corr. = 0.431
Gene expression
0
1

### Supp. Fig. 21

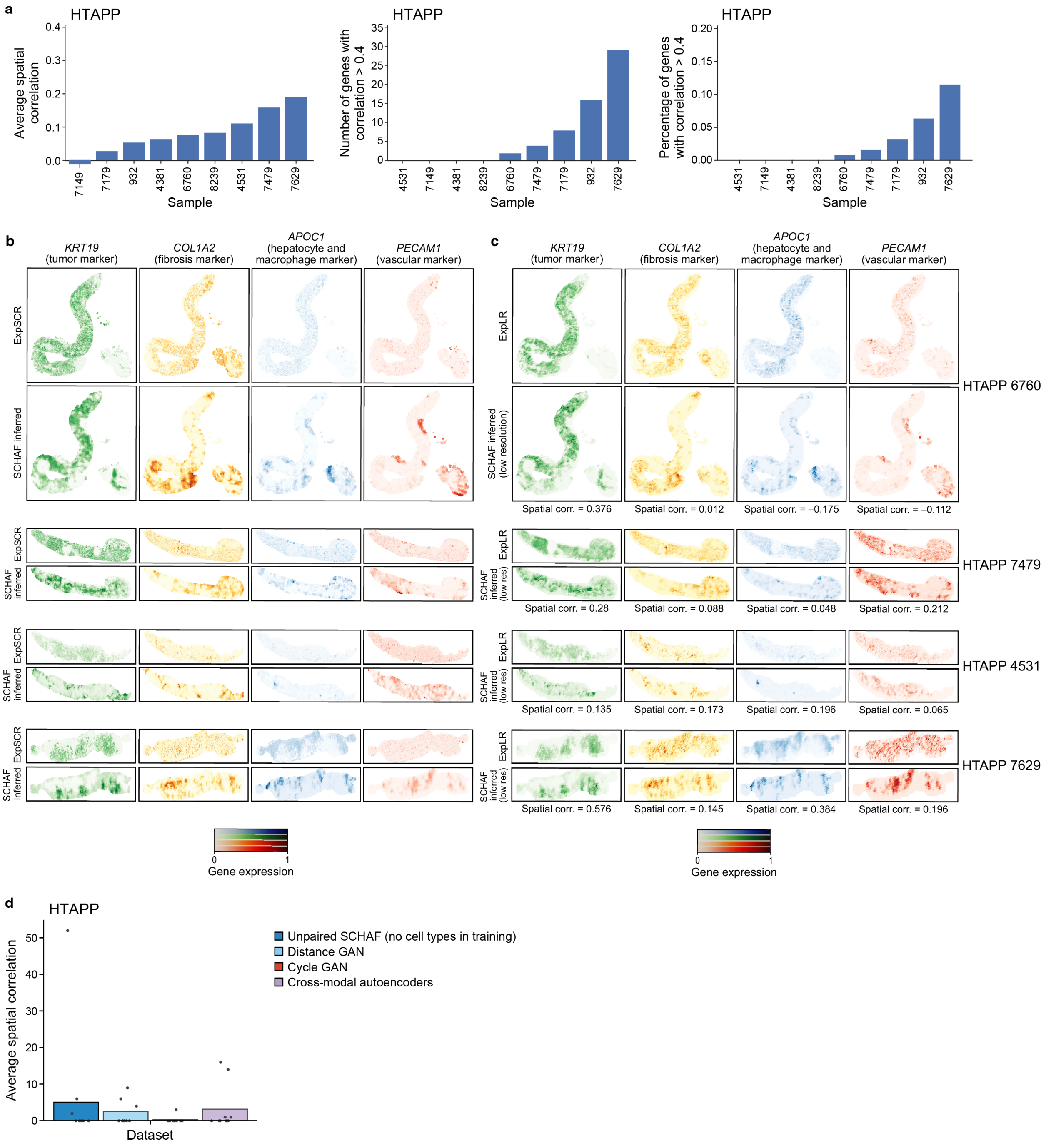
a
HTAPP
Average spatial correlation
Sample
7149
7179
932
4381
6760
8239
4531
7479
7629
Number of genes with correlation > 0.4
4531
7149
4381
8239
6760
7479
7179
932
7629
Percentage of genes with correlation > 0.4
b
KRT19 (tumor marker)
COL1A2 (fibrosis marker)
APOC1 (hepatocyte and macrophage marker)
PECAM1 (vascular marker)
ExpSCR
SCHAF inferred
c
ExpLR
SCHAF inferred (low resolution)
Spatial corr. = 0.376
Spatial corr. = 0.012
Spatial corr. = –0.175
Spatial corr. = –0.112
HTAPP 6760
SCHAF inferred (low res)
Spatial corr. = 0.28
Spatial corr. = 0.088
Spatial corr. = 0.048
Spatial corr. = 0.212
HTAPP 7479
Spatial corr. = 0.135
Spatial corr. = 0.173
Spatial corr. = 0.196
Spatial corr. = 0.065
HTAPP 4531
Spatial corr. = 0.576
Spatial corr. = 0.145
Spatial corr. = 0.384
Spatial corr. = 0.196
HTAPP 7629
Gene expression
d
HTAPP
Average spatial correlation
Dataset
Unpaired SCHAF (no cell types in training)
Distance GAN
Cycle GAN
Cross-modal autoencoders
